# pHLARE: A Genetically Encoded Ratiometric Lysosome pH Biosensor

**DOI:** 10.1101/2020.06.03.132720

**Authors:** Bradley A. Webb, Jessica Cook, Torsten Wittmann, Diane L. Barber

**Author notes:** To whom correspondence should be addressed Diane L. Barber, Box 0512, 513 Parnassus Ave., San Francisco, CA 94143.

## Abstract

Many lysosome functions are determined by a lumenal pH of ~5.0, including the activity of resident acid-activated hydrolases. Lysosome pH (pHlys) is increased in neurodegenerative disorders and predicted to be decreased in cancers, making it a potential target for therapeutics to limit the progression of these diseases. Accurately measuring pHlys, however, is limited by currently used dyes that accumulate in multiple intracellular compartments and cannot be propagated in clonal cells for longitudinal studies or *in vivo* determinations. To resolve this limitation, we developed a genetically encoded ratiometric pHlys biosensor, pHLARE (<u>pH L</u>ysosomal <u>A</u>ctivity <u>RE</u>porter), which localizes predominantly in lysosomes, has a dynamic range of pH 4.0 to 6.5, and can be stably expressed in cells. Using pHLARE we show decreased pHlys with inhibiting activity of the mammalian target of rapamycin complex 1 (mTORC1), in breast and pancreatic cancer cells compared with tissue-matched untransformed cells, and with the activated oncogene H-RasV12. pHLARE is a new tool to accurately measure pHlys, for improved understanding of lysosome dynamics that could be a promising therapeutic target.

**Summary Statement:** Most lysosome functions require a low lumenal pH, which is dysregulated in many human diseases. We report a new genetically biosensor to accurately measure lysosome pH dynamics, which we use to show decreased lysosome pH in cancer cell lines.

## Introduction

Lysosomes function as catabolic hubs independently as well as downstream of autophagy and nutrient sensing by mammalian target of rapamycin complex 1 (mTORC1). Additionally, lysosomes contribute to trafficking of intracellular vesicles, plasma membrane repair, pathogen degradation, resistance to chemotherapies, and a broad range of homeostatic responses to environmental cues (Xu and Ren, 2015; Perera and Zoncu, 2016). The lumenal pH of lysosomes (pHlys) is a major determinant of many lysosome functions, including catabolism by lumenal acid-activated hydrolyases (Mindell, 2012), fusion with endosomes and cargo sorting (Marshansky and Futai, 2008; Scott and Gruenberg, 2011), and roles in Ca^2+^ homeostasis (Lee et al., 2015). Although pHlys in normal cells is thought to be tightly regulated at ~ 5.0, it is increasingly recognized to be dysregulated in diseases. Increased pHlys occurs in neurodegenerative disorders (Majumdar et al., 2007; Wolfe et al., 2013; Lee et al., 2015) and diabetic nephropathy (Liu et al., 2015), which attenuates activity of lumenal acid-activated hydrolases with a predicted consequence of decreased protein degradation leading to protein aggregation. Increased pHlys is also a determinant in some pathologies of lysosomal storage diseases (Colacurcio and Nixon, 2016) and in osteopetrosis (Kornack et al., 2001). In contrast, decreased pHlys may occur in cancers compared with untransformed cells, based on changes in autophagosome activity (Kenific and Debnath, 2015), roles in multidrug resistance (Daniel et al, 2013; Zhitomirsky and Assaraf, 2016), and reversed pHlys and cytosolic pH dynamics (Liu et al., 2018), with the latter confirmed to be higher in most cancers (Webb et al., 2011; White et al., 2017b). A lower pHlys in cancer would be consistent with cancer cells having increased lysosomal catabolism of macromolecules (Perera and Zoncu, 2016), which likely contributes to metabolic reprograming for increasing biomass to fuel rapid proliferation. Accordingly, targeting mechanisms of dysregulated pHlys dynamics is proposed as a therapeutic approach to limit some disease pathologies (Wolfe et al., 2013; Appelqvist et al. 2013; Piao and Amaravadi, 2016; Fraldi et al., 2016).

Accurately determining pHlys, however, is currently limited by the lack of selective and effective probes, particularly for time-lapse or *in vivo* studies. This is in contrast to a number of highly reliable reagents for accurately measuring cytosolic pH, including fluorescent dyes and genetically encoded biosensors (Grillo-Hill et al., 2014). The most commonly used reagents for measuring pHlys, fluorescein dextran, LysoTracker, and LysoSensor, have a number of caveats with the most significant being that their signals are not exclusively in lysosomes because these reagents are incorporated by phagocytosis and accumulate in multiple types of intracellular vesicles. Also, dye loading into cells requires hours; often 6-8 h for uptake and another 24 h for transport through endo-lysosomal trafficking (Nilsson et al., 2010; Wolfe et al, 2013; Johnson et al., 2016). A recent comparison indicates additional dye-specific limitations (Wolfe et al., 2013). Fluorescein dextran has a low fluorescent signal in acidic compartments and it photobleaches rapidly. LysoTracker is not ratiometric and hence allows only qualitative determinations. Lysosensor, a dextran-conjugated ratiometric dye that allows quantitative measurements and has the best dynamic range of dyes used to measure pHlys but is not suitable for measurements of more than 5 min because it induces an increase in pHlys. Additionally, because LysoSensor and LysoTracker are substrates for P-glycoprotein (Zhitomirsky et al., 2018), their use in determining the role of pHlys in resistance to cancer therapies is problematic. Oregon green 488–dextran is reported as an improved probe to measure pHlys, but for quantitative ratiometric imaging cells are co-loaded with the pH-insensitive tetramethylrhodamine-dextran and loading is achieved over 24 h by phagocytosis and trafficking to lysosomes (Johnson, et al., 2016). Nanoprobes that are pH sensitive have also been used for determining organelle pH, but lysosome targeting remains problematic, and a recently developed two-photon fluorophore conjugated to a lysosome targeting morpholine (Wang et al., 2018) achieves specificity, but cellular uptake is similar to dyes and nanoprobes, and as reported is not quantitative.

For an improved method to accurately and selectively quantify pHlys in real-time over long time periods we developed a genetically encoded and ratiometric pHlys biosensor designated pHLARE (<u>pH L</u>ysosome <u>A</u>ctivity <u>RE</u>porter). Here, we describe the pHLARE design of the lysosomal associated membrane protein 1 (LAMP1) tagged at lumenal and cytosolic domains with distinct fluorophores, validate that pHLARE is predominantly expressed in lysosomes and has strong sensitivity with a broad dynamic range, and use pHLARE to show changes in pHlys with known pharmacological regulators, including a decrease with inhibiting mTORC1 activity. We also confirm a lower pHlys in clonal breast and pancreatic cancer cells compared with tissue-matched untransformed cells and in mammary epithelial cells expressing oncogenic H-RasV12.

## Results

### pHLARE design and validation

To selectively quantify pHlys we generated pHLARE, a genetically encoded and ratiometric biosensor that can be stably expressed in cells. pHLARE encodes rat LAMP1, a lysosome transmembrane protein, tagged at the lumenal amino-terminus with superfolder GFP (sfGFP) (Fig. 1A), a GFP variant having a pKa ~ 5.0. We tested tagging the lumenal domain of LAMP1 with several fluorophores and found that sfGFP is optimal for dynamic range and limited photobleaching. EGFP has a pKa too high for the low lysosome lumenal pH. mT-Sapphire and mCherry each have an appropriate pKa ~ 4.5, but mT-Sapphire had substantial photobleaching over five min imaging times and mCherry aggregated in the lysosome lumen and responded poorly for calibrating fluorescent signal to pH. In contrast, mCherry did not aggregate in the cytoplasm, hence we tagged the cytoplasmic carboxyl-terminus of LAMP1 with mCherry (Fig. 1A). The low pKa of mCherry makes it insensitive to changes in the cytosolic pH range and with sfGFP allows ratiometric analysis normalized for abundance of the biosensor. When expressed in human retinal pigment epithelial (RPE) cells, rat pHLARE localized predominantly in lysosomes, as indicated by co-labeling with endogenous human LAMP1 (Fig. 1B) and also with with SiR-lysosome, a membrane-permeant far-red-tagged pepstatin A that binds cathepsin D (Fig. 1C). By incubating cells in buffers at known pH values and containing the protonophore nigericin that allows intracellular membrane-bound compartments to equilibrate with extracellular pH, pHLARE had a ratiometric dynamic range between pH 4.0 and 6.5 (Fig. 1D) and no apparent degradation, as indicated by immunoblotting cell lysates with antibodies to RFP (Fig. 1E)

**Figure 1.**
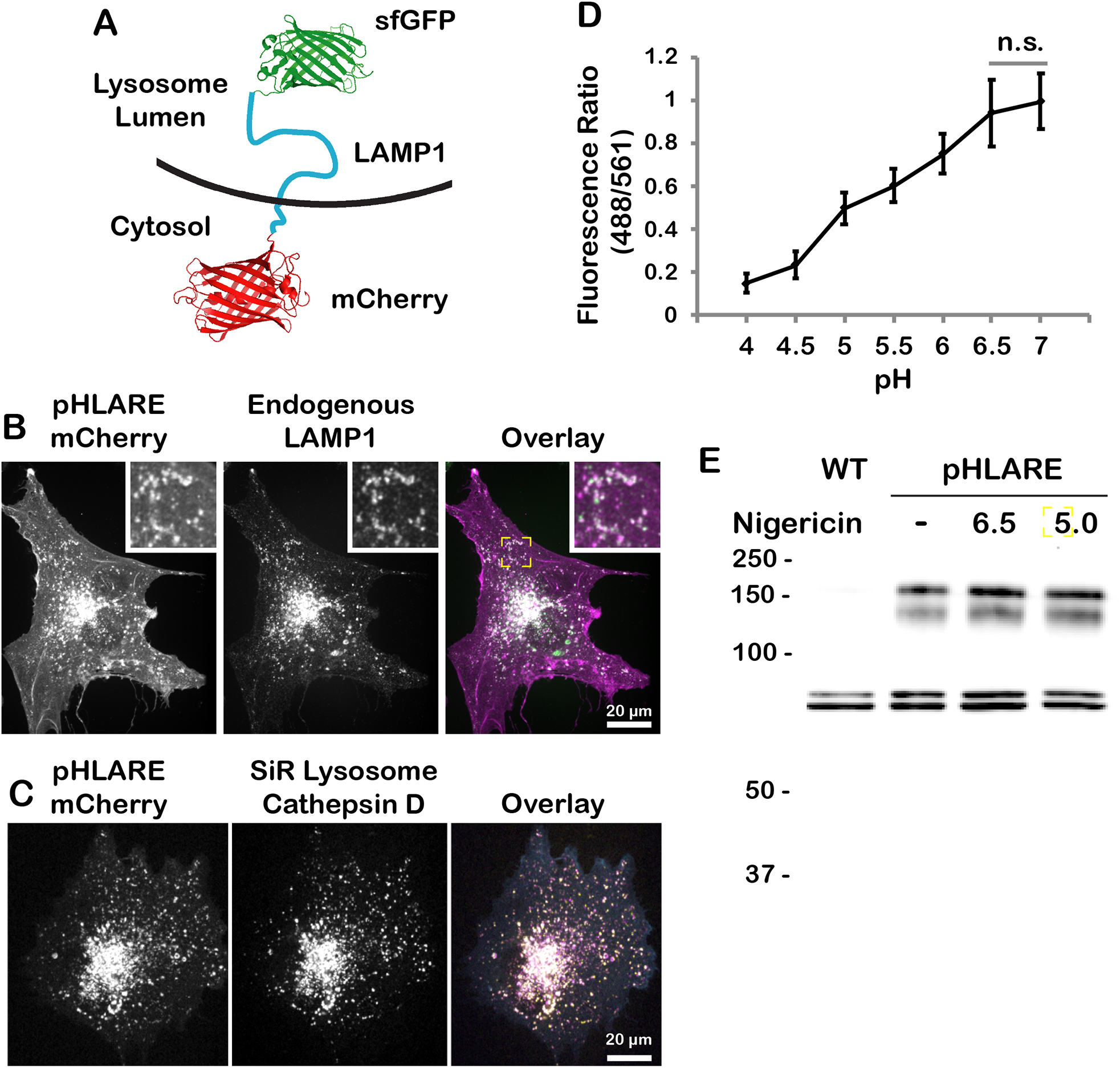
pHLARE localizes to lysosomes. **(A)** Schematic of pHLARE with rat LAMP1 tagged at the lumenal amino-terminus with sfGFP and at the cytoplasmic carboxyl-terminus with mCherry. (**B)** pHLARE stably expressed in human RPE cells, visualized with anti-mCherry antibodies, co-localizes with endogenous LAMP1, visualized with anti-human LAMP1 antibodies. **(C)** Images of live RPE cells stably expressing pHLARE and stained with SiR-lysosome, a far-red pepstatin A that binds cathepsin D. **(D)** Fluorescence ratios of pHLARE in RPE cells in nigericin-containing buffers between pH 4.0 to 7.0. Data are means ± SD of 15 cells from three separate cell preparations. Statistical analysis by Tukey-Kramer HSD indicates significant differences at all pH values except between pH 6.5 and 7.0. **(E)** Representative RFP immunoblot of two preparations of lysates from RPE cells, wild type (WT) and stably expressing pHLARE, and untreated or treated with nigericin buffer at the indicated pH values for 5 min.

Calibrating fluorescence ratios of pHLARE to pH with nigericin (Fig. 2A, B) is similar to calibrating measurements of cytosolic pH (Grillo-Hill et al., 2014). However, because lysosomes have differences in size, shape, aggregation and localization, calibrating pHLARE fluorescence is more complex than calibrating fluorescent probes for cytosolic pH. To manage this complexity, we developed an image analysis algorithm to objectively and reproducibly define lysosome objects in the pHlys-insensitive mCherry channel (Fig. 2C). The average fluorescence intensities of these objects and corresponding local background areas in the two pHLARE fluorescence channels were then used to calculate fluorescence ratios and pH values for each lysosome object based on the nigericin calibration. Although we typically measured nearly a hundred lysosome objects per cell, lysosomes were often close to one another and many lysosome objects would contain more than one lysosome. While it may be possible to estimate the number of individual lysosomes based on the area of a lysosome cluster, we did not attempt this. In addition, because pHLARE fluorescence channels are acquired sequentially, a very small number of lysosomes that move substantially between the two image acquisitions will yield erroneous measurements. To minimize the impact of these outliers, we used the median value of all individual lysosome object measurements per cell to report the average pHlys per cell. Because we were concerned that pHlys equilibration would be incomplete in nigericin due to potentially high buffering capacity, we tested two different calibration methods either using the local background around lysosomes to measure pHLARE fluorescence ratios in nigericin or the lysosome signal itself. We validated this analysis by comparing pHlys measurements in six RPE cells (Fig. 2D). The two calibration methods gave median pH values that were not significantly different (local background calibration pHlys = 4.95 ± 0.22 (mean and SD); lysosome signal calibration pHlys = 5.06 ± 0.21); hence, going forward we used the local background calibration. Fluorescence at the plasma membrane, which is excluded from the pHlys analysis, could be due to lysosome exocytosis (Rodriguez et al., 1997; Jaiswal et al., 2002) or direct trafficking of LAMP1 to the plasma membrane (Rohrer et al., 1996).

**Figure 2.**
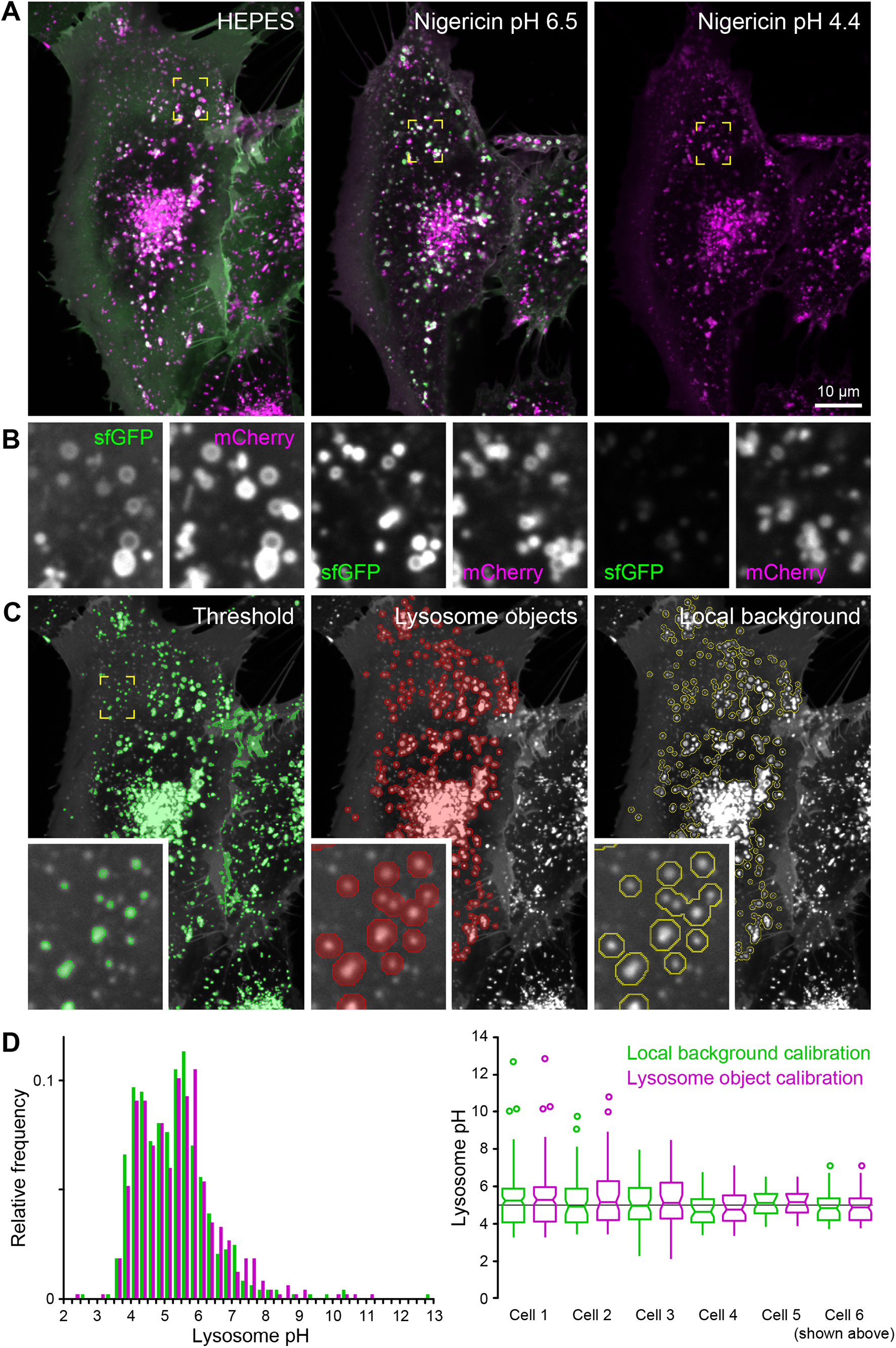
pH calibration of pHLARE. (**A)** pHLARE-expressing RPE cell in buffer and in nigericin buffers of two different calibration pH values showing sfGFP (green) and mCherry (magenta) fluorescence. The three images are scaled identically showing the dramatic decrease of sfGFP fluorescence at low pH. **(B)** Higher magnification of individual pHLARE fluorescence channels from the regions indicated in (A). Note that pHLARE lysosome membrane localization is only clearly resolved in larger lysosomes. **(C)** Example image segmentation showing the raw intensity threshold, lysosome and local background objects. **(D)** Comparison of lysosome pH measurements in six different pHLARE-expressing RPE cells obtained with the two different calibration methods. The histogram on the left shows the distribution of pH values in all lysosome objects from these six cells (n = 485 lysosome objects). The box-and-whisker plot on the right shows the median pHlys for each of these six cells separately. Cell number 6 is the one shown in (A-C), and this data set is included in the supplemental information.

Using pHLARE, we determined steady-state pHlys in untransformed cells from different species and tissue origins. Human RPE and canine kidney MDCK epithelial cells (Fig. 3A) as well as human MCF10A mammary epithelial cells (Fig. 5B, G) and HPDE pancreatic ductal epithelial cells (Fig. 5E) had a mean cell pHlys between 5.0 to 5.4 that is similar to the pHlys of 5.1 to 5.2 in HeLa cells determined using cresyl violet (Ostrowski et al., 2016) or Oregon green-dextran (Johnson et al., 2016) but substantially higher than the ~4.3 in keratinocytes determined using FITC-conjugated dextran and flow cytometry (Nilsson et al., 2010). For MCF10A cells there was no difference in pHlys when pHLARE was transiently or stably expressed cells (Fig. S3). Mouse NIH 3T3 fibroblasts had a higher mean cell pHlys of ~ 6.0 compared with RPE and MDCK epithelial cells (Fig. 5A) and is also higher than the ~ 4.9 in mouse Swiss 3T3 fibroblasts determined by fluorescence-conjugated dextran (Lin et al., 2003) and in embryonic fibroblasts determined using LysoSensor (Wolfe et al., 2013). Our data indicate that the steady-state pHlys of most untransformed cells we tested is ~5.0, except for a higher pHlys in NIH 3T3 fibroblasts, the latter warranting further investigation to determine a possible functional significance.

**Figure 3.**
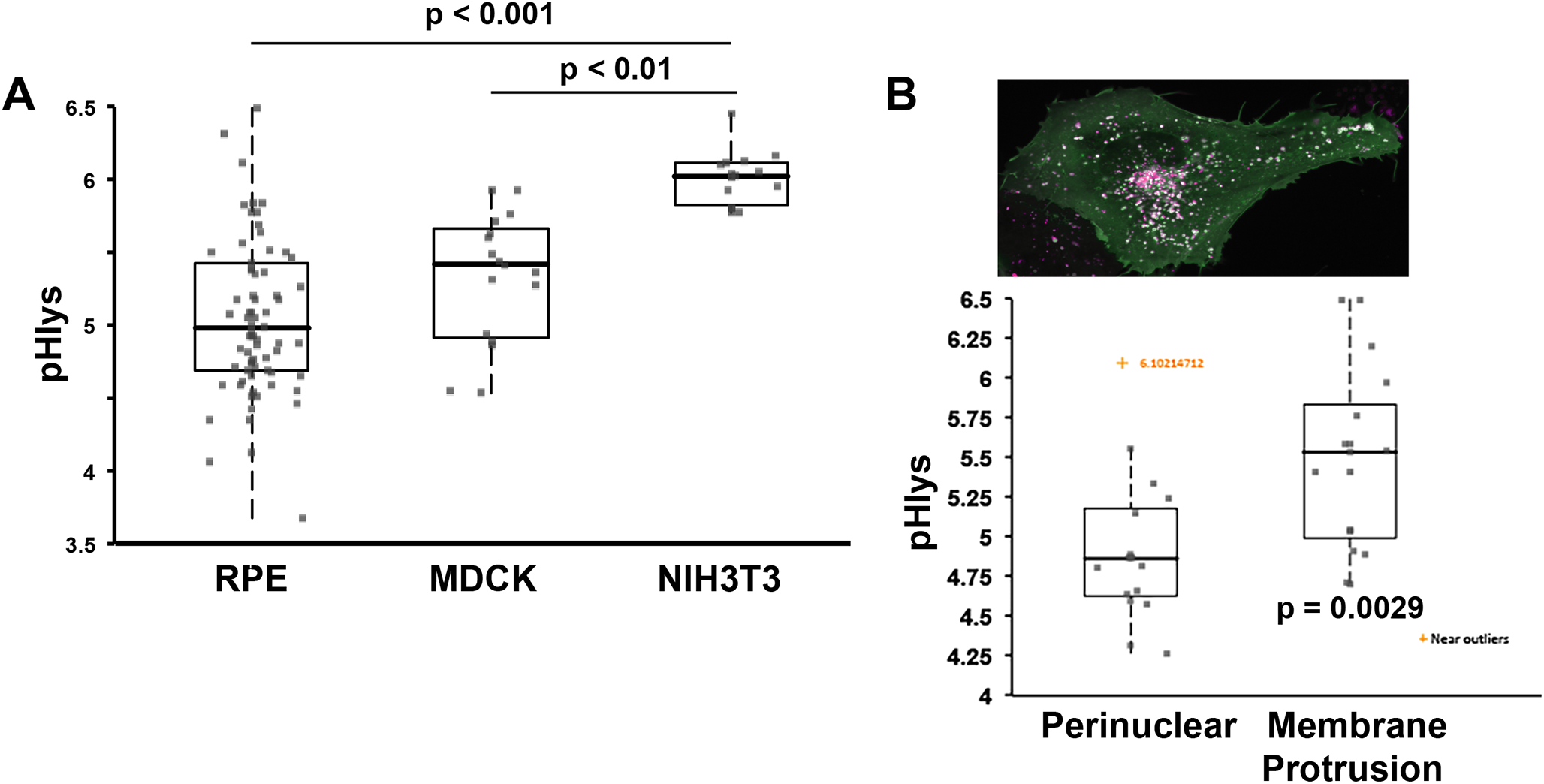
Cell-specific pHlys. **(A)** Average steady-state pHlys in RPE cells stably expressing pHLARE (n = 74 cells) and in MDCK and NIH-3T3 cells transiently expressing pHLARE (n = 17 and n = 15 cells, respectively) and maintained in growth medium. Box plots show median, first and third quartile, with whiskers extending to observations within 1.5 times the interquartile range, and all individual data points include data for individual cells (each closed circle) obtained from five separate cell preparations for RPE cells, and three separate cell preparations for MDCK and NIH-3T3 cells. Statistical analysis by Tukey-Kramer HSD test. **(B)** Steady-state pHlys of lysosomes within 3 ∝m of the nuclear membrane (perinuclear) and 3 ∝m of the distal margin of membrane protrusions in RPE cells. Data are expressed as described in A and obtained from six separate cell preparations.

The values described above are averages for all lysosomes in the indicated cell types; however, recent findings suggest that pHlys within each cell may be heterogeneous (Johnson et al., 2016), which we also see (Fig. 2D). In many cell types lysosomes have a predominantly perinuclear localization, but smaller populations are scattered within the cytosol and are also peripheral near the plasma membrane. Perinuclear lysosomes are suggested to have a lower lumenal pH than peripheral lysosomes (Johnson et al., 2016), although this was not quantitatively determined but rather predicted from qualitative differences in the fluorescence of Alexa Fluor-conjugated dextran. Additionally, spatially heterogeneous pHlys may depend on cell type and conditions. For human osteocarcoma cells, lowering extracellular pH of the medium causes a redistribution of lysosomes from predominantly perinuclear to predominantly peripheral (Walton et al., 2018). We found that in RPE cells with clear membrane protrusions, lysosomes within 3 ∝mof the distal membrane have an average pHlys of 5.48 ± 0.12 (SD) that is significantly higher than the average of 4.91 ± 0.12 of lysosomes within 3 ∝m of the nuclear membrane (Fig. 3B), indicating spatially distinct lysosomes have different steady-state lumenal pH values.

### pHLARE measures dynamic changes in pHlys

We used pHLARE to confirm increased pHlys when RPE cells are incubated with NH_4_Cl (5 mM) for 18 h (Fig. S4), which increases the pH of membrane-bound compartments as NH_3_ entering compartments complex with H_+_ (White et al., 2017a). We also confirmed increased pHlys in RPE cells treated for 60 min with bafilomycin (100 nM), a V-ATPase inhibitor, and with chloroquine (100 μM), which neutralizes acidic intracellular compartments (Fig. 4A, B). In control RPE cells, lysosome size is within the reported range of 100-1000 nm (Appelqvist et al., 2013; Bandyopadhyay et al., 2014; Xu and Ren, 2015), with the majority of lysosomes having a mean size of 784 ± 133 nm (SD). With chloroquine but not with bafilomycin, however, lysosomes are larger with a mean size of 934 ± 244 nm and fewer lysosomes smaller than 600 nm and more lysosomes larger than 1000 nm (Fig. 4A, C). Lysosome size can increase as a result of accumulated undigested material. However, the increased pHlys with bafilomycin and chloroquine were similar; hence, both presumably decreased catabolism but not an increase in lysosome size, which suggests a different mechanism for increased size with chloroquine. Possible mechanisms include enlargement from chloroquine accumulating in the lumen or activation of the transcription factor EB (TFEB) pathway that increases lysosome biogenesis in response to lysosomal storage dysfunction (Sardiello et al., 2009). Although not quantified, there are visually more lysosomes in cells treated with chloroquine compared with bafilomycin or untreated RPE cells (Fig. 4A).

**Figure 4.**
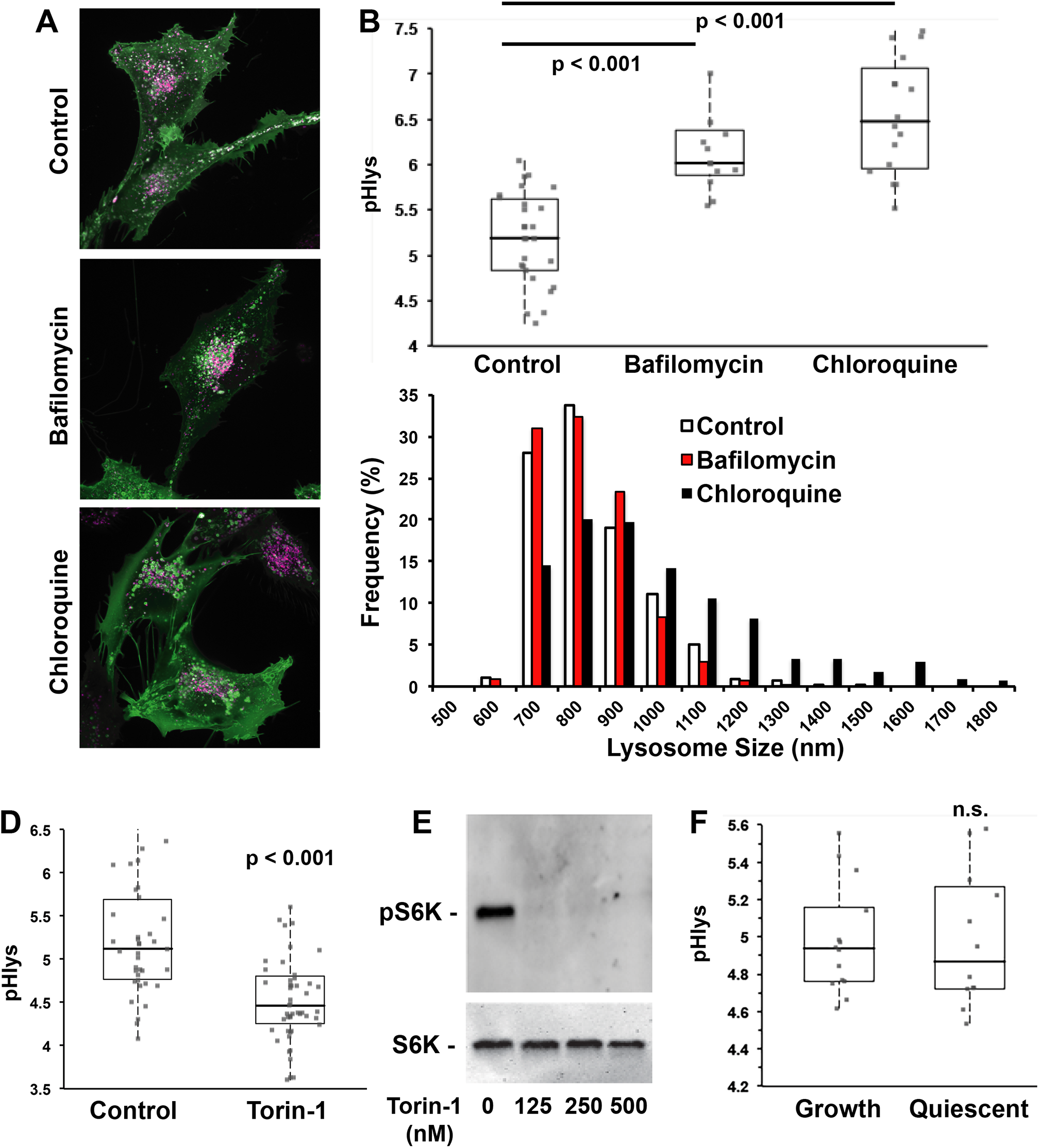
pHLARE measures dynamic changes in pHlys. **(A, B)** RPE cells stably expressing pHLARE were untreated (control, n = 27 cells) or treated with 100 nM of the V-ATPase inhibitor Bafilomycin A1 (n = 15 cells) or 100 ∝M chloroquine (n = 15 cells) for 2 h before acquiring images, which were used to calculate pHlys. **(C)** Lysosome size with indicated conditions determined by measuring more than 1,000 lysosomes in four cell preparations for controls and three cell preparations with bafilomycin and chloroquine treatment. **(D)** pHlys of RPE cells stably expressing pHLARE untreated (control, n = 39 cells) or treated with 250 nM Torin-1 (n = 50 cells) for 2 h before acquiring images. **(E)** Representative immunoblot of lysates from two separate RPE cell preparations untreated or treated with the indicated concentrations of Torin-1 and probed with antibodies to total and phosphorylated S6K. **(F)** pHlys of RPE cells stably expressing pHLARE maintained for 24 h before imaging in growth medium (10% FBS; n = 18 cells) or quiescent medium (0.2% FBS; n = 15 cells). Box plots show median, first and third quartile, with whiskers extending to observations within 1.5 times the interquartile range, and all individual data points. Statistical analysis by Tukey-Kramer HSD test, with data obtained from three separate cell preparations in B, C, and F, and four separate cell preparations in D.

Lysosome functions in nutrient sensing and responses are in part determined by mTORC1, which when active is associated with the lysosomal membrane. Inside (lysosome) to outside (cytoplasm) signaling regulates mTOCR1 activity (Settembre et al., 2013; Wang et al., 2015; Rebsamen et al., 2015). Whether mTORC1 activity regulates pHlys, however, has received limited attention. Pharmacologically inhibiting mTOCR1 increases autophagy and decreases pHlys; however, the latter was determined qualitatively by increased LysoTracker and LysoSensor staining but not quantitatively by calibrating the fluorescence signal to pH (Zhou, 2013). We used RPE cells stably expressing pHLARE and quantitative measurements to show that Torin-1, an ATP-competitive inhibitor of mTORC1 activity (Thoreen et al., 2009), decreased pHlys from 5.22 ± 0.10 in controls to 4.63 ± 0.09 (mean ± s.e.m., 4 cell preparations) (Fig. 4D). We confirmed that Torin-1 decreases phosphorylation of ribosomal protein S6 kinase beta-1 (S6K1) (Fig. 4E), an mTORC1 substrate, and treated RPE cells with 250 nM Torin-1 for 2 h as previously described (Thoreen et al., 2012). Decreased pHlys with inhibiting mTORC1 activity is consistent with the established role of active mTORC1 in stimulating biosynthetic pathways and inhibiting cellular catabolism (Saxton and Sabatini, 2017). Accordingly, our findings suggest that active mTORC1 might increase pHlys to limit catabolism. However, pHlys was not different in quiescent RPE cells maintained for 24 h in medium containing 0.2% FBS compared with cells maintained in growth medium containing 10% FBS (Fig. 4F). Previous findings indicate that acute (2-6 h) serum-deprivation markedly changes lysosome morphology from circular to tubular and increases pHlys, although the latter was determined qualitatively but not quantitatively using the acid-dependent dye Lysotracker, and by 24 h lysosome morphology and lumenal pH was restored to steady-state conditions by a feedback loop involving increased mTOR activity (Yu et al., 2010). Our findings suggest that pHlys dynamics over longer periods may be more responsive signals related to nutrient deprivation and not general growth conditions.

### Decreased pHlys in breast, pancreatic, and glioblastoma cancer cells

Cancer progression and metastasis are associated with striking changes in lysosomes, including their volume, composition, cellular distribution, and lumenal enzyme activities (Appelqvist et al., 2013; Perera and Bardeesy, 2015; Hamalisto and Jaattela, 2016). Despite many of these properties being determined by pHlys, there are limited data on dysregulated pHlys in cancer compared with tissue-matched untransformed cells (Nilsson et al., 2010). We used pHLARE to measure steady-state pHlys in clonal human breast and pancreatic cancer cells. The pHlys of MDA-MB-157 and MDA-MB-453 basal, triple negative, and invasive breast cancer cells (4.56 ± 0.11, and 4.31 ± 0.28, respectively, mean ± s.e.m.) was significantly lower than in immortalized but not transformed MCF10A mammary epithelial (5.36 ± 0.09) (Fig. 5A, B). In contrast, the pHlys of MCF7 lumenal, estrogen receptor- and progesterone receptor-positive, and benign breast cancer cells (5.34 ± 0.12) was not different than MCF10A cells (Fig. A, B), which was also reported (Altan et. al., 1999). In MDA-MB-157 but not MCF7 cells there were more peripherally localized lysosomes compared with MCF10A cells (Fig. 5C), determined by mCherry fluorescence within 3 μm of the plasma membrane, which is consistent with the increased invasiveness of basal breast cancer cells (Neve et. al., 2006).

**Figure 5.**
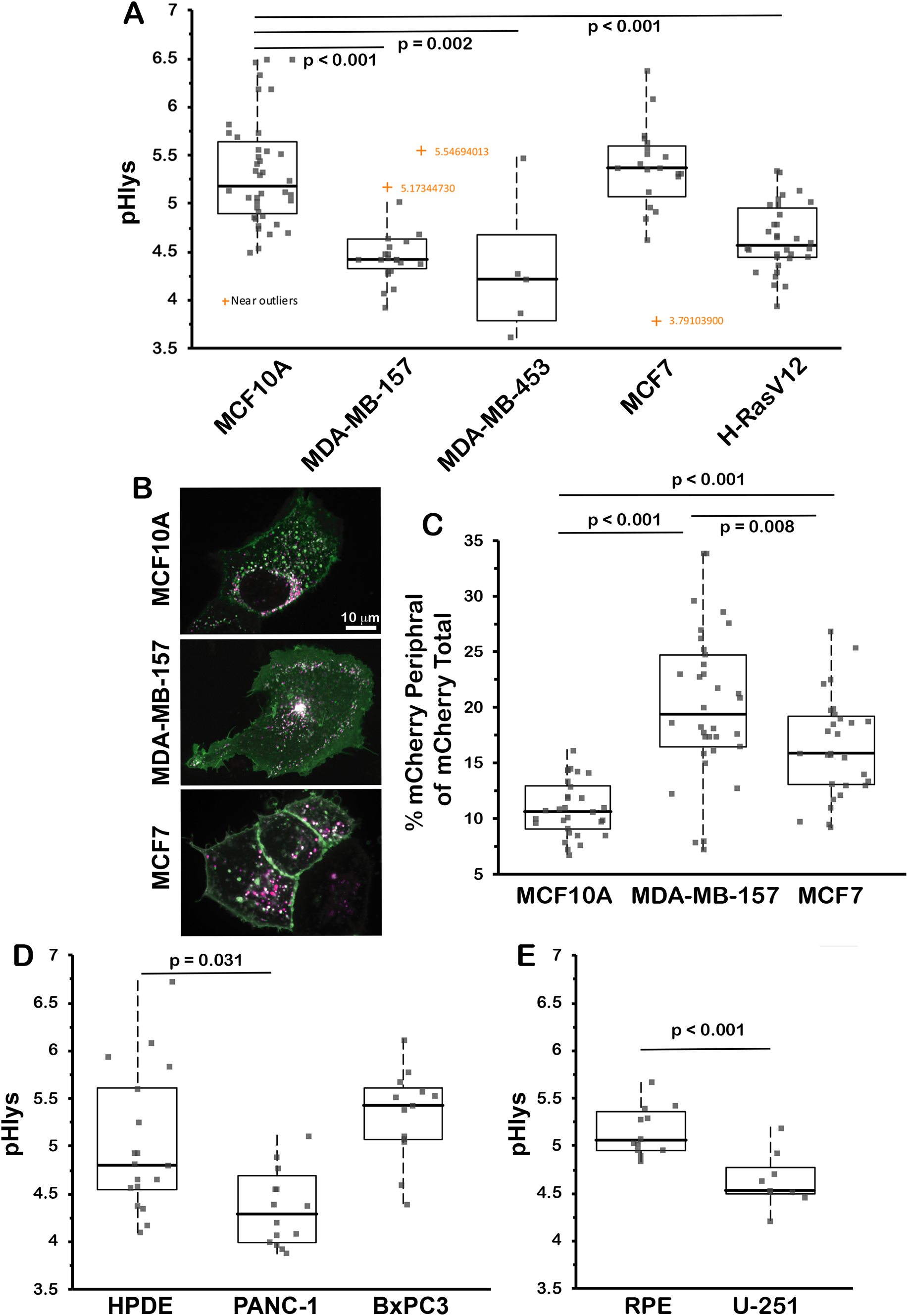
Differences in pHlys in cancer cell lines and with oncogene transformation. **(A)** The average pHlys of human untransformed MCF10A mammary epithelial cells (n = 36 cells), breast cancer MDA-MB-157 (n = 24), MDA-MB-453 (n = 5), and MCF7 (n = 21) cells, and MCF10A cells stably expressing H-RasV12 (n = 24). **(B, C)** Representative images of MCF10A, MDA-MB-157, and MCF7 cells expressing pHLARE (B) and quantification of lysosomes within 3 μm of the plasma membrane indicated by mCherry fluorescence (C). **(D)** The average pHlys of human untransformed HPDE pancreatic ductal epithelial cells (n = 19 cells) and pancreatic adenocarcinoma PANC-1 (n = 17) and BxPC3 (n = 13) cells. **(F)** The average pHlys of human untransformed RPE cells (n = 12 cells) and glioblastoma U-251 cells (n = 9 cells). Box plots show median, first and third quartile, with whiskers extending to observations within 1.5 times the interquartile range, and all individual data points. Statistical analysis by Tukey-Kramer HSD test, with data obtained from three to four separate preparations of all cells except for 2 preparations of MDA-MB-453.

The pHlys of PANC-1 but not BxPC3 pancreatic ductal carcinoma cells (4.39 ± 0.19 and 5.33 ± 0.14, respectively) was also significantly lower than HPDE normal pancreatic ductal epithelial cells (5.12 ± 0.18) (Fig. 5D). Additionally, the pHlys of U-251 human glioblastoma cells (4.67 ± 0.10) was significantly lower than RPE cells (Fig. 5E) and not different than PANC-1, MDA-MB-157 or MDA-MB-453 cells, although we did not use tissue matched untransformed cells to compare with U-251 cells. All measurements were made with cells in growth medium containing FBS, suggesting it the lower pHlys in the indicated cancer cells was not a result of limited nutrient availability.

The cancer cell lines described above have different mutational signatures. To determine whether a single activated oncogene is sufficient to change pHlys, we used MCF10A cells stably expressing H-RasV12. We previously reported that these cells have a higher cytosolic pH compared with wild-type MCF10A cells (Grillo-Hill et. al., 2015), which we confirmed in the current study (Fig. S3). We now found that pHlys is significantly lower (4.67 ± 0.07 compared with 5.23 ± 0.08) (Fig. 5F, G). These data indicate that a lower pHlys can be induced by a single activated oncogene and does not require the complex mutational signature of a cancer cell.

## Discussion

We report the design, validation, and use of pHLARE, a new genetically encoded pHlys biosensor. As a ratiometric biosensor localized predominantly in lysosomes with a broad dynamic range, pHLARE resolves a number of limitations of currently used fluorescent dyes for measuring pHlys, including improved organelle specificity, quantitative determinations, and stability with the ease of being stably expressed in cells. The lumenal pH of each organelle is different, requiring the use of organelle-specific pH indicators, and it crucial for organelle function, including the pH-dependent function of proteins that reflect adaptations to the cellular compartment where they are localized (Chan and Warwicker. 2009). The steady-state pHlys of ~5.0, the lowest of any cellular organelle, is particularly critical for the activity of lumenal acid-activated hydrolases, but recent findings suggest that pHlys may also contribute to Ca^2+^ homeostasis (Lee et al., 2015) as well as the localization of lysosomes in cells (Johnson et al., 2016), with the latter suggested to be important for cancer metastasis (Tu et al., 2008; Pu et al., 2016: Hamalisto and Jaattela, 2016). pHLARE was developed by fusing sfGFP and mCherry to lumenal and cytosolic domains, respectively, of LAMP1. Although LAMP1 is predominantly localized in organelles considered to be lysosomes or at the transition of late-endosomes to lysosomes, which we confirmed by co-localization of pHLARE with Rab7 that is a marker for late endosomes/lysosomes (Rabinowitz et al., 1992), recent findings indicate that LAMP1 is not exclusively in degradative lysosomes containing detectable hydrolases (Cheng et al., 2018; Yap et al., 2018). Hydrolase-deficient lysosomes raise questions on catabolic-independent functions requiring a low lumenal pH that use of pHLARE could contribute to resolving.

A number of diseases, including neurodegenerative disorders and cancer have dysregulated lysosome functions in catabolism, nutrient sensing, and trafficking (reviewed in Kallunki et al., 2012; Appelqvist et al., 2013; Perera and Zoncu 2016). Despite many of these functions being regulated by pHlys, whether pHlys is also dysregulated in these diseases, and the determinants and consequences of dysregulated pHlys remain incompletely understood. As a new tool, pHLARE could contribute to resolving a number of current questions on the role of lysosomes in disease pathologies. A constitutively increased pHlys is confirmed to occur with neurodegenerative disorders (Majumdar et al., 2007; Wolfe et al., 2013; Lee et al., 2015) and in diabetic nephropathy (Liu et al., 2015). The consequence of increased pHlys in these diseases is decreased activity of lumenal acid-activated hydrolases and catabolism of proteins and macromolecules.

Whether pHlys is dysregulated in cancer cells, which rely on increased catabolism of macromolecules to generate biomass for rapid proliferation, remains unresolved. To our knowledge, only one report compared pHlys in tissue matched normal and cancer cells and found that clonal head and neck cancer cells have a higher pHlys than untransformed keratinocytes (Nilsson et al., 2010). However, in this previous study pHlys was determined after flow cytometry of cells loaded with FITC-conjugated dextran, and flow cytometry requires disrupting cell-matrix adhesions and using cells in suspension, which could change steady-state pHlys. We used pHLARE in adherent cells maintained in growth medium to show that MDA-MB-157 and MDA-MB-453 human breast cancer cells and PANC-1 human pancreatic cancer cells had a significantly lower pHlys compared with tissue-matched untransformed MCF10A and HPDE cells, respectively. We also observed a lower pHlys in U-251 human glioblastoma cells and in human mammary epithelial cells expressing oncogenic H-RasV12. However, the pHlys of MCF7 human breast cancer cells and BxPC3 human pancreatic cells was not different compared with tissue-matched untransformed cells. Additionally, clonal HeLa cervical cancer cells have a normal-like pHlys of 5.1-5.2 (Ostrowski et al., 2016; Johnson et al., 2016). The causes and consequences of these differences in pHlys between distinct cancer cells remain to be determined. Although MCF7 cells are estrogen receptor- and progesterone receptor-positive and MCA-MB-157 and MDA-MB-453 are triple negative, their different pHlys is likely not determined by their mutational signature, as suggested by a single oncogene, RasV12, inducing a lower pHlys. However, different pHlys values could be related to invasiveness. MDA-MB-157 cells are basal-type and more mesenchymal and invasive than lumenal MCF7 cells that have tight cell-cell junctions (Neve et al., 2006) Also, PANC-1 cells are more invasive and migrate as single cells compared with BxPC3 cells that have tighter cell-cell contacts and migrate as collective cell sheets [Deer et al., 2010]. Lysosomes can contribute to cell invasion by the endocytic release of lysosomal hydrolases degrading the extracellular matrix (Tu et al., 2008; Pu et al., 2016; Hamalisto and Jaattela, 2016), and we found that MDA-MB-157 have significantly more pHLARE within 3 μm of the plasma membrane compared with MCF7 and MCF10A cells.

Differences in pHlys between cancer cells could also reflect drug resistance, as lysosomes contribute to resistance by sequestering and degrading chemotherapeutics (Zhitomirsky and Assaraf, 2016; Piao and Amaravadi, 2016). We favor the possibility that differences in pHlys are related to cancer cell-specific metabolic requirements and cellular mechanisms for generating biomass, according to the view that lysosomes contribute to metabolic reprogramming in some cancer cells along with changes in glycolysis and mitochondrial respiration (Perera and Bardeesy, 2015). Supporting the prediction that a lower pHlys may function as an alternative metabolic reprogramming mechanism, BxPC3 cells but not PANC-1 cells rely on a classically defined metabolic reprogramming of increased glycolysis and decreased mitochondrial respiration (Kovalenko et al., 2016; Tataranni et al., 2017), and inhibiting glycolysis decreases proliferation of BxPC3 cells but not PANC-1 cells (Tataranni et al., 2017, Nishi et al,, 2018).

As a new reagent to more accurately measure pHlys that can be propagated in cells and used over long time periods, pHLARE will be invaluable for many objectives. First is to identify resident lysosome proteins regulating pHlys. The V-ATPase is currently the only well-accepted regulator of pHlys, with roles for a number of lysosome-localized transporters and pumps being controversial (Xu and Ren, 2015). Endogenous regulators of pHlys could be targets for therapeutics to restore dysregulated pHlys in diseases such as neurodegeneration and cancer. Second is to determine lysosome exocytosis, particularly in the context of cancer metastasis, by measuring increased sfGFP fluorescence upon exposure to the extracellular space. Third, because pHLARE is genetically encoded it can be expressed in animal models to study lysosome dynamics in diseases and also pHlys dynamics during development, which remains unknown.

## Methods and Materials

### Cloning and DNA constructs

Mammalian expression plasmids encoding rat LAMP1 (mCherry-Lysosomes-20, Addgene; 55073), mApple-LAMP1-pHluorin (Addgene; 54918), superfolderGFP (sfGFP, Addgene; 54579) were obtained from the Michael Davidson collection at UCSF. Gibson assembly was used to generate pHLARE. Briefly, PCR was used to amplify the prolactin endoplasmic reticulum (ER) signal peptide (MDSKGSSQKGSRLLLLLVVSNLLLCQGVVS) from mApple-LAMP1-pHluorin, sfGFP from sfGFP-C1, and rat LAMP1 without the ER signal peptide (amino acids 22-407) from mCherry-Lysosome-20. The PCR inserts were assembled with restriction enzyme digested pmCherry-N1 backbone in Gibson Assembly Master Mix (New England Biolabs; E2611S) as per manufacturer recommendations and the assembled plasmids transformed into NEB5*a* cells (New England Biolabs; C3019). DNA sequencing was performed to verify the integrity of the construct.

### Cell culture and transfection

All cell lines were maintained at 37°C and 5% CO2 except MDA-MB-157 and MDA-MB-453, which were maintained at 37°C and atmospheric CO2. Media obtained from Thermo Fisher Scientific were supplemented with Penicillin-Streptomycin. RPE cells, obtained from Sophie Dumont (UCSF), and MCF7 cells, obtained from the UCSF Cell Culture Facility, were maintained in DMEM/F21 supplemented with 10% heat inactivated fetal bovine serum (FBS, Atlanta Biological) and 2 mM L-Glutamine. MDCK cells, obtained from the UCSF Cell Culture Facility, and PANC-1 cells, obtained from Rushika Perera (UCSF) were maintained in DMEM-H21 medium supplemented with 10% FBS. NIH-3T3 cells, obtained from Jeroen Roose (UCSF), were maintained in McCoy’s 5a medium supplemented with 10% FBS. MDA-MB-157 cells and MDA-MB-453 cells, obtained from the UCSF Cell Culture Facility and from ATCC, respectively, were maintained in Leibovitz L15 medium supplemented with 15% FBS. HDPE cells, obtained from Rushika Perera (UCSF), were maintained Keratinocyte medium supplemented with 10% FBS. MCF10A cells, wild type and stably expressing RasV12, were obtained from Jay Debnath (UCSF) and maintained and studied in 2D cultures as previously described (Debnath et al., 2003; Grillo-Hill et al., 2015). Heterologous expression of proteins was achieved by transfecting cells with 1 □g of plasmid DNA/35 mm using FuGENE HD (Promega; E2311), as previously described (Webb et al., 2015). For transient expression of pHLARE, cells were washed 24 h after transfecting and imaged after an additional 24 h. For stable expression of pHLARE, 24 h after transfection cells were sorted for sfGFP and mCherry fluorescence on a FACS Aria II (BD Biosciences) at the UCSF Parnassus Flow Cytometry Core.

### Immunolabeling and Immunoblotting

For immunolabeling, RPE cells stably expressing pHLARE were plated at a density of ~20 × 104 cells/well of a 6-well plate containing 100 mm glass coverslips were maintained in complete growth media in a 37°C/5% CO2 incubator. Cells plated for 48 h were fixed in 4% paraformaldehyde in PBS for 10 min, permeabilized in 0.1% Triton X-100 in PBS for 10 min, and incubated with blocking buffer (3% bovine serum albumin, 1% non-immune goat serum, and 1% cold water fish gelatin) for 1 hour. After washing in PBS, cells were incubated overnight at 4°C with primary antibodies to mCherry (Thermo Fisher Scientific; M11217) and human LAMP1 (Cell Signaling Technologies; 9091) diluted in blocking buffer. After 24 h cells were washed with PBS and incubated with Alexa Fluor 568 goat anti-mouse (Thermo Fisher Scientific; A-11077) and Alexa Fluor 647 goat anti-rabbit (Thermo Fisher Scientific; A-21244) for 1 h. Coverslips were washed 3X in PBS, with Hoechst 33342 (Thermo Fisher Scientific; 62249) included in the second wash. and mounted on glass sides in ProLong gold antifade mountant (Thermo Fisher Scientific; P36930). Where indicated, RPE cells in growth medium were incubated with far-red SiR lysosome (Cytoskeleton) for 60 min and washed before imaging live cells expressing pHLARE. Images were acquired by spinning disk confocal microscopy as described for imaging pHLARE.

For immunoblotting, proteins in total cell lysates were separated by SDS-PAGE and transferred to polyvinylidene difluoride membranes. The membranes were incubated for 24 h at 4°C with antibodies to RFP (Abcam 62341) or total and phosphorylated S6K1 (Cell Signaling Technologies; 9292 and 9205, respectively). After washing, membranes were incubated with peroxidase-conjugated secondary antibodies (Jackson ImmunoResearch Laboratories, Inc.) for 1 h at RT, and bound antibodies were developed by enhanced chemiluminescence using SuperSignal West Femto (Thermo Fisher Scientific) and imaged using an Alpha Innotech FluorChem Q (Alpha Innotech). Densitometery measurements were made using Image J software.

### Microscopy and pH measurements

For live-cell imaging, cells were plated on 35 mm MatTek dishes catalog number (MatTek Corporation; P35G-1.5-10-C). Fluorescent images were acquired using a customized spinning disk confocal (Yokogawa CSU-X1) on a Nikon Ti-E microscope with a 60X Plan TIRF 1.49NA objective equipped with a Photometrics cMYO cooled CCD camera, as described (Stehbens et al., 2012. Image acquisition settings that caused less than 5% loss of signal were used in further experiments. Conditions for experimentally changing pHlys included incubating cells for 2 h prior to imaging with 100 nM bafilomycin A1 (Sigma-Aldrich; B1793), 100 ∝M chloroquine (Sigma-Aldrich; C6628), or 250 nM Torin-1 (Tocris Bioscience; 4247) added to growth medium.

Fluorescence ratios of pHLARE were first acquired in cell culture medium and then cells were sequentially incubated in a potassium-phosphate buffer (50 mM potassium phosphate, 80 mM potassium chloride, 1 mM magnesium chloride) containing 20 μM nigericin free acid (Thermo Fisher Scientific; N1495) at pH 6.5 and 5.0. Cells were incubated in nigericin buffers for 5 minutes before imaging to equilibrate extracellular and intracellular pH.

Intracellular pH was measured as previously described (Grillo-Hill et al., 2015; White et al., 2017a). In brief, cells plated for 48 h in 24-well dishes were washed and incubated for 15 min in a NaHCO_3_-containing buffer containing 1 □M of the pH-sensitive dye BCECF. BCECF fluorescence ratios, measured on a SpectraMax fluorescence plate reader (Molecular Devices), were calibrated to pHi using nigericin-containing buffers.

### Image analysis

To reproducibly define lysosome objects and corresponding local background regions that were then used to calculate fluorescence ratios and pHlys values, we implemented an image analysis algorithm in NIS Elements software based on mCherry fluorescence intensity thresholding and morphometric manipulation of the resulting binary images. Briefly, the mCherry channel was thresholded at an intensity three times above the non-lysosomal cell membrane pHLARE background. The thresholded binary was smoothened and dilated such that each lysosome area included sufficient surrounding background, and a corresponding local background was defined as the outer 1-pixel contour of each lysosome object. Fluorescent intensity measurements from both pHLARE channels of all lysosome and corresponding local background objects were exported into Microsoft Excel for further analysis.

The pHLARE fluorescence ratio r for each lysosome object was calculated as

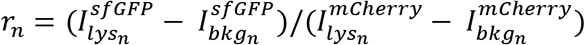

in which n is the index for each lysosome object, I_lys_ is the fluorescence intensity of the lysosome object and I_bkg_ the fluorescence intensity of the corresponding local background. The lysosome pH was calculated in two different ways, either using the mean local background signal or the mean lysosome signal minus camera offset of all lysosome objects to calibrate the pHLARE fluorescence ratio based on the fluorescence intensities in the nigericin images. pH values for each lysosome object were then calculated assuming a linear relationship between pH and pHLARE fluorescence ratio. To reduce the impact of measurement error outliers, the cellular lysosome pH was calculated as the median of all individual lysosome object pH values per cell. An NIS Elements Macro ‘pHLARE_analysis.mac’ performing the image operations semi-automatically that also allows masking the cell of interest in the measurement and nigericin images, an example Excel worksheet for the pHlys calculations ‘pHLARE_analysis_worksheet.xlsx’, an example data set ‘RPE_cell_6.zip’ as well as further step-by-step instructions on how to utilize the macro and excel worksheet are included in the supplemental information.

To determine spatially localized pHlys in RPE cells we used the analysis pipeline described above to measure lysosomes selectively within 3 μm of the distal margin of membrane protrusions and 3 μm of the nuclear membrane, and in MDA-MB-157 and MCF10A cells we measured the total mCherry fluorescence of pHLARE within 3 μm of the plasma membrane relative to the total mCherry fluorescence of the cell. To determine lysosome size, we also used the analysis program described above that includes the area of lysosome objects. To evaluate the size of single lysosomes and not lysosome clusters we used only the area of objects with a shape value between 0.9 to 1.0.

### Data presentation and statistical analysis

Box-and whisker plots were generated using Analyse-It for Microsoft Excel and show median, first and third quartile, observations within 1.5 times the interquartile range, and all individual data points. Significance of multiple comparisons was calculated by Tukey– Kramer honest significant difference (HSD) test in Analyse-It for Microsoft Excel or by Student’s Paired *t*-test. Figures were assembled in Adobe Illustrator CS5.

## Supporting information

supplementary data analysis

## Abbreviations used

pHlys: lysosome pH
mTORC1: mammalian target of rapamycin complex 1
pHLARE: pH lysosome activity reporter
lysosomal associated membrane protein 1: LAMP1
S6K1: ribosomal protein S6 kinase beta-1

## Acknowledgments

We gratefully acknowledge the contributions of technical assistance and expertise from Andreas Ettinger (Helmholtz Center Munich, Germany) and cell lines from Sophie Dumont, Rushika Perera, Jeroen Roose and Jay Debnath (UCSF). This work was supported by National Institutes of Health grants CA197855 (D.L.B.) and NS107480 and S10RR026758 (T.W), and by funding from the Paul G. Allen Family Foundation. The authors declare no competing financial interests.

## Author contributions

B.W. developed pHLARE, validated properties, and with D.L.B. and J.C. acquired data. T.W developed the pHLARE image analysis pipeline and with D.L.B and J.C. performed image analysis. All authors contributed to writing the manuscript

---

**Figure S1.**
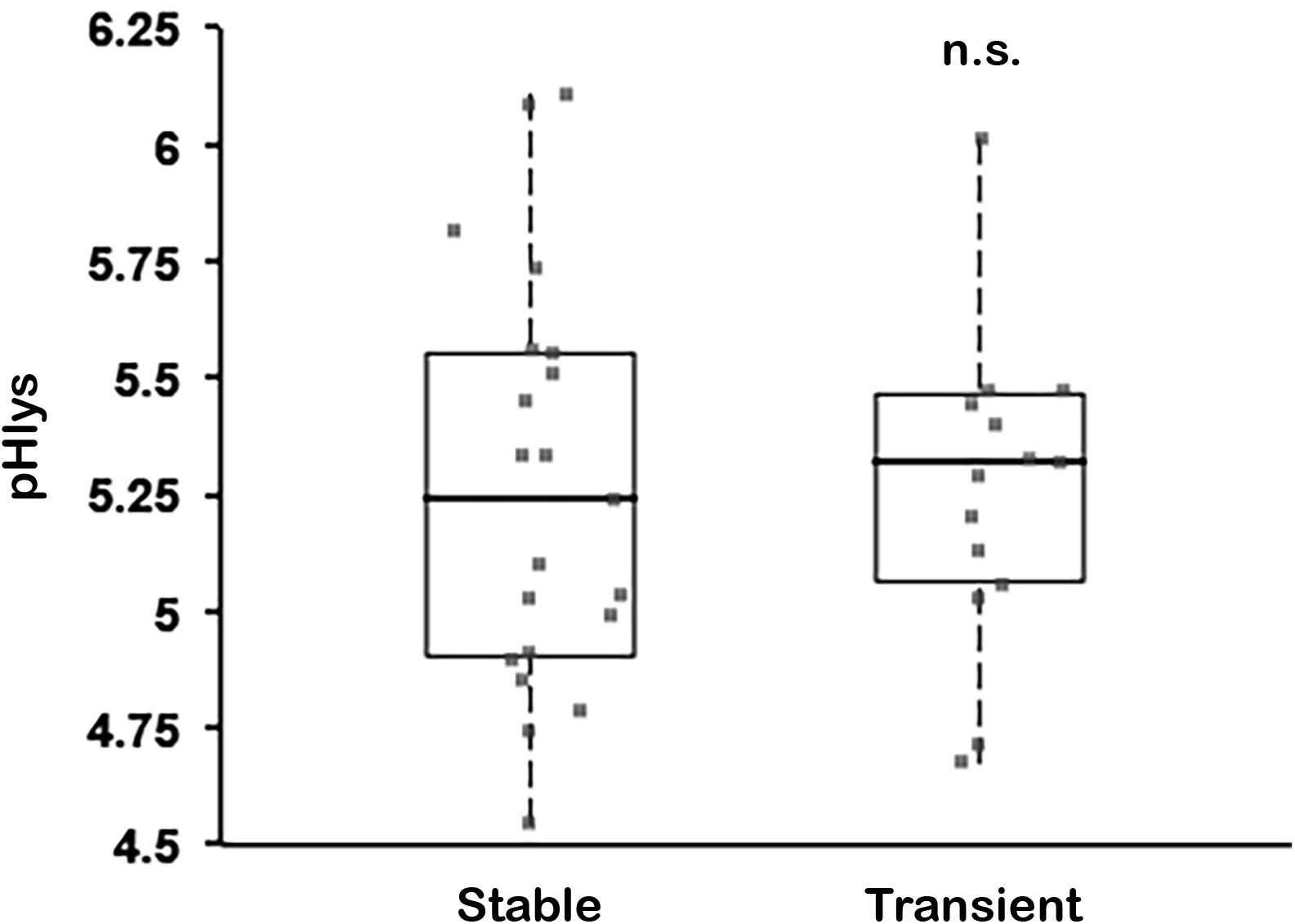
pHlys is similar in MCF10A cells stably or transiently expressing pHLARE. Average steady-state pHlys in MCF10A cells. Box plots show median, first and third quartile, with whiskers extending to observations within 1.5 times the interquartile range, and all individual data points include data for individual cells (stable, n = 18 cells; transient, n=14 cells) obtained from three separate cell preparations. Statistical analysis by Tukey-Kramer HSD test.

**Figure S2.**
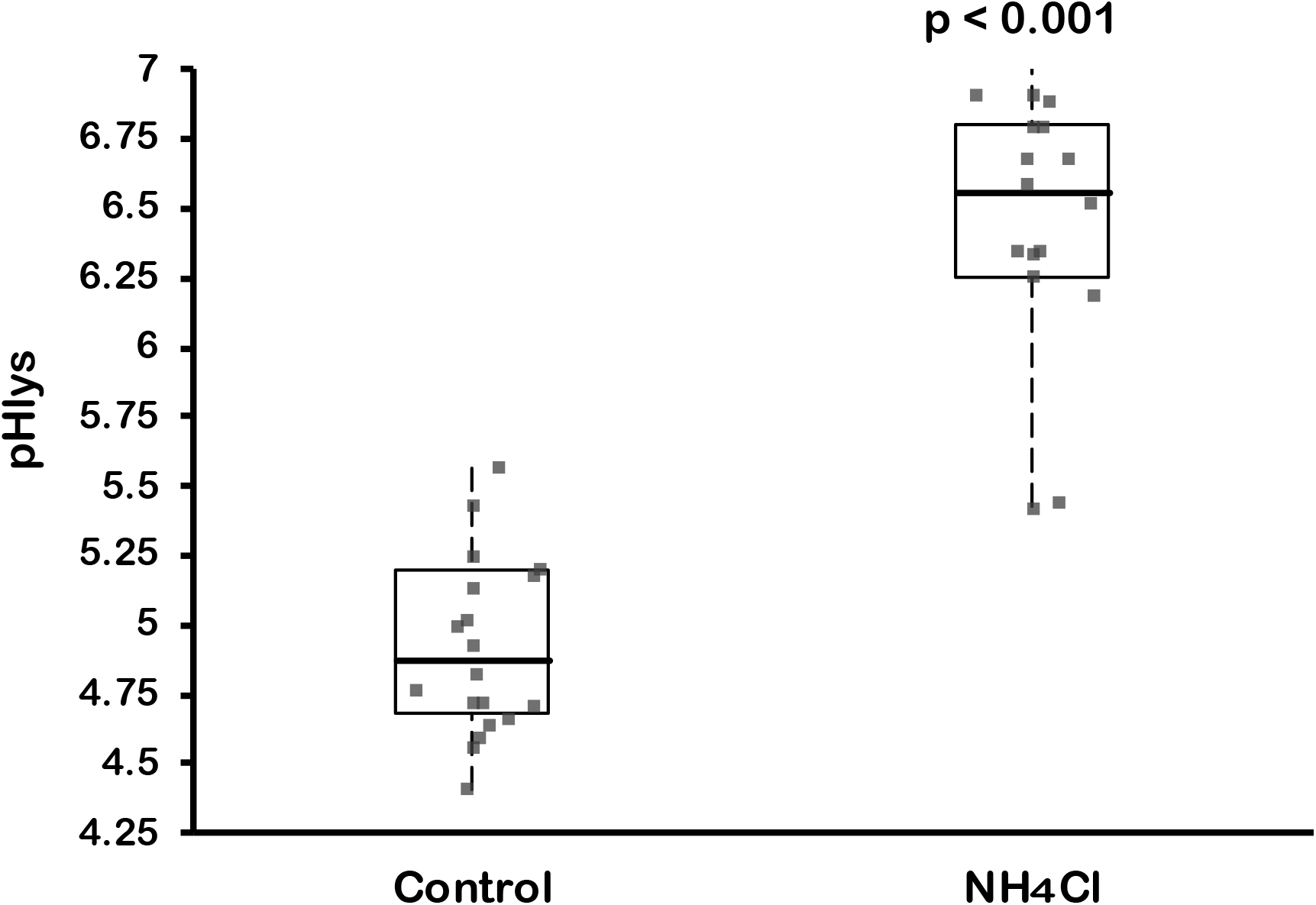
NH4Cl increases pHlys. Average pHlys in RPE cells stably expressing pHLARE in control medium (n = 20 cells) or medium containing 5 mM NH4Cl for 18 h (n=18 cells). Box plots show median, first and third quartile, with whiskers extending to observations within 1.5 times the interquartile range, and all individual data points include data for individual cells (each closed circle) obtained from three separate cell preparations. Statistical analysis by Tukey-Kramer HSD test.

**Figure S3.**
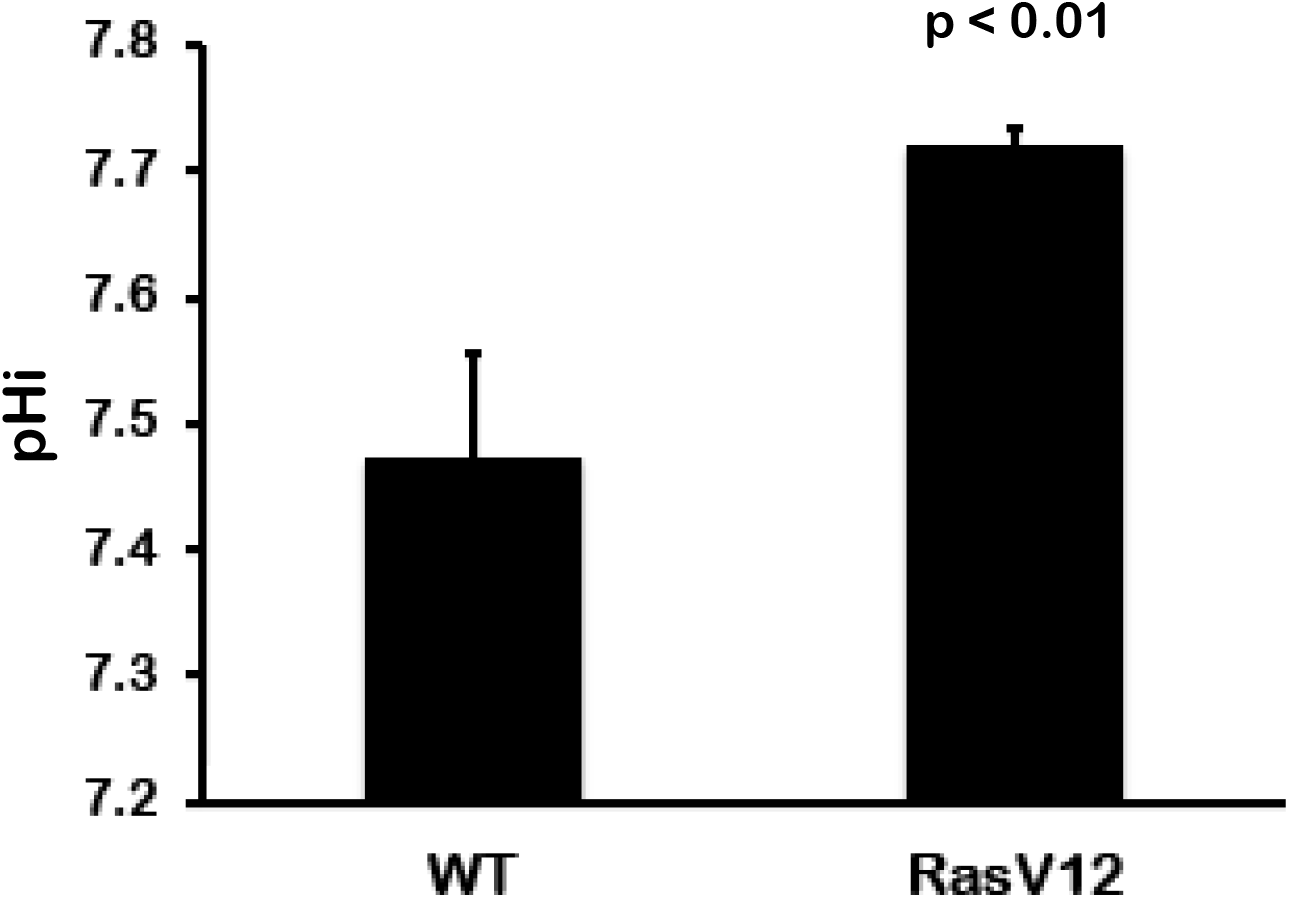
MCF10A cells expressing H-RasV12 have increased cytosolic pH. Cytosolic pH was determined in MCF10A control cells and cells stably expressing H-Ras-V12 loaded with the pH-sensitive dye BCECF in a NaHCO3-containing buffer. Data are the means ± s.e.m. obtained from three separate cell preparations. Statistical analysis by Student’s Paired *t*-test.

## Notes

### Competing Interest Statement

The authors have declared no competing interest.

## References

Altan, N., Y. Chen, M. Schindler, and S.M Simon. 1999. Tamoxifen inhibits acidification in cells independent of the estrogen receptor. Proc Natl Acad Sci U S A. 96:4432–4437.

Appelqvist, H., P. Wäster, K. Kågedal and K. Öllinger. 2013. The lysosome: from waste bag to potential therapeutic target. J. Mol. Cell Biol. 5:214–26.

Bandyopadhyay, D., A. Cyphersmith, Zapata JA, Kim YJ, Payne CK. 2014. Lysosome transport as a function of lysosome diameter. PLoS One. 9(1):e86847.

Cheng X.T., Y.X. Xie, B. Zhou, N. Huang, T. Farfel-Becker, Z.H. Sheng. 2018. Characterization of LAMP-1 labeled nondegradative lysosomal and endocytic compartments in neurons. J. Cell Biol. 217(9):3127–3139.

Colacurcio D.J. and R.A. Nixon. 2016. Disorders of lysosomal acidification - The emerging role of v-ATPase in aging and neurodegenerative disease. Ageing Res Rev. 32:75–88.

Chan, P., and J. Warwicker. 2009. Evidence for the adaptation of protein pH-dependence to subcellular pH. BMC Biol. 7:69–78.

Daniel, C., C. Bell, C. Burton, S. Harguindey, S.J Reshkin, and C. Rauch. 2013. The role of proton dynamics in the development and maintenance of multidrug resistance in cancer. Biochim. Biophys. Acta. 1832:606–617.

Debnath, J., S.K. Muthuswamy, and J.S. Brugge. 2003. Morphogenesis and oncogenesis of MCF-10A mammary epithelial acini grown in three-dimensional basement membrane cultures. Methods 30:256–268.

Deer, E.L., J. Gonzalez-Hernandez, J.D. Coursen, J.E. Shea, J. Ngatia, C.L Scaife, M.A. Firpo, and S.J. Mulvihill. 2010. Phenotype and genotype of pancreatic cancer cell lines. Pancreas. 39:425–35.

Fraldi, A., A.D. Klein, D.L. Medina, C. Settembre. 2016. Brain disorders due to lysosomal dysfunction. Annu Rev Neurosci. 39:277–95.

Grillo-Hill, B.K., B.A. Webb, and D.L. Barber. 2014. Ratiometric imaging of pH probes. Methods Cell Biol. 123:429–48.

Grillo-Hill, B.K., C. Choi, M. Jimenez-Vidal, and D.L. Barber. 2015 Increased H+ efflux is sufficient to induce dysplasia and necessary for viability with oncogene expression. eLife 4:e03270.

Hamalisto, S. and M. Jaattela. 2016. Lysosomes in cancer-living on the edge (of the cell). Curr. Opin. Cell Biol. 39:69–76.

Jaiswal, J.K, N.W. Andrews, and S. M. Simon. 2002. Membrane proximal lysosomes are the major vesicles responsible for calcium-dependent exocytosis in nonsecretory cells. J. Cell Biol. 159:625–35.

Johnson, D.E., P. Ostrowski, V. Jaumouille, and S. Grinstein. 2016. The position of lysosomes within the cell determines their luminal pH. J. Cell Biol. 212:677–92.

Kallunki, T., O.D. Olsen, and M. Jaattela. 2012 Cancer-associated lysosomal changes: friends or foes? Oncogene 32:1995–2004.

Kenific C.M. and J. Debnath. 2015. Cellular and metabolic functions for autophagy in cancer cells. Trends Cell Biol. 25(1):37–45.

Kornak, U., D. Kasper, M.R Bösl, E. Kaiser, M. Schweizer, A. Schulz, W. Friedrich, G. Delling, and T.J. Jentsch. 2001. Loss of the ClC-7 chloride channel leads to osteopetrosis in mice and man. Cell. 104:205–215.

Kovalenko, I., A. Glasauer, L. Schöckel, D.R. Sauter, A. Ehrmann, F. Sohler, A. Hägebarth I. Novak I, and S. Christian PLoS One. 2016. Identification of KCa3.1 Channel as a Novel Regulator of Oxidative Phosphorylation in a Subset of Pancreatic Carcinoma Cell Lines. 11(8):e0160658.

Lee, J.H., M.K. McBrayer, D.M. Wolfe, L.J Haslett, A. Kumar, Y. Sato, P.P Lie, P. Mohan, E.E. Coffey, U. Kompella, C.H. Mitchell, E. Lloyd-Evans, and R.A. Nixon. 2015. Presenilin 1 maintains lysosomal Ca2+ homeostasis via TRPML1 by regulating vATPase-mediated lysosome acidification. Cell Rep. 12:1430–44.

Lin, HJ, P. Herman, J.R. Lakowicz. 2003. Fluorescence lifetime-resolved pH imaging of living cells. Cytometry A. 2003 Apr;52(2):77–89.

Liu B, Palmfeldt J, Lin L, Colaço A, Clemmensen KKB, Huang J, Xu F, Liu X, Maeda K, Luo Y, Jäättelä M. 2018. STAT3 associates with vacuolar H_+_-ATPase and regulates cytosolic and lysosomal pH. Cell Res. 28(10):996–1012.

Liu, W.J., T.T. Shen, R.H. Chen, H.L Wu, Y.J. Wang, J.K. Deng, Q.H. Chen, Q. Pan, C.M. Huang, C.M. Fu, J.L Tao, D. Liang D and H.F. Liu. 2015. Autophagy-lysosome pathway in renal tubular epithelial cells is disrupted by advanced glycation end products in diabetic nephropathy. J. Biol. Chem. 290:20499–510.

Ma, X., Y. Wang, T. Zhao, Y. Li, L.C. Su, Z. Wang, G. Huang, B.D. Sumer, and J. Gao. 2014. Ultra-pH sensitive nanoprobe library with broad pH tenability and fluorescence emissions. J. Am. Chem. Soc. 136:11085–92.

Mu, F.T., J.M. Callaghan, O. Steele-Mortimer, O., et al. 1995. EEA1, an early endosome-associated protein. EEA1 is a conserved alpha-helical peripheral membrane protein flanked by cysteine ‘fingers’ and contains a calmodulin binding IQ motif. J. Biol. Chem. 270:13503–13511.

Majumdar, A., D. Cruz, N. Asamoah, A. Buxbaum, I. Sohar, P. Lobel, and F.R. Maxfield. 2007. Activation of microglia acidifies lysosomes and leads to degradation of Alzheimer amyloid fibrils. Mol. Biol. Cell 18:1490–96.

Marshansky, V., and M. Futai. 2008. The V-type Hfl-ATPase in vesicular trafficking: Targeting, regulation and function. Curr. Opin. Cell Biol. 20: 415–26.

Mindell, J.A. 2012. Lysosomal acidification mechanisms. Annu. Rev. Physiol. 74:69–86.

Neve, R.M., K. Chin, J. Fridlyand, J. Yeh, F.L. Baehner FL, Fevr T, Clark, N. Bayani, J.P Coppe, F. Tong, T. Speed, P.T. Spellman, S. DeVries, A. Lapuk, N.J. Wang, W.L. Kuo, J.L. Stilwell, D. Pinkel, D.G. Albertson, F.M. Waldman, F. McCormick, R.B. Dickson, M.D. Johnson, M. Lippman, S. Ethier, A. Gazdar, J.W. Gray. 2006. A collection of breast cancer cell lines for the study of functionally distinct cancer subtypes. Cancer Cell. 10(6):515–27.

Nilsson, C., K. Roberg, R.C. Grafström, and K. Ollinger. 2010. Intrinsic differences in cisplatin sensitivity of head and neck cancer cell lines: Correlation to lysosomal pH. Head Neck. 32:1185–1194.

Nishi, K., M. Suzuki, N. Yamamoto, A. Matsumoto, Y. Iwase, K. Yamasaki, M. Otagiri, and N. Yumita. 2018. Glutamine deprivation enhances acetyl-CoA carboxylase inhibitor-induced death of human pancreatic cancer cells. Anticancer Res. 38:6683–6689.

Perera, R.M. and N. Bardeesy. 2015. Pancreatic cancer metabolism: Breaking it down to build it back up. Cancer Discov. 5:1247–1261.

Perera, R.M. and R. Zoncu. 2016. The lysosome as a regulatory hub. Annu. Rev. Cell Dev. Biol. 32:223–53.

Piao, S. and R.K. Amaravadi. 2016. Targeting the lysosome in cancer. Ann. N. Y. Acad. Sci. 1371:45–54.

Pu, J., C.M. Guardia, T. Keren-Kaplan, and J.S. Bonifacino. 2016. Mechanisms and functions of lysosome positioning. J. Cell Sci. 129:4329–39.

Rebsamen, M., L. Pochini, T. Stasyk, M.E. de Araujo, M. Galluccio, R.K. Kandasamy, B. Snijder, A. Fauster, E.L Rudashevskaya, M. Bruckner, S. Scorzoni, P.A. Filipek, K.V. Huber, J.W. Bigenzahn, L.X. Heinz, C. Kraft, K.L. Bennett, C. Indiveri, L.A. Huber, and G. Superti-Furga. 2015. SLC38A9 is a component of the lysosomal amino acid sensing machinery that controls mTORC1. Nature. 519:477–481.

Rodríguez, A., P. Webster, J. Ortego, and N.W. Andrews. 1997. Lysosomes behave as Ca2+-regulated exocytic vesicles in fibroblasts and epithelial cells. J. Cell Biol. 137:93–104.

Rohrer J., A. Schweizer, D. Russell, S. Kornfeld. 1996. The targeting of Lamp1 to lysosomes is dependent on the spacing of its cytoplasmic tail tyrosine sorting motif relative to the membrane. J. Cell Biol. 132(4):565–76.

Sardiello, M., M. Palmieri, A. di Ronza, D.L. Medina, M. Valenza, V.A. Gennarino, C. Di Malta, F. Donaudy, V. Embrione, R.S. Polishchuk, S. Banfi, G. Parenti, E. Cattaneo, and A. Ballabio. 2009. A gene network regulating lysosomal biogenesis and function. Science. 325(5939):473–477.

Saxton, R.A., and D.M. Sabatini. 2017. mTOR signaling in growth, metabolism and disease. Cell. 168:960–976.

Settembre, C., R. De Cegli, G. Mansueto, P.K. Saha, F. Vetrini, O. Visvikis, T. Huynh, A. Carissimo, D. Palmer, T.J. Klisch, A.C. Wollenberg, D. Di Bernardo, L. Chan, J.E. Irazoqui, and A. Ballabio. 2013. TFEB controls cellular lipid metabolism through a starvation-induced autoregulatory loop. Nat. Cell Biol. 15:647–658.

Scott, C.C., and J. Gruenberg. 2013. Ion flux and the function of endosomes and lysosomes: pH is just the start: the flux of ions across endosomal membranes influences endosome function not only through regulation of the luminal pH. Bioessays. 33(2):103–110.

Stehbens, S., H. Pemble, L. Murrow, and T. Wittmann. 2012. Imaging intracellular protein dynamics by spinning disk confocal microscopy. Methods Enzymol. 504:293–313.

Tataranni, T., F. Agriesti, V. Ruggieri, C. Mazzoccoli, V. Simeon, I. Laurenzana, R. Scrima, V. Pazienza, N. Capitanio, and C. Piccoli. 2017. Rewiring carbohydrate catabolism differentially affects survival of pancreatic cancer cell lines with diverse metabolic profiles. Oncotarget. 8:41265–41281.

Thoreen, C.C., L. Chantranupong, H.R. Keys, T. Wang, N.S. Gray, and D.M. Sabatini. 2012. A unifying model for mTORC1-mediated regulation of mRNA translation. Nature. 485:109–113.

Thoreen, C.C., S.A. Kang, J.W. Chang, Q. Liu, J. Zhang, Y. Gao, L.J. Reichling, T. Sim, D.M. Sabatini, and N.S. Gray. 2009. An ATP-competitive mammalian target of rapamycin inhibitor reveals rapamycin-resistant functions of mTORC1. J. Biol. Chem. 284:8023–8032.

Tu, C., C.F. Ortega-Cava, G. Chen, N.D. Fernandes, D. Cavallo-Medved, B.F. Sloane, V. Band, and H. Band. 2008. Lysosomal cathepsin B participates in the podosome-mediated extracellular matrix degradation and invasion via secreted lysosomes in v-Src fibroblasts. Cancer Res. 68:9147–9156.

Walton, Z.E., C.H. Patel, R.C. Brooks, Y. Yu, A. Ibrahim-Hashim, M. Riddle, A. Porcu, T. Jiang, B.L. Ecker, F. Tameire, C. Koumenis, A.T Weeraratna, D.K. Welsh, R. Gillies, J.C. Alwine, L. Zhang, J.D. Powell, and C.V Dang. 2018. Acid Suspends the Circadian Clock in Hypoxia through Inhibition of mTOR. Cell. 174:72–87 e32.

Wang, C., B. Dong, X. Kong, N. Zhang, W. Song, W. Lin. 2018. Dual site-controlled two-photon fluorescent probe for the imaging of lysosomal pH in living cells. Luminescence. 33:1275–1280.

Wang, S., Z.Y. Tsun, R.L. Wolfson, K. Shen, G.A. Wyant, M.E. Plovanich, E.D. Yuan, T.D. Jones, L. Chantranupong, W. Comb, T. Wang, L. Bar-Peled, R. Zoncu, C. Straub, C. Kim, J. Park, B.L Sabatini, D.M. Sabatini. 2015. Metabolism. Lysosomal amino acid transporter SLC38A9 signals arginine sufficiency to mTORC1. Science 347:188–194.

Webb, B.E., M. Chimenti, M.P. Jacobson, and D.L. Barber. 2011. Dysregulated pH: a perfect storm for cancer progression. Nature Cancer Rev. 11:671–677.

Webb, B.A., F. Forouhar, F.E. Szu, J. Seetharaman, L. Tong, and D.L. Barber. 2015. Structures of human phosphofructokinase-1 and atomic basis of cancer-associated mutations. Nature 523:111–114.

White, KA, R.G. Garrido Ruiz, Z.A. Szpiech, N.B. Strauli, R.D. Hernandez, J.P Jacobson, D.L. Barber. 2017a. Cancer-associated arginine to histidine mutations confer a gain in pH sensing to mutant proteins. Sci Signaling 10(495). pii: eaam9931.

White, K.A., B.K. Grillo-Hill, and D.L. Barber. 2017b. Dysregulated pH dynamics enables cancer cell behaviors. J. Cell Sci. 130:663–669.

Wolfe, D.M., J.H. Lee, A. Kumar, S. Lee, S.J. Orenstein, and R.A. Nixon. 2013. Autophagy failure in Alzheimer’s disease and the role of defective lysosomal acidification. Eur. J. Neurosci. 37:1949–1961.

Xu, H. and D. Ren. Lysosomal physiology. 2015. Annu. Rev. Physiol. 77:57–80.

Yap C.C., L. Digilio, L.P. McMahon, A.D.R. Garcia, B. Winckler. 2018. Degradation of dendritic cargos requires Rab7-dependent transport to somatic lysosomes. J. Cell Biol. 217(9):3141–3159.

Yu, L., C.K. McPhee, L. Zheng, G.A. Mardones, Y. Rong, J. Peng, N. Mi, Y. Zhao, Z. Liu, F. Wan, D.W. Hailey, V. Oorschot, J. Klumperman, E. H. Baehrecke, and M. J. Lenardo. 2010. Termination of autophagy and reformation of lysosomes regulated by mTOR. Nature 465:942–46.

Zhitomirsky, B., and Y.G Assaraf. 2016. Lysosomes as mediators of drug resistance in cancer. Drug Resist. Updat. 24:23–33.

Zhitomirsky, B., H. Farber, and Y.G. Assaraf. 2018. LysoTracker and MitoTracker Red are transport substrates of P-glycoprotein: implications for anticancer drug design evading multidrug resistance. J. Cell Mol. Med. 22:2131–2141.

Zhou, J., S.H. Tan, V. Nicolas, C. Bauvy, N.D. Yang, J. Shang, Y., Xue, P. Codogno, and H.M. Shen. 2013. Activation of lysosomal function in the course of autophagy via mTORC1 suppression and autophagosome-lysosome fusion. Cell Res. 23:508–23.

