## supplementary data analysis for "pHLARE: A Genetically Encoded Ratiometric Lysosome pH Biosensor"

### Supplemental Methods

**pHLARE Image Analysis:** To semi-automatically segment lysosomes in the pH-insensitive mCherry channel and determine lysosome pH based on a two-point nigericin calibration from pHLARE images, we implemented an image analysis algorithm in NIS Elements software based on mCherry fluorescence intensity thresholding and morphometric manipulation of the resulting binary images to reproducibly define lysosome objects and corresponding local background regions that are then used to calculate fluorescence ratios and pH values.

We implemented this algorithm as a Macro in NIS Elements. The annotated 'pHLARE_analysis.mac' was tested in NIS Elements v5 (built 1266 and 1377). Because the NIS Elements macro commands have changed over time, it will not work in older versions without adapting the command input structure. The Macro expects a dual-wavelength ND2 document with three time frames. The first frame is the image from which lysosome pH is going to be calculated. The second frame is the high pH nigericin calibration, and the third frame the low nigericin calibration. The first wavelength tab should be named '488' (sfGFP channel), and the second wavelength tab should be named '561' (mCherry channel). The Macro walks the user through a series of steps defining background and cell regions in the mCherry channel mostly utilizing the NIS Elements 'Binary Editor'. In the following, we describe the rationale underlying the different analysis steps as well as explain required user input:

*Step 1: Define dark background area outside the cell, i.e. camera offset.* Using the 'Polygon Fill' tool in foreground mode ('FG') select a dark area that does not contain any fluorescence signal. This is only done in frame 1. Note that any mistakes in the binary editor can be corrected using background mode ('BG') to delete regions before exiting the editor. Exit editor by pressing the 'esc' key or the 'Exit Editor' button.

*Step 2: Define non-lysosome intracellular fluorescence.* Using the 'Polygon Fill' tool select one or more areas inside the cell that do not contain lysosomes and are representative of the cell membrane-bound pHLARE fluorescence background. This is only done in frame 1. Exit editor.

*Step 3: Adjust threshold (if necessary).* The algorithm sets an initial threshold at three times the pHLARE background intensity in the cell. If this seems to stringent (too many lysosomes not detected) the low threshold limit can be adjusted down a little bit. However, care should be taken for the threshold to be sufficiently stringent to exclude bright objects that are not lysosomes (i.e. overlapping cell membrane folds etc.). When satisfied, press 'OK'.

*Step 4: Define cell of interest.* Using the 'Polygon Fill' tool select the cell (or region) of interest. This needs to be done separately in all three frames because the cell (or stage) might move during acquisition and nigericin buffer changes. This step can also be used to manually exclude areas of high fluorescence that are not lysosomes. Exit editor. The algorithm next performs several morphometric operations on the thresholded binary to smooth and dilate thresholded objects sufficiently large to include all lysosome signal.

*Step 5: Remove objects touching borders.* Select 'OK'. This removes objects that touch the image edge to avoid mismatches in the number of lysosome and local background objects. Similarly, the algorithm next fills holes in lysosome objects and defines a 1-pixel wide exterior contour of each lysosome object as the corresponding local background and applies the cell mask to lysosome objects and local background objects. These steps are required to ensure that the number of lysosome and local background objects is identical and that each background corresponds to the correct lysosome.

*Step 6: Define dark background outside cell.* As above, using the ‘Polygon Fill’ tool select a dark background region in which there is no fluorescence in all three frames. Exit editor.

If the Macro finishes correctly, three binary layers are displayed in this order and selected: ‘dark_bkg’, the dark background outside the cell, ‘lyso_bkg’, local background objects around lysosomes, ‘lyso_signal’, corresponding lysosome objects. The Macro also opens the ‘Automated Measurement Results’ dialog box. To measure the fluorescence intensity in the objects, click ‘Update ND measurements’, ‘All frames’, ‘OK’. The output is a list of ‘lyso_signal’ values, followed by the same number of ‘lyso_bkg’ values, and one ‘dark_bkg’ value. This list order is repeated three times with increasing ‘ND.T’ index for the three image frames. All sorting must be turned off in ‘Automated Measurements’ to get this correct order of fluorescence measurements.

Next, the measured data are exported to an ‘Object Data’ worksheet in Excel. To simplify copying of the data into an empty pHLARE analysis worksheet, in ‘Measurement options’, ‘Data export’ only the columns included on the data analysis worksheet should be selected. After copying these columns into an empty pHLARE analysis worksheet, pHlys values are calculated automatically and are found in the purple boxes. Nigericin calibration pH values need to be entered in the green box, and pHlys calculations automatically update when these values are changed. If errors appear in the ‘ROI check’ area, exported data most likely were not copied correctly and there is a mismatch between lysosome and local background objects. If this is the case calculated pHlys will not be correct.

We tested the image analysis macro and corresponding excel worksheet extensively, but the user should be aware that we cannot anticipate all potential sources of error and cannot guarantee that this will function error-free in all circumstances and may have to be adapted for specific experimental setups.

A sample data set of control RPE pHLARE cells is included for testing purposes corresponding to ‘cell 6’ in Fig. 2. This data set has been analyzed as described above and also includes the saved binary layers, and data from this cell are also included in the ‘pHLARE_analysis_worksheet.xlsx’.

```
//tested with v5.02.00 built 1266

global double OFFS, CYTO, LYSO

ViewComponents("01");
ShowControl("ROIStatisticsControl",1);
Uncalibrate();                                         //uncalibrate
document, all size measurements are in pixels
ShowControl("BinaryLayers",1);
BinLayerDelete("*");
OverlayTransparency(70);                      //set up view and reset
previous binary layers

ImageEdit();                                           //Step 1: define
dark background, i.e. camera offset
ROIStatisticsGetData(1,2,&OFFS,NULL);
BinLayerDelete("Binary Edit");

ImageEdit();                                           //Step 2: define
cellular background
ROIStatisticsGetData(1,2,&CYTO,NULL);
BinLayerDelete("Binary Edit");

LYSO = 3*(CYTO-OFFS);                                  //Step 3: define
initial threshhold

DefineThresholdMCHOperation(1);                        //set up threshhold
parameters
DefineThresholdMCHChannel(0,16383,0,0);
DefineThresholdMCHChannel(1,LYSO,16383,0);
DefineThresholdProcessingOnChannel(1,1,1,0,0);
DefineThresholdRestrictionSizeOnChannel(1,1,3.000000,500.000000);
DefineThresholdRestrictionCircularityOnChannel(1,0,0.600000,1.000000);
_DefineThreshold();

BinLayerMergeND("", "Threshold (561), Threshold (561)", 3, 1, 1, 1);
BinLayerSelect("Subtraction");

ImageEdit();                                           //Step 4: define
cell of interest in all three frames
BinLayerMergeND("", "Subtraction, Threshold (561)", 2, 1, 1, 1);

BinLayerDuplicate("lyso_signal","Intersection");
BinLayerDelete("Subtraction");
BinLayerDelete("Intersection");

SmoothBinaryND(1,1,1,1);                      //binary operations to
define circular regions around
DilateBinaryND(3,5,1,1,1);                             //thresholded
objects
```

```
RemoveAllTouchingBinary();                              //Step 5: remove
objects that touch the image edge
BinLayerDuplicate("temp_bkg","lyso_signal");
FillHolesND(1,1,1);                                     //fill holes to
avoid interiour contours
ContourND(1,1,1);                                       //define local
background contour around lysosome objects
BinLayerMergeND("", "lyso_signal, temp_bkg", 8, 1, 1, 1);
BinLayerDelete("lyso_signal");
BinLayerDuplicate("lyso_signal","Having");
BinLayerDelete("Having");
BinLayerDuplicate("lyso_bkg","temp_bkg");
BinLayerDelete("temp_bkg");                             //cell mask applied

BinLayerMergeND("", "Threshold (561), Threshold (561)", 3, 1, 1, 1);
BinLayerSelect("Subtraction");
ImageEdit();                                            //Step 6: define
dark background in all three frames
BinLayerDuplicate("dark_bkg","Subtraction");
BinLayerDelete("Subtraction");
BinLayerDelete("Threshold (561)");

BinLayerSelect("dark_bkg,lyso_bkg,lyso_signal");
ShowControl("AutoMeasResults",1);

ResetObjectFeatures();
SelectObjectFeature("Area");
SelectObjectFeature("Circularity");
SelectObjectFeature("ShapeFactor");
SelectObjectFeature("CentreX");
SelectObjectFeature("CentreY");
SelectObjectFeature("MeanChannel1");
SelectObjectFeature("MeanChannel2");
```

| Item | Source | BinaryID | ObjID | ND.T | Area [px²] | Circularity | ShapeFacto | CentreX [px |
|---|---|---|---|---|---|---|---|---|
| 1 | sample_dat | lyso_signal | 1 | 1 | 232 | 1 | 1 | 364.5 |
| 2 | sample_dat | lyso_signal | 2 | 1 | 4788 | 0.307 | 0.698 | 341.99 |
| 3 | sample_dat | lyso_signal | 3 | 1 | 326 | 0.971 | 1 | 403.56 |
| 4 | sample_dat | lyso_signal | 4 | 1 | 319 | 0.997 | 1 | 278.41 |
| 5 | sample_dat | lyso_signal | 5 | 1 | 527 | 0.89 | 0.92 | 437.17 |
| 6 | sample_dat | lyso_signal | 6 | 1 | 1347 | 0.391 | 0.654 | 284.07 |
| 7 | sample_dat | lyso_signal | 7 | 1 | 391 | 1 | 1 | 493.43 |
| 8 | sample_dat | lyso_signal | 8 | 1 | 3329 | 0.273 | 0.614 | 457.15 |
| 9 | sample_dat | lyso_signal | 9 | 1 | 250 | 1 | 1 | 409.15 |
| 10 | sample_dat | lyso_signal | 10 | 1 | 743 | 0.646 | 0.729 | 512.08 |
| 11 | sample_dat | lyso_signal | 11 | 1 | 215 | 1 | 1 | 318 |
| 12 | sample_dat | lyso_signal | 12 | 1 | 324 | 1 | 1 | 377.1 |
| 13 | sample_dat | lyso_signal | 13 | 1 | 216 | 1 | 1 | 402.5 |
| 14 | sample_dat | lyso_signal | 14 | 1 | 396 | 0.77 | 0.861 | 586.53 |
| 15 | sample_dat | lyso_signal | 15 | 1 | 306 | 1 | 1 | 545.11 |
| 16 | sample_dat | lyso_signal | 16 | 1 | 474 | 1 | 1 | 506.43 |
| 17 | sample_dat | lyso_signal | 17 | 1 | 269 | 1 | 1 | 356.38 |
| 18 | sample_dat | lyso_signal | 18 | 1 | 276 | 1 | 1 | 618.63 |
| 19 | sample_dat | lyso_signal | 19 | 1 | 249 | 1 | 1 | 309 |
| 20 | sample_dat | lyso_signal | 20 | 1 | 8723 | 0.182 | 0.586 | 533.18 |
| 21 | sample_dat | lyso_signal | 21 | 1 | 449 | 0.94 | 0.963 | 286.33 |
| 22 | sample_dat | lyso_signal | 22 | 1 | 5997 | 0.233 | 0.576 | 633.17 |
| 23 | sample_dat | lyso_signal | 23 | 1 | 291 | 1 | 1 | 457.75 |
| 24 | sample_dat | lyso_signal | 24 | 1 | 284 | 1 | 1 | 359.5 |
| 25 | sample_dat | lyso_signal | 25 | 1 | 4557 | 0.218 | 0.622 | 441.87 |
| 26 | sample_dat | lyso_signal | 26 | 1 | 267 | 1 | 1 | 335.6 |
| 27 | sample_dat | lyso_signal | 27 | 1 | 200 | 1 | 1 | 307.5 |
| 28 | sample_dat | lyso_signal | 28 | 1 | 863 | 0.521 | 0.681 | 350.78 |
| 29 | sample_dat | lyso_signal | 29 | 1 | 244 | 1 | 1 | 391.88 |
| 30 | sample_dat | lyso_signal | 30 | 1 | 423 | 1 | 1 | 332.13 |
| 31 | sample_dat | lyso_signal | 31 | 1 | 227 | 1 | 1 | 355.38 |
| 32 | sample_dat | lyso_signal | 32 | 1 | 286 | 1 | 1 | 293.86 |
| 33 | sample_dat | lyso_signal | 33 | 1 | 442 | 0.996 | 1 | 308.22 |
| 34 | sample_dat | lyso_signal | 34 | 1 | 185 | 1 | 1 | 340 |
| 35 | sample_dat | lyso_signal | 35 | 1 | 304 | 1 | 1 | 470.62 |
| 36 | sample_dat | lyso_signal | 36 | 1 | 415 | 1 | 1 | 310.11 |
| 37 | sample_dat | lyso_signal | 37 | 1 | 458 | 1 | 1 | 538.38 |
| 38 | sample_dat | lyso_signal | 38 | 1 | 4692 | 0.209 | 0.493 | 446.3 |
| 39 | sample_dat | lyso_signal | 39 | 1 | 305 | 1 | 1 | 269.64 |
| 40 | sample_dat | lyso_signal | 40 | 1 | 2777 | 0.662 | 0.863 | 348.76 |
| 41 | sample_dat | lyso_signal | 41 | 1 | 1709 | 0.535 | 0.727 | 551.89 |
| 42 | sample_dat | lyso_signal | 42 | 1 | 172 | 1 | 1 | 396.5 |
| 43 | sample_dat | lyso_signal | 43 | 1 | 535 | 0.993 | 1 | 601.83 |
| 44 | sample_dat | lyso_signal | 44 | 1 | 239 | 1 | 1 | 290.75 |
| 45 | sample_dat | lyso_signal | 45 | 1 | 297 | 1 | 1 | 616.39 |
| 46 | sample_dat | lyso_signal | 46 | 1 | 303 | 1 | 1 | 264.83 |
| 47 | sample_dat | lyso_signal | 47 | 1 | 185 | 1 | 1 | 304 |

| | | | | | | | | |
|---|---|---|---|---|---|---|---|---|
| 48 | sample_dat | lyso_signal | 48 | 1 | 488 | 0.996 | 1 | 496.51 |
| 49 | sample_dat | lyso_signal | 49 | 1 | 1061 | 0.668 | 0.775 | 591.28 |
| 50 | sample_dat | lyso_signal | 50 | 1 | 347 | 1 | 1 | 399.03 |
| 51 | sample_dat | lyso_signal | 51 | 1 | 384 | 0.696 | 0.804 | 266.97 |
| 52 | sample_dat | lyso_signal | 52 | 1 | 291 | 1 | 1 | 318.49 |
| 53 | sample_dat | lyso_signal | 53 | 1 | 50728 | 0.142 | 0.654 | 454.51 |
| 54 | sample_dat | lyso_signal | 54 | 1 | 1102 | 0.605 | 0.694 | 277.95 |
| 55 | sample_dat | lyso_signal | 55 | 1 | 279 | 1 | 1 | 358.38 |
| 56 | sample_dat | lyso_signal | 56 | 1 | 384 | 0.967 | 0.998 | 333.72 |
| 57 | sample_dat | lyso_signal | 57 | 1 | 668 | 0.939 | 0.959 | 262.55 |
| 58 | sample_dat | lyso_signal | 58 | 1 | 880 | 0.654 | 0.772 | 607.61 |
| 59 | sample_dat | lyso_signal | 59 | 1 | 448 | 0.98 | 0.995 | 298.62 |
| 60 | sample_dat | lyso_signal | 60 | 1 | 300 | 1 | 1 | 330.5 |
| 61 | sample_dat | lyso_signal | 61 | 1 | 256 | 1 | 1 | 178.5 |
| 62 | sample_dat | lyso_signal | 62 | 1 | 297 | 1 | 1 | 199.88 |
| 63 | sample_dat | lyso_signal | 63 | 1 | 2348 | 0.35 | 0.651 | 317.1 |
| 64 | sample_dat | lyso_signal | 64 | 1 | 750 | 0.902 | 0.918 | 551.31 |
| 65 | sample_dat | lyso_signal | 65 | 1 | 522 | 1 | 1 | 581.95 |
| 66 | sample_dat | lyso_signal | 66 | 1 | 537 | 1 | 1 | 550.34 |
| 67 | sample_dat | lyso_signal | 67 | 1 | 233 | 1 | 1 | 357.14 |
| 68 | sample_dat | lyso_signal | 68 | 1 | 311 | 1 | 1 | 515.23 |
| 69 | sample_dat | lyso_signal | 69 | 1 | 587 | 0.916 | 0.968 | 386.76 |
| 70 | sample_dat | lyso_signal | 70 | 1 | 573 | 0.781 | 0.865 | 547.19 |
| 71 | sample_dat | lyso_signal | 71 | 1 | 521 | 1 | 1 | 490.9 |
| 72 | sample_dat | lyso_signal | 72 | 1 | 1742 | 0.608 | 0.79 | 432.01 |
| 73 | sample_dat | lyso_signal | 73 | 1 | 413 | 0.986 | 1 | 262.99 |
| 74 | sample_dat | lyso_signal | 74 | 1 | 449 | 1 | 1 | 371.89 |
| 75 | sample_dat | lyso_signal | 75 | 1 | 562 | 0.949 | 0.965 | 589.04 |
| 76 | sample_dat | lyso_signal | 76 | 1 | 353 | 0.856 | 0.936 | 556.45 |
| 77 | sample_dat | lyso_signal | 77 | 1 | 1210 | 0.449 | 0.698 | 312.11 |
| 78 | sample_dat | lyso_signal | 78 | 1 | 458 | 0.987 | 1 | 238.33 |
| 79 | sample_dat | lyso_signal | 79 | 1 | 6524 | 0.162 | 0.494 | 408.59 |
| 80 | sample_dat | lyso_signal | 80 | 1 | 261 | 1 | 1 | 216.89 |
| 81 | sample_dat | lyso_signal | 81 | 1 | 354 | 1 | 1 | 563.9 |
| 82 | sample_dat | lyso_signal | 82 | 1 | 204 | 1 | 1 | 588.89 |
| 83 | sample_dat | lyso_signal | 83 | 1 | 742 | 0.936 | 0.95 | 467.46 |
| 84 | sample_dat | lyso_signal | 84 | 1 | 794 | 0.674 | 0.824 | 540.28 |
| 85 | sample_dat | lyso_signal | 85 | 1 | 712 | 0.909 | 0.948 | 293.89 |
| 86 | sample_dat | lyso_signal | 86 | 1 | 216 | 1 | 1 | 576.5 |
| 87 | sample_dat | lyso_signal | 87 | 1 | 185 | 1 | 1 | 446 |
| 88 | sample_dat | lyso_signal | 88 | 1 | 256 | 1 | 1 | 494 |
| 89 | sample_dat | lyso_signal | 89 | 1 | 334 | 1 | 1 | 307.37 |
| 90 | sample_dat | lyso_signal | 90 | 1 | 2554 | 0.487 | 0.657 | 535.65 |
| 91 | sample_dat | lyso_signal | 91 | 1 | 1104 | 0.925 | 0.933 | 571.43 |
| 92 | sample_dat | lyso_signal | 92 | 1 | 200 | 1 | 1 | 279 |
| 93 | sample_dat | lyso_signal | 93 | 1 | 1384 | 0.681 | 0.814 | 453.35 |
| 94 | sample_dat | lyso_signal | 94 | 1 | 400 | 0.888 | 0.927 | 385.21 |
| 95 | sample_dat | lyso_signal | 95 | 1 | 284 | 0.937 | 0.99 | 315 |

| | | | | | | | | |
|---|---|---|---|---|---|---|---|---|
| 96 | sample_dat | lyso_signal | 96 | 1 | 216 | 1 | 1 | 351.5 |
| 97 | sample_dat | lyso_signal | 97 | 1 | 540 | 0.802 | 0.881 | 308.04 |
| 98 | sample_dat | lyso_signal | 98 | 1 | 708 | 0.876 | 0.933 | 485.55 |
| 99 | sample_dat | lyso_signal | 99 | 1 | 216 | 1 | 1 | 380.5 |
| 100 | sample_dat | lyso_signal | 100 | 1 | 1454 | 0.469 | 0.585 | 499.61 |
| 101 | sample_dat | lyso_signal | 101 | 1 | 1835 | 0.376 | 0.569 | 385.92 |
| 102 | sample_dat | lyso_signal | 102 | 1 | 274 | 1 | 1 | 451.5 |
| 103 | sample_dat | lyso_signal | 103 | 1 | 4178 | 0.25 | 0.484 | 421.02 |
| 104 | sample_dat | lyso_signal | 104 | 1 | 287 | 1 | 1 | 465.88 |
| 105 | sample_dat | lyso_signal | 105 | 1 | 287 | 1 | 1 | 391.39 |
| 106 | sample_dat | lyso_bkg | 1 | 1 | 62 | 0.082 | 0.281 | 364.5 |
| 107 | sample_dat | lyso_bkg | 2 | 1 | 540 | 0.009 | 0.079 | 342.51 |
| 108 | sample_dat | lyso_bkg | 3 | 1 | 78 | 0.068 | 0.241 | 403.5 |
| 109 | sample_dat | lyso_bkg | 4 | 1 | 74 | 0.068 | 0.239 | 278.43 |
| 110 | sample_dat | lyso_bkg | 5 | 1 | 104 | 0.049 | 0.181 | 437.15 |
| 111 | sample_dat | lyso_bkg | 6 | 1 | 252 | 0.02 | 0.122 | 285.29 |
| 112 | sample_dat | lyso_bkg | 7 | 1 | 82 | 0.061 | 0.216 | 493.46 |
| 113 | sample_dat | lyso_bkg | 8 | 1 | 482 | 0.01 | 0.089 | 456.78 |
| 114 | sample_dat | lyso_bkg | 9 | 1 | 66 | 0.08 | 0.277 | 409.09 |
| 115 | sample_dat | lyso_bkg | 10 | 1 | 148 | 0.035 | 0.145 | 512.33 |
| 116 | sample_dat | lyso_bkg | 11 | 1 | 60 | 0.086 | 0.294 | 318 |
| 117 | sample_dat | lyso_bkg | 12 | 1 | 74 | 0.068 | 0.237 | 377.05 |
| 118 | sample_dat | lyso_bkg | 13 | 1 | 60 | 0.086 | 0.294 | 402.5 |
| 119 | sample_dat | lyso_bkg | 14 | 1 | 96 | 0.053 | 0.209 | 586.58 |
| 120 | sample_dat | lyso_bkg | 15 | 1 | 72 | 0.071 | 0.244 | 545.07 |
| 121 | sample_dat | lyso_bkg | 16 | 1 | 90 | 0.055 | 0.194 | 506.46 |
| 122 | sample_dat | lyso_bkg | 17 | 1 | 68 | 0.077 | 0.264 | 356.43 |
| 123 | sample_dat | lyso_bkg | 18 | 1 | 70 | 0.077 | 0.264 | 618.59 |
| 124 | sample_dat | lyso_bkg | 19 | 1 | 64 | 0.078 | 0.27 | 309 |
| 125 | sample_dat | lyso_bkg | 20 | 1 | 972 | 0.005 | 0.065 | 530.36 |
| 126 | sample_dat | lyso_bkg | 21 | 1 | 90 | 0.054 | 0.193 | 286.54 |
| 127 | sample_dat | lyso_bkg | 22 | 1 | 652 | 0.008 | 0.063 | 630.98 |
| 128 | sample_dat | lyso_bkg | 23 | 1 | 70 | 0.072 | 0.249 | 457.84 |
| 129 | sample_dat | lyso_bkg | 24 | 1 | 68 | 0.072 | 0.249 | 359.5 |
| 130 | sample_dat | lyso_bkg | 25 | 1 | 550 | 0.009 | 0.075 | 439.82 |
| 131 | sample_dat | lyso_bkg | 26 | 1 | 68 | 0.077 | 0.268 | 335.57 |
| 132 | sample_dat | lyso_bkg | 27 | 1 | 58 | 0.09 | 0.308 | 307.5 |
| 133 | sample_dat | lyso_bkg | 28 | 1 | 178 | 0.029 | 0.14 | 350.44 |
| 134 | sample_dat | lyso_bkg | 29 | 1 | 64 | 0.08 | 0.275 | 391.92 |
| 135 | sample_dat | lyso_bkg | 30 | 1 | 86 | 0.059 | 0.207 | 332.26 |
| 136 | sample_dat | lyso_bkg | 31 | 1 | 62 | 0.084 | 0.288 | 355.42 |
| 137 | sample_dat | lyso_bkg | 32 | 1 | 70 | 0.073 | 0.255 | 293.91 |
| 138 | sample_dat | lyso_bkg | 33 | 1 | 90 | 0.058 | 0.206 | 308.18 |
| 139 | sample_dat | lyso_bkg | 34 | 1 | 56 | 0.095 | 0.324 | 340 |
| 140 | sample_dat | lyso_bkg | 35 | 1 | 72 | 0.071 | 0.244 | 470.58 |
| 141 | sample_dat | lyso_bkg | 36 | 1 | 84 | 0.059 | 0.207 | 310.07 |
| 142 | sample_dat | lyso_bkg | 37 | 1 | 88 | 0.055 | 0.197 | 538.43 |
| 143 | sample_dat | lyso_bkg | 38 | 1 | 638 | 0.008 | 0.067 | 447.26 |

| | | | | | | | |
|---|---|---|---|---|---|---|---|
| 144 | sample_dat lyso_bkg | 39 | 1 | 72 | 0.071 | 0.244 | 269.78 |
| 145 | sample_dat lyso_bkg | 40 | 1 | 276 | 0.019 | 0.086 | 347.75 |
| 146 | sample_dat lyso_bkg | 41 | 1 | 246 | 0.02 | 0.105 | 552.98 |
| 147 | sample_dat lyso_bkg | 42 | 1 | 52 | 0.095 | 0.323 | 396.5 |
| 148 | sample_dat lyso_bkg | 43 | 1 | 98 | 0.051 | 0.184 | 601.5 |
| 149 | sample_dat lyso_bkg | 44 | 1 | 64 | 0.082 | 0.281 | 290.84 |
| 150 | sample_dat lyso_bkg | 45 | 1 | 70 | 0.071 | 0.244 | 616.43 |
| 151 | sample_dat lyso_bkg | 46 | 1 | 72 | 0.071 | 0.247 | 264.89 |
| 152 | sample_dat lyso_bkg | 47 | 1 | 56 | 0.095 | 0.324 | 304 |
| 153 | sample_dat lyso_bkg | 48 | 1 | 94 | 0.055 | 0.194 | 496.5 |
| 154 | sample_dat lyso_bkg | 49 | 1 | 172 | 0.029 | 0.126 | 591.83 |
| 155 | sample_dat lyso_bkg | 50 | 1 | 78 | 0.067 | 0.232 | 399.01 |
| 156 | sample_dat lyso_bkg | 51 | 1 | 102 | 0.052 | 0.213 | 266.98 |
| 157 | sample_dat lyso_bkg | 52 | 1 | 70 | 0.072 | 0.25 | 318.49 |
| 158 | sample_dat lyso_bkg | 53 | 1 | 2380 | 0.002 | 0.031 | 448.6 |
| 159 | sample_dat lyso_bkg | 54 | 1 | 186 | 0.027 | 0.117 | 279 |
| 160 | sample_dat lyso_bkg | 55 | 1 | 68 | 0.073 | 0.254 | 358.43 |
| 161 | sample_dat lyso_bkg | 56 | 1 | 84 | 0.061 | 0.218 | 333.89 |
| 162 | sample_dat lyso_bkg | 57 | 1 | 118 | 0.046 | 0.169 | 262.47 |
| 163 | sample_dat lyso_bkg | 58 | 1 | 156 | 0.031 | 0.137 | 608.71 |
| 164 | sample_dat lyso_bkg | 59 | 1 | 90 | 0.056 | 0.2 | 298.4 |
| 165 | sample_dat lyso_bkg | 60 | 1 | 70 | 0.069 | 0.239 | 330.5 |
| 166 | sample_dat lyso_bkg | 61 | 1 | 66 | 0.078 | 0.27 | 178.5 |
| 167 | sample_dat lyso_bkg | 62 | 1 | 70 | 0.071 | 0.244 | 199.93 |
| 168 | sample_dat lyso_bkg | 63 | 1 | 368 | 0.014 | 0.102 | 317.06 |
| 169 | sample_dat lyso_bkg | 64 | 1 | 122 | 0.04 | 0.149 | 551.21 |
| 170 | sample_dat lyso_bkg | 65 | 1 | 98 | 0.054 | 0.194 | 581.96 |
| 171 | sample_dat lyso_bkg | 66 | 1 | 98 | 0.052 | 0.187 | 550.21 |
| 172 | sample_dat lyso_bkg | 67 | 1 | 64 | 0.084 | 0.288 | 357.09 |
| 173 | sample_dat lyso_bkg | 68 | 1 | 72 | 0.069 | 0.24 | 515.14 |
| 174 | sample_dat lyso_bkg | 69 | 1 | 110 | 0.048 | 0.181 | 387.15 |
| 175 | sample_dat lyso_bkg | 70 | 1 | 116 | 0.044 | 0.175 | 547.13 |
| 176 | sample_dat lyso_bkg | 71 | 1 | 96 | 0.052 | 0.185 | 490.78 |
| 177 | sample_dat lyso_bkg | 72 | 1 | 236 | 0.022 | 0.107 | 431.51 |
| 178 | sample_dat lyso_bkg | 73 | 1 | 86 | 0.059 | 0.209 | 262.97 |
| 179 | sample_dat lyso_bkg | 74 | 1 | 88 | 0.057 | 0.201 | 371.93 |
| 180 | sample_dat lyso_bkg | 75 | 1 | 104 | 0.049 | 0.179 | 588.9 |
| 181 | sample_dat lyso_bkg | 76 | 1 | 86 | 0.06 | 0.228 | 556.47 |
| 182 | sample_dat lyso_bkg | 77 | 1 | 224 | 0.022 | 0.129 | 310.76 |
| 183 | sample_dat lyso_bkg | 78 | 1 | 90 | 0.055 | 0.198 | 238.39 |
| 184 | sample_dat lyso_bkg | 79 | 1 | 890 | 0.006 | 0.067 | 408.46 |
| 185 | sample_dat lyso_bkg | 80 | 1 | 66 | 0.077 | 0.264 | 216.92 |
| 186 | sample_dat lyso_bkg | 81 | 1 | 76 | 0.063 | 0.219 | 563.93 |
| 187 | sample_dat lyso_bkg | 82 | 1 | 58 | 0.088 | 0.301 | 588.93 |
| 188 | sample_dat lyso_bkg | 83 | 1 | 120 | 0.042 | 0.154 | 467.45 |
| 189 | sample_dat lyso_bkg | 84 | 1 | 146 | 0.034 | 0.151 | 539.29 |
| 190 | sample_dat lyso_bkg | 85 | 1 | 120 | 0.042 | 0.16 | 294.24 |
| 191 | sample_dat lyso_bkg | 86 | 1 | 60 | 0.086 | 0.294 | 576.5 |

| | | | | | | | | |
|---|---|---|---|---|---|---|---|---|
| 192 | sample_dat | lyso_bkg | 87 | 1 | 56 | 0.095 | 0.324 | 446 |
| 193 | sample_dat | lyso_bkg | 88 | 1 | 66 | 0.078 | 0.27 | 494 |
| 194 | sample_dat | lyso_bkg | 89 | 1 | 76 | 0.068 | 0.235 | 307.42 |
| 195 | sample_dat | lyso_bkg | 90 | 1 | 316 | 0.016 | 0.081 | 535.43 |
| 196 | sample_dat | lyso_bkg | 91 | 1 | 148 | 0.034 | 0.125 | 571.49 |
| 197 | sample_dat | lyso_bkg | 92 | 1 | 58 | 0.09 | 0.308 | 279 |
| 198 | sample_dat | lyso_bkg | 93 | 1 | 198 | 0.026 | 0.116 | 452.58 |
| 199 | sample_dat | lyso_bkg | 94 | 1 | 90 | 0.057 | 0.208 | 385.16 |
| 200 | sample_dat | lyso_bkg | 95 | 1 | 74 | 0.072 | 0.258 | 315 |
| 201 | sample_dat | lyso_bkg | 96 | 1 | 60 | 0.086 | 0.294 | 351.5 |
| 202 | sample_dat | lyso_bkg | 97 | 1 | 110 | 0.046 | 0.18 | 307.93 |
| 203 | sample_dat | lyso_bkg | 98 | 1 | 124 | 0.042 | 0.163 | 485.73 |
| 204 | sample_dat | lyso_bkg | 99 | 1 | 60 | 0.086 | 0.294 | 380.5 |
| 205 | sample_dat | lyso_bkg | 100 | 1 | 240 | 0.02 | 0.097 | 500.13 |
| 206 | sample_dat | lyso_bkg | 101 | 1 | 306 | 0.016 | 0.095 | 384.05 |
| 207 | sample_dat | lyso_bkg | 102 | 1 | 68 | 0.075 | 0.259 | 451.5 |
| 208 | sample_dat | lyso_bkg | 103 | 1 | 560 | 0.009 | 0.065 | 423.45 |
| 209 | sample_dat | lyso_bkg | 104 | 1 | 70 | 0.073 | 0.254 | 465.93 |
| 210 | sample_dat | lyso_bkg | 105 | 1 | 70 | 0.073 | 0.254 | 391.43 |
| 211 | sample_dat | dark_bkg | 1 | 1 | 18627 | 0.545 | 0.506 | 88.39 |
| 212 | sample_dat | lyso_signal | 1 | 2 | 266 | 1 | 1 | 374.5 |
| 213 | sample_dat | lyso_signal | 2 | 2 | 833 | 0.798 | 0.868 | 361.73 |
| 214 | sample_dat | lyso_signal | 3 | 2 | 323 | 1 | 1 | 390.37 |
| 215 | sample_dat | lyso_signal | 4 | 2 | 875 | 0.681 | 0.798 | 410.37 |
| 216 | sample_dat | lyso_signal | 5 | 2 | 3881 | 0.505 | 0.827 | 348.86 |
| 217 | sample_dat | lyso_signal | 6 | 2 | 563 | 0.782 | 0.865 | 303.86 |
| 218 | sample_dat | lyso_signal | 7 | 2 | 526 | 0.729 | 0.821 | 402.33 |
| 219 | sample_dat | lyso_signal | 8 | 2 | 400 | 1 | 1 | 438.04 |
| 220 | sample_dat | lyso_signal | 9 | 2 | 2161 | 0.417 | 0.588 | 401.88 |
| 221 | sample_dat | lyso_signal | 10 | 2 | 877 | 0.77 | 0.822 | 304.55 |
| 222 | sample_dat | lyso_signal | 11 | 2 | 551 | 0.9 | 0.916 | 448.22 |
| 223 | sample_dat | lyso_signal | 12 | 2 | 323 | 0.985 | 1 | 334.49 |
| 224 | sample_dat | lyso_signal | 13 | 2 | 564 | 0.824 | 0.887 | 478.96 |
| 225 | sample_dat | lyso_signal | 14 | 2 | 200 | 1 | 1 | 457.5 |
| 226 | sample_dat | lyso_signal | 15 | 2 | 442 | 1 | 1 | 504.98 |
| 227 | sample_dat | lyso_signal | 16 | 2 | 200 | 1 | 1 | 304.5 |
| 228 | sample_dat | lyso_signal | 17 | 2 | 189 | 1 | 1 | 340.88 |
| 229 | sample_dat | lyso_signal | 18 | 2 | 648 | 0.878 | 0.909 | 403.73 |
| 230 | sample_dat | lyso_signal | 19 | 2 | 445 | 0.811 | 0.881 | 304.56 |
| 231 | sample_dat | lyso_signal | 20 | 2 | 215 | 1 | 1 | 263 |
| 232 | sample_dat | lyso_signal | 21 | 2 | 725 | 0.982 | 0.985 | 465.77 |
| 233 | sample_dat | lyso_signal | 22 | 2 | 344 | 1 | 1 | 285.36 |
| 234 | sample_dat | lyso_signal | 23 | 2 | 232 | 1 | 1 | 330 |
| 235 | sample_dat | lyso_signal | 24 | 2 | 491 | 0.823 | 0.88 | 528.97 |
| 236 | sample_dat | lyso_signal | 25 | 2 | 581 | 1 | 1 | 498.62 |
| 237 | sample_dat | lyso_signal | 26 | 2 | 306 | 1 | 1 | 449.89 |
| 238 | sample_dat | lyso_signal | 27 | 2 | 410 | 1 | 1 | 306.4 |
| 239 | sample_dat | lyso_signal | 28 | 2 | 616 | 0.983 | 0.997 | 407.03 |

| | | | | | | | | |
|---|---|---|---|---|---|---|---|---|
| 240 | sample_dat | lyso_signal | 29 | 2 | 410 | 1 | 1 | 336.45 |
| 241 | sample_dat | lyso_signal | 30 | 2 | 5587 | 0.173 | 0.452 | 376.28 |
| 242 | sample_dat | lyso_signal | 31 | 2 | 1008 | 0.788 | 0.847 | 524.03 |
| 243 | sample_dat | lyso_signal | 32 | 2 | 672 | 0.97 | 0.976 | 317.21 |
| 244 | sample_dat | lyso_signal | 33 | 2 | 249 | 1 | 1 | 265 |
| 245 | sample_dat | lyso_signal | 34 | 2 | 541 | 0.794 | 0.849 | 473.46 |
| 246 | sample_dat | lyso_signal | 35 | 2 | 215 | 1 | 1 | 550 |
| 247 | sample_dat | lyso_signal | 36 | 2 | 938 | 0.856 | 0.89 | 616.58 |
| 248 | sample_dat | lyso_signal | 37 | 2 | 249 | 1 | 1 | 491 |
| 249 | sample_dat | lyso_signal | 38 | 2 | 232 | 1 | 1 | 291.5 |
| 250 | sample_dat | lyso_signal | 39 | 2 | 489 | 0.852 | 0.9 | 320.04 |
| 251 | sample_dat | lyso_signal | 40 | 2 | 1879 | 0.573 | 0.663 | 575.24 |
| 252 | sample_dat | lyso_signal | 41 | 2 | 2758 | 0.505 | 0.696 | 430.61 |
| 253 | sample_dat | lyso_signal | 42 | 2 | 239 | 1 | 1 | 245.75 |
| 254 | sample_dat | lyso_signal | 43 | 2 | 324 | 1 | 1 | 309.12 |
| 255 | sample_dat | lyso_signal | 44 | 2 | 249 | 1 | 1 | 283 |
| 256 | sample_dat | lyso_signal | 45 | 2 | 583 | 0.979 | 0.992 | 496.92 |
| 257 | sample_dat | lyso_signal | 46 | 2 | 287 | 1 | 1 | 644.12 |
| 258 | sample_dat | lyso_signal | 47 | 2 | 903 | 0.686 | 0.775 | 571.13 |
| 259 | sample_dat | lyso_signal | 48 | 2 | 200 | 1 | 1 | 251.5 |
| 260 | sample_dat | lyso_signal | 49 | 2 | 565 | 0.663 | 0.804 | 413.16 |
| 261 | sample_dat | lyso_signal | 50 | 2 | 185 | 1 | 1 | 443 |
| 262 | sample_dat | lyso_signal | 51 | 2 | 463 | 1 | 1 | 609.99 |
| 263 | sample_dat | lyso_signal | 52 | 2 | 432 | 0.894 | 0.933 | 336.13 |
| 264 | sample_dat | lyso_signal | 53 | 2 | 425 | 0.978 | 1 | 459.5 |
| 265 | sample_dat | lyso_signal | 54 | 2 | 387 | 0.959 | 0.992 | 371.1 |
| 266 | sample_dat | lyso_signal | 55 | 2 | 39062 | 0.129 | 0.666 | 467.53 |
| 267 | sample_dat | lyso_signal | 56 | 2 | 767 | 0.769 | 0.853 | 606.21 |
| 268 | sample_dat | lyso_signal | 57 | 2 | 568 | 0.906 | 0.935 | 283.37 |
| 269 | sample_dat | lyso_signal | 58 | 2 | 239 | 1 | 1 | 611 |
| 270 | sample_dat | lyso_signal | 59 | 2 | 662 | 0.853 | 0.903 | 587.44 |
| 271 | sample_dat | lyso_signal | 60 | 2 | 456 | 0.963 | 0.991 | 323.57 |
| 272 | sample_dat | lyso_signal | 61 | 2 | 567 | 1 | 1 | 351.37 |
| 273 | sample_dat | lyso_signal | 62 | 2 | 524 | 0.942 | 0.962 | 260.02 |
| 274 | sample_dat | lyso_signal | 63 | 2 | 185 | 1 | 1 | 211 |
| 275 | sample_dat | lyso_signal | 64 | 2 | 304 | 1 | 1 | 296.58 |
| 276 | sample_dat | lyso_signal | 65 | 2 | 249 | 1 | 1 | 194 |
| 277 | sample_dat | lyso_signal | 66 | 2 | 629 | 0.88 | 0.892 | 355.65 |
| 278 | sample_dat | lyso_signal | 67 | 2 | 325 | 1 | 1 | 302.38 |
| 279 | sample_dat | lyso_signal | 68 | 2 | 338 | 1 | 1 | 217.5 |
| 280 | sample_dat | lyso_signal | 69 | 2 | 636 | 0.661 | 0.788 | 263.97 |
| 281 | sample_dat | lyso_signal | 70 | 2 | 284 | 1 | 1 | 287.5 |
| 282 | sample_dat | lyso_signal | 71 | 2 | 450 | 1 | 1 | 572.9 |
| 283 | sample_dat | lyso_signal | 72 | 2 | 302 | 1 | 1 | 349 |
| 284 | sample_dat | lyso_signal | 73 | 2 | 1205 | 0.851 | 0.88 | 566.78 |
| 285 | sample_dat | lyso_signal | 74 | 2 | 821 | 0.714 | 0.802 | 289.51 |
| 286 | sample_dat | lyso_signal | 75 | 2 | 1210 | 0.617 | 0.756 | 287.67 |
| 287 | sample_dat | lyso_signal | 76 | 2 | 1041 | 0.626 | 0.737 | 359.99 |

| | | | | | | | | |
|---|---|---|---|---|---|---|---|---|
| 288 | sample_dat | lyso_signal | 77 | 2 | 1417 | 0.481 | 0.73 | 513.68 |
| 289 | sample_dat | lyso_signal | 78 | 2 | 1594 | 0.699 | 0.824 | 416.67 |
| 290 | sample_dat | lyso_signal | 79 | 2 | 624 | 0.961 | 0.979 | 376.68 |
| 291 | sample_dat | lyso_signal | 80 | 2 | 701 | 0.936 | 0.949 | 460.17 |
| 292 | sample_dat | lyso_signal | 81 | 2 | 649 | 0.797 | 0.855 | 342.14 |
| 293 | sample_dat | lyso_signal | 82 | 2 | 244 | 1 | 1 | 532.12 |
| 294 | sample_dat | lyso_signal | 83 | 2 | 401 | 1 | 1 | 380.63 |
| 295 | sample_dat | lyso_signal | 84 | 2 | 476 | 0.856 | 0.927 | 332.22 |
| 296 | sample_dat | lyso_signal | 85 | 2 | 644 | 0.762 | 0.849 | 243.49 |
| 297 | sample_dat | lyso_signal | 86 | 2 | 244 | 1 | 1 | 430.88 |
| 298 | sample_dat | lyso_signal | 87 | 2 | 286 | 1 | 1 | 455.89 |
| 299 | sample_dat | lyso_signal | 88 | 2 | 337 | 1 | 1 | 344.76 |
| 300 | sample_dat | lyso_signal | 89 | 2 | 1700 | 0.898 | 0.937 | 418.26 |
| 301 | sample_dat | lyso_signal | 90 | 2 | 1381 | 0.507 | 0.726 | 407.86 |
| 302 | sample_dat | lyso_signal | 91 | 2 | 216 | 1 | 1 | 377.5 |
| 303 | sample_dat | lyso_signal | 92 | 2 | 483 | 0.922 | 0.95 | 293.73 |
| 304 | sample_dat | lyso_signal | 93 | 2 | 314 | 1 | 1 | 340.37 |
| 305 | sample_dat | lyso_signal | 94 | 2 | 634 | 0.775 | 0.879 | 306.41 |
| 306 | sample_dat | lyso_signal | 95 | 2 | 338 | 1 | 1 | 404.5 |
| 307 | sample_dat | lyso_signal | 96 | 2 | 352 | 1 | 1 | 364.62 |
| 308 | sample_dat | lyso_signal | 97 | 2 | 604 | 0.908 | 0.925 | 373.15 |
| 309 | sample_dat | lyso_signal | 98 | 2 | 254 | 1 | 1 | 402.78 |
| 310 | sample_dat | lyso_signal | 99 | 2 | 1519 | 0.622 | 0.756 | 400.42 |
| 311 | sample_dat | lyso_signal | 100 | 2 | 908 | 0.778 | 0.813 | 453.88 |
| 312 | sample_dat | lyso_signal | 101 | 2 | 274 | 1 | 1 | 402.5 |
| 313 | sample_dat | lyso_bkg | 1 | 2 | 66 | 0.075 | 0.259 | 374.5 |
| 314 | sample_dat | lyso_bkg | 2 | 2 | 142 | 0.037 | 0.148 | 361.52 |
| 315 | sample_dat | lyso_bkg | 3 | 2 | 74 | 0.068 | 0.237 | 390.23 |
| 316 | sample_dat | lyso_bkg | 4 | 2 | 154 | 0.032 | 0.141 | 410.46 |
| 317 | sample_dat | lyso_bkg | 5 | 2 | 386 | 0.013 | 0.082 | 351.59 |
| 318 | sample_dat | lyso_bkg | 6 | 2 | 118 | 0.046 | 0.181 | 303.46 |
| 319 | sample_dat | lyso_bkg | 7 | 2 | 114 | 0.044 | 0.178 | 402.3 |
| 320 | sample_dat | lyso_bkg | 8 | 2 | 82 | 0.06 | 0.211 | 438.02 |
| 321 | sample_dat | lyso_bkg | 9 | 2 | 312 | 0.016 | 0.085 | 400.38 |
| 322 | sample_dat | lyso_bkg | 10 | 2 | 148 | 0.035 | 0.139 | 304.66 |
| 323 | sample_dat | lyso_bkg | 11 | 2 | 102 | 0.047 | 0.17 | 448.31 |
| 324 | sample_dat | lyso_bkg | 12 | 2 | 76 | 0.068 | 0.24 | 334.51 |
| 325 | sample_dat | lyso_bkg | 13 | 2 | 112 | 0.046 | 0.176 | 478.66 |
| 326 | sample_dat | lyso_bkg | 14 | 2 | 58 | 0.09 | 0.308 | 457.5 |
| 327 | sample_dat | lyso_bkg | 15 | 2 | 88 | 0.058 | 0.205 | 504.99 |
| 328 | sample_dat | lyso_bkg | 16 | 2 | 58 | 0.09 | 0.308 | 304.5 |
| 329 | sample_dat | lyso_bkg | 17 | 2 | 56 | 0.092 | 0.316 | 340.93 |
| 330 | sample_dat | lyso_bkg | 18 | 2 | 114 | 0.043 | 0.16 | 403.61 |
| 331 | sample_dat | lyso_bkg | 19 | 2 | 100 | 0.051 | 0.198 | 304.53 |
| 332 | sample_dat | lyso_bkg | 20 | 2 | 60 | 0.086 | 0.294 | 263 |
| 333 | sample_dat | lyso_bkg | 21 | 2 | 114 | 0.043 | 0.155 | 465.65 |
| 334 | sample_dat | lyso_bkg | 22 | 2 | 76 | 0.065 | 0.228 | 285.42 |
| 335 | sample_dat | lyso_bkg | 23 | 2 | 62 | 0.082 | 0.281 | 330 |

| | | | | | | | |
|---|---|---|---|---|---|---|---|
| 336 | sample_dat lyso_bkg | 24 | 2 | 102 | 0.048 | 0.183 | 528.88 |
| 337 | sample_dat lyso_bkg | 25 | 2 | 100 | 0.049 | 0.174 | 498.57 |
| 338 | sample_dat lyso_bkg | 26 | 2 | 72 | 0.071 | 0.244 | 449.93 |
| 339 | sample_dat lyso_bkg | 27 | 2 | 84 | 0.059 | 0.208 | 306.45 |
| 340 | sample_dat lyso_bkg | 28 | 2 | 106 | 0.047 | 0.172 | 407.02 |
| 341 | sample_dat lyso_bkg | 29 | 2 | 84 | 0.06 | 0.21 | 336.48 |
| 342 | sample_dat lyso_bkg | 30 | 2 | 792 | 0.006 | 0.064 | 378.05 |
| 343 | sample_dat lyso_bkg | 31 | 2 | 152 | 0.032 | 0.128 | 524.95 |
| 344 | sample_dat lyso_bkg | 32 | 2 | 112 | 0.045 | 0.163 | 317.37 |
| 345 | sample_dat lyso_bkg | 33 | 2 | 64 | 0.078 | 0.27 | 265 |
| 346 | sample_dat lyso_bkg | 34 | 2 | 110 | 0.045 | 0.173 | 473.61 |
| 347 | sample_dat lyso_bkg | 35 | 2 | 60 | 0.086 | 0.294 | 550 |
| 348 | sample_dat lyso_bkg | 36 | 2 | 142 | 0.035 | 0.135 | 616.85 |
| 349 | sample_dat lyso_bkg | 37 | 2 | 64 | 0.078 | 0.27 | 491 |
| 350 | sample_dat lyso_bkg | 38 | 2 | 62 | 0.082 | 0.281 | 291.5 |
| 351 | sample_dat lyso_bkg | 39 | 2 | 102 | 0.05 | 0.188 | 320.13 |
| 352 | sample_dat lyso_bkg | 40 | 2 | 250 | 0.02 | 0.088 | 575.1 |
| 353 | sample_dat lyso_bkg | 41 | 2 | 324 | 0.015 | 0.082 | 431.41 |
| 354 | sample_dat lyso_bkg | 42 | 2 | 64 | 0.082 | 0.281 | 245.84 |
| 355 | sample_dat lyso_bkg | 43 | 2 | 74 | 0.068 | 0.235 | 309.08 |
| 356 | sample_dat lyso_bkg | 44 | 2 | 64 | 0.078 | 0.27 | 283 |
| 357 | sample_dat lyso_bkg | 45 | 2 | 106 | 0.05 | 0.18 | 496.92 |
| 358 | sample_dat lyso_bkg | 46 | 2 | 70 | 0.073 | 0.254 | 644.07 |
| 359 | sample_dat lyso_bkg | 47 | 2 | 158 | 0.032 | 0.136 | 571.15 |
| 360 | sample_dat lyso_bkg | 48 | 2 | 58 | 0.09 | 0.308 | 251.5 |
| 361 | sample_dat lyso_bkg | 49 | 2 | 128 | 0.041 | 0.182 | 412.51 |
| 362 | sample_dat lyso_bkg | 50 | 2 | 56 | 0.095 | 0.324 | 443 |
| 363 | sample_dat lyso_bkg | 51 | 2 | 90 | 0.056 | 0.2 | 609.99 |
| 364 | sample_dat lyso_bkg | 52 | 2 | 94 | 0.055 | 0.203 | 336.17 |
| 365 | sample_dat lyso_bkg | 53 | 2 | 88 | 0.058 | 0.21 | 459.51 |
| 366 | sample_dat lyso_bkg | 54 | 2 | 84 | 0.06 | 0.215 | 371.23 |
| 367 | sample_dat lyso_bkg | 55 | 2 | 2276 | 0.002 | 0.039 | 475.28 |
| 368 | sample_dat lyso_bkg | 56 | 2 | 134 | 0.037 | 0.149 | 606.25 |
| 369 | sample_dat lyso_bkg | 57 | 2 | 106 | 0.047 | 0.174 | 283.62 |
| 370 | sample_dat lyso_bkg | 58 | 2 | 64 | 0.082 | 0.281 | 611 |
| 371 | sample_dat lyso_bkg | 59 | 2 | 116 | 0.041 | 0.158 | 586.92 |
| 372 | sample_dat lyso_bkg | 60 | 2 | 92 | 0.055 | 0.2 | 323.91 |
| 373 | sample_dat lyso_bkg | 61 | 2 | 102 | 0.051 | 0.181 | 351.47 |
| 374 | sample_dat lyso_bkg | 62 | 2 | 100 | 0.051 | 0.184 | 259.78 |
| 375 | sample_dat lyso_bkg | 63 | 2 | 56 | 0.095 | 0.324 | 211 |
| 376 | sample_dat lyso_bkg | 64 | 2 | 72 | 0.071 | 0.246 | 296.56 |
| 377 | sample_dat lyso_bkg | 65 | 2 | 64 | 0.078 | 0.27 | 194 |
| 378 | sample_dat lyso_bkg | 66 | 2 | 110 | 0.043 | 0.156 | 355.72 |
| 379 | sample_dat lyso_bkg | 67 | 2 | 74 | 0.068 | 0.235 | 302.43 |
| 380 | sample_dat lyso_bkg | 68 | 2 | 76 | 0.067 | 0.234 | 217.5 |
| 381 | sample_dat lyso_bkg | 69 | 2 | 136 | 0.039 | 0.168 | 263.93 |
| 382 | sample_dat lyso_bkg | 70 | 2 | 68 | 0.072 | 0.249 | 287.5 |
| 383 | sample_dat lyso_bkg | 71 | 2 | 88 | 0.057 | 0.2 | 572.94 |

| | | | | | | | | |
|---|---|---|---|---|---|---|---|---|
| 384 | sample_dat | lyso_bkg | 72 | 2 | 70 | 0.069 | 0.239 | 349 |
| 385 | sample_dat | lyso_bkg | 73 | 2 | 166 | 0.032 | 0.121 | 566.87 |
| 386 | sample_dat | lyso_bkg | 74 | 2 | 148 | 0.035 | 0.145 | 290.01 |
| 387 | sample_dat | lyso_bkg | 75 | 2 | 192 | 0.026 | 0.12 | 287.74 |
| 388 | sample_dat | lyso_bkg | 76 | 2 | 176 | 0.028 | 0.125 | 359.9 |
| 389 | sample_dat | lyso_bkg | 77 | 2 | 234 | 0.021 | 0.12 | 513.36 |
| 390 | sample_dat | lyso_bkg | 78 | 2 | 204 | 0.024 | 0.105 | 417.81 |
| 391 | sample_dat | lyso_bkg | 79 | 2 | 110 | 0.047 | 0.173 | 376.73 |
| 392 | sample_dat | lyso_bkg | 80 | 2 | 120 | 0.044 | 0.163 | 460.03 |
| 393 | sample_dat | lyso_bkg | 81 | 2 | 122 | 0.041 | 0.161 | 341.84 |
| 394 | sample_dat | lyso_bkg | 82 | 2 | 64 | 0.08 | 0.275 | 532.08 |
| 395 | sample_dat | lyso_bkg | 83 | 2 | 84 | 0.061 | 0.216 | 380.58 |
| 396 | sample_dat | lyso_bkg | 84 | 2 | 100 | 0.051 | 0.195 | 332.07 |
| 397 | sample_dat | lyso_bkg | 85 | 2 | 124 | 0.04 | 0.163 | 243.15 |
| 398 | sample_dat | lyso_bkg | 86 | 2 | 64 | 0.08 | 0.275 | 430.92 |
| 399 | sample_dat | lyso_bkg | 87 | 2 | 70 | 0.073 | 0.255 | 455.94 |
| 400 | sample_dat | lyso_bkg | 88 | 2 | 76 | 0.067 | 0.232 | 344.87 |
| 401 | sample_dat | lyso_bkg | 89 | 2 | 188 | 0.026 | 0.104 | 418.57 |
| 402 | sample_dat | lyso_bkg | 90 | 2 | 224 | 0.023 | 0.118 | 407.31 |
| 403 | sample_dat | lyso_bkg | 91 | 2 | 60 | 0.086 | 0.294 | 377.5 |
| 404 | sample_dat | lyso_bkg | 92 | 2 | 98 | 0.053 | 0.193 | 293.61 |
| 405 | sample_dat | lyso_bkg | 93 | 2 | 74 | 0.071 | 0.244 | 340.42 |
| 406 | sample_dat | lyso_bkg | 94 | 2 | 124 | 0.042 | 0.172 | 306.9 |
| 407 | sample_dat | lyso_bkg | 95 | 2 | 76 | 0.067 | 0.234 | 404.5 |
| 408 | sample_dat | lyso_bkg | 96 | 2 | 78 | 0.065 | 0.228 | 364.58 |
| 409 | sample_dat | lyso_bkg | 97 | 2 | 110 | 0.046 | 0.169 | 373.41 |
| 410 | sample_dat | lyso_bkg | 98 | 2 | 66 | 0.078 | 0.271 | 402.68 |
| 411 | sample_dat | lyso_bkg | 99 | 2 | 216 | 0.023 | 0.107 | 399.94 |
| 412 | sample_dat | lyso_bkg | 100 | 2 | 146 | 0.034 | 0.131 | 453.66 |
| 413 | sample_dat | lyso_bkg | 101 | 2 | 68 | 0.075 | 0.259 | 402.5 |
| 414 | sample_dat | dark_bkg | 1 | 2 | 26142 | 0.53 | 0.508 | 79.63 |
| 415 | sample_dat | lyso_signal | 1 | 3 | 428 | 0.873 | 0.911 | 345.43 |
| 416 | sample_dat | lyso_signal | 2 | 3 | 551 | 0.935 | 0.958 | 412.27 |
| 417 | sample_dat | lyso_signal | 3 | 3 | 232 | 1 | 1 | 440 |
| 418 | sample_dat | lyso_signal | 4 | 3 | 314 | 1 | 1 | 391.35 |
| 419 | sample_dat | lyso_signal | 5 | 3 | 200 | 1 | 1 | 490.5 |
| 420 | sample_dat | lyso_signal | 6 | 3 | 965 | 0.664 | 0.776 | 449.34 |
| 421 | sample_dat | lyso_signal | 7 | 3 | 323 | 1 | 1 | 411.64 |
| 422 | sample_dat | lyso_signal | 8 | 3 | 386 | 0.965 | 0.996 | 484.09 |
| 423 | sample_dat | lyso_signal | 9 | 3 | 242 | 1 | 1 | 435.35 |
| 424 | sample_dat | lyso_signal | 10 | 3 | 227 | 1 | 1 | 304.13 |
| 425 | sample_dat | lyso_signal | 11 | 3 | 446 | 1 | 1 | 395.81 |
| 426 | sample_dat | lyso_signal | 12 | 3 | 211 | 1 | 1 | 514.63 |
| 427 | sample_dat | lyso_signal | 13 | 3 | 339 | 0.999 | 1 | 331.65 |
| 428 | sample_dat | lyso_signal | 14 | 3 | 2534 | 0.303 | 0.521 | 379.53 |
| 429 | sample_dat | lyso_signal | 15 | 3 | 208 | 1 | 1 | 531.5 |
| 430 | sample_dat | lyso_signal | 16 | 3 | 965 | 0.703 | 0.753 | 512.91 |
| 431 | sample_dat | lyso_signal | 17 | 3 | 502 | 0.985 | 0.999 | 311.22 |

| | | | | | | | | |
|---|---|---|---|---|---|---|---|---|
| 432 | sample_dat | lyso_signal | 18 | 3 | 185 | 1 | 1 | 469 |
| 433 | sample_dat | lyso_signal | 19 | 3 | 326 | 1 | 1 | 434.97 |
| 434 | sample_dat | lyso_signal | 20 | 3 | 473 | 0.9 | 0.924 | 539.32 |
| 435 | sample_dat | lyso_signal | 21 | 3 | 232 | 1 | 1 | 458 |
| 436 | sample_dat | lyso_signal | 22 | 3 | 2945 | 0.344 | 0.655 | 428.15 |
| 437 | sample_dat | lyso_signal | 23 | 3 | 297 | 1 | 1 | 315.88 |
| 438 | sample_dat | lyso_signal | 24 | 3 | 200 | 1 | 1 | 565 |
| 439 | sample_dat | lyso_signal | 25 | 3 | 1191 | 0.602 | 0.726 | 349.58 |
| 440 | sample_dat | lyso_signal | 26 | 3 | 255 | 1 | 1 | 470.51 |
| 441 | sample_dat | lyso_signal | 27 | 3 | 1738 | 0.618 | 0.749 | 585.38 |
| 442 | sample_dat | lyso_signal | 28 | 3 | 248 | 1 | 1 | 590.5 |
| 443 | sample_dat | lyso_signal | 29 | 3 | 281 | 1 | 1 | 569.74 |
| 444 | sample_dat | lyso_signal | 30 | 3 | 353 | 1 | 1 | 611.59 |
| 445 | sample_dat | lyso_signal | 31 | 3 | 320 | 1 | 1 | 624.5 |
| 446 | sample_dat | lyso_signal | 32 | 3 | 527 | 0.966 | 0.982 | 286.47 |
| 447 | sample_dat | lyso_signal | 33 | 3 | 359 | 1 | 1 | 603.72 |
| 448 | sample_dat | lyso_signal | 34 | 3 | 808 | 0.633 | 0.796 | 542.14 |
| 449 | sample_dat | lyso_signal | 35 | 3 | 29444 | 0.171 | 0.673 | 465.33 |
| 450 | sample_dat | lyso_signal | 36 | 3 | 216 | 1 | 1 | 569.5 |
| 451 | sample_dat | lyso_signal | 37 | 3 | 348 | 0.978 | 1 | 607.63 |
| 452 | sample_dat | lyso_signal | 38 | 3 | 185 | 1 | 1 | 583 |
| 453 | sample_dat | lyso_signal | 39 | 3 | 248 | 1 | 1 | 214.5 |
| 454 | sample_dat | lyso_signal | 40 | 3 | 435 | 0.73 | 0.832 | 273.87 |
| 455 | sample_dat | lyso_signal | 41 | 3 | 379 | 1 | 1 | 568.01 |
| 456 | sample_dat | lyso_signal | 42 | 3 | 1681 | 0.574 | 0.645 | 545.35 |
| 457 | sample_dat | lyso_signal | 43 | 3 | 323 | 1 | 1 | 275.59 |
| 458 | sample_dat | lyso_signal | 44 | 3 | 447 | 0.992 | 1 | 343.94 |
| 459 | sample_dat | lyso_signal | 45 | 3 | 623 | 1 | 1 | 295.24 |
| 460 | sample_dat | lyso_signal | 46 | 3 | 316 | 1 | 1 | 259.89 |
| 461 | sample_dat | lyso_signal | 47 | 3 | 424 | 0.997 | 1 | 356.21 |
| 462 | sample_dat | lyso_signal | 48 | 3 | 624 | 0.877 | 0.913 | 511.39 |
| 463 | sample_dat | lyso_signal | 49 | 3 | 604 | 0.678 | 0.786 | 406.27 |
| 464 | sample_dat | lyso_signal | 50 | 3 | 1051 | 0.777 | 0.812 | 403.88 |
| 465 | sample_dat | lyso_signal | 51 | 3 | 430 | 0.699 | 0.804 | 556.75 |
| 466 | sample_dat | lyso_signal | 52 | 3 | 261 | 1 | 1 | 533.62 |
| 467 | sample_dat | lyso_signal | 53 | 3 | 546 | 0.976 | 0.992 | 370 |
| 468 | sample_dat | lyso_signal | 54 | 3 | 315 | 1 | 1 | 312.38 |
| 469 | sample_dat | lyso_signal | 55 | 3 | 200 | 1 | 1 | 534 |
| 470 | sample_dat | lyso_signal | 56 | 3 | 204 | 1 | 1 | 379.62 |
| 471 | sample_dat | lyso_signal | 57 | 3 | 328 | 0.983 | 1 | 357.18 |
| 472 | sample_dat | lyso_signal | 58 | 3 | 249 | 1 | 1 | 241.61 |
| 473 | sample_dat | lyso_signal | 59 | 3 | 290 | 1 | 1 | 449.56 |
| 474 | sample_dat | lyso_signal | 60 | 3 | 2119 | 0.597 | 0.718 | 413.47 |
| 475 | sample_dat | lyso_signal | 61 | 3 | 286 | 1 | 1 | 334.42 |
| 476 | sample_dat | lyso_signal | 62 | 3 | 451 | 0.993 | 1 | 368.47 |
| 477 | sample_dat | lyso_signal | 63 | 3 | 339 | 1 | 1 | 385.73 |
| 478 | sample_dat | lyso_signal | 64 | 3 | 200 | 1 | 1 | 290.5 |
| 479 | sample_dat | lyso_signal | 65 | 3 | 430 | 0.961 | 0.975 | 405.49 |

| | | | | | | | | |
|---|---|---|---|---|---|---|---|---|
| 480 | sample_dat | lyso_signal | 66 | 3 | 216 | 1 | 1 | 397.5 |
| 481 | sample_dat | lyso_signal | 67 | 3 | 232 | 1 | 1 | 410 |
| 482 | sample_dat | lyso_signal | 68 | 3 | 261 | 1 | 1 | 460.89 |
| 483 | sample_dat | lyso_signal | 69 | 3 | 285 | 1 | 1 | 370.34 |
| 484 | sample_dat | lyso_signal | 70 | 3 | 1543 | 0.433 | 0.589 | 397.72 |
| 485 | sample_dat | lyso_signal | 71 | 3 | 200 | 1 | 1 | 433.5 |
| 486 | sample_dat | lyso_signal | 72 | 3 | 200 | 1 | 1 | 459 |
| 487 | sample_dat | lyso_signal | 73 | 3 | 250 | 1 | 1 | 422.64 |
| 488 | sample_dat | lyso_signal | 74 | 3 | 741 | 0.954 | 0.958 | 451.18 |
| 489 | sample_dat | lyso_bkg | 1 | 3 | 94 | 0.055 | 0.2 | 345.23 |
| 490 | sample_dat | lyso_bkg | 2 | 3 | 102 | 0.049 | 0.177 | 412.37 |
| 491 | sample_dat | lyso_bkg | 3 | 3 | 62 | 0.082 | 0.281 | 440 |
| 492 | sample_dat | lyso_bkg | 4 | 3 | 72 | 0.068 | 0.236 | 391.42 |
| 493 | sample_dat | lyso_bkg | 5 | 3 | 58 | 0.09 | 0.308 | 490.5 |
| 494 | sample_dat | lyso_bkg | 6 | 3 | 162 | 0.03 | 0.13 | 448.89 |
| 495 | sample_dat | lyso_bkg | 7 | 3 | 74 | 0.068 | 0.237 | 411.59 |
| 496 | sample_dat | lyso_bkg | 8 | 3 | 86 | 0.062 | 0.222 | 484 |
| 497 | sample_dat | lyso_bkg | 9 | 3 | 64 | 0.08 | 0.277 | 435.41 |
| 498 | sample_dat | lyso_bkg | 10 | 3 | 62 | 0.084 | 0.288 | 304.08 |
| 499 | sample_dat | lyso_bkg | 11 | 3 | 90 | 0.058 | 0.205 | 395.89 |
| 500 | sample_dat | lyso_bkg | 12 | 3 | 60 | 0.088 | 0.301 | 514.58 |
| 501 | sample_dat | lyso_bkg | 13 | 3 | 76 | 0.065 | 0.229 | 331.59 |
| 502 | sample_dat | lyso_bkg | 14 | 3 | 402 | 0.012 | 0.083 | 380.62 |
| 503 | sample_dat | lyso_bkg | 15 | 3 | 58 | 0.086 | 0.294 | 531.5 |
| 504 | sample_dat | lyso_bkg | 16 | 3 | 158 | 0.031 | 0.123 | 512.91 |
| 505 | sample_dat | lyso_bkg | 17 | 3 | 98 | 0.055 | 0.195 | 311.28 |
| 506 | sample_dat | lyso_bkg | 18 | 3 | 56 | 0.095 | 0.324 | 469 |
| 507 | sample_dat | lyso_bkg | 19 | 3 | 74 | 0.067 | 0.233 | 434.97 |
| 508 | sample_dat | lyso_bkg | 20 | 3 | 94 | 0.051 | 0.184 | 539.27 |
| 509 | sample_dat | lyso_bkg | 21 | 3 | 62 | 0.082 | 0.281 | 458 |
| 510 | sample_dat | lyso_bkg | 22 | 3 | 388 | 0.012 | 0.086 | 429.43 |
| 511 | sample_dat | lyso_bkg | 23 | 3 | 70 | 0.071 | 0.244 | 315.93 |
| 512 | sample_dat | lyso_bkg | 24 | 3 | 58 | 0.09 | 0.308 | 565 |
| 513 | sample_dat | lyso_bkg | 25 | 3 | 194 | 0.026 | 0.118 | 348.6 |
| 514 | sample_dat | lyso_bkg | 26 | 3 | 66 | 0.078 | 0.271 | 470.52 |
| 515 | sample_dat | lyso_bkg | 27 | 3 | 230 | 0.022 | 0.099 | 587.99 |
| 516 | sample_dat | lyso_bkg | 28 | 3 | 64 | 0.078 | 0.27 | 590.5 |
| 517 | sample_dat | lyso_bkg | 29 | 3 | 70 | 0.075 | 0.259 | 569.66 |
| 518 | sample_dat | lyso_bkg | 30 | 3 | 78 | 0.065 | 0.228 | 611.56 |
| 519 | sample_dat | lyso_bkg | 31 | 3 | 74 | 0.069 | 0.24 | 624.5 |
| 520 | sample_dat | lyso_bkg | 32 | 3 | 98 | 0.051 | 0.183 | 286.3 |
| 521 | sample_dat | lyso_bkg | 33 | 3 | 78 | 0.064 | 0.223 | 603.64 |
| 522 | sample_dat | lyso_bkg | 34 | 3 | 156 | 0.033 | 0.154 | 541.28 |
| 523 | sample_dat | lyso_bkg | 35 | 3 | 1574 | 0.003 | 0.036 | 473.91 |
| 524 | sample_dat | lyso_bkg | 36 | 3 | 60 | 0.086 | 0.294 | 569.5 |
| 525 | sample_dat | lyso_bkg | 37 | 3 | 80 | 0.065 | 0.231 | 607.6 |
| 526 | sample_dat | lyso_bkg | 38 | 3 | 56 | 0.095 | 0.324 | 583 |
| 527 | sample_dat | lyso_bkg | 39 | 3 | 64 | 0.078 | 0.27 | 214.5 |

| | | | | | | | |
|---|---|---|---|---|---|---|---|
| 528 | sample_dat lyso_bkg | 40 | 3 | 106 | 0.05 | 0.203 | 273.9 |
| 529 | sample_dat lyso_bkg | 41 | 3 | 80 | 0.062 | 0.218 | 568 |
| 530 | sample_dat lyso_bkg | 42 | 3 | 230 | 0.021 | 0.088 | 544.08 |
| 531 | sample_dat lyso_bkg | 43 | 3 | 76 | 0.069 | 0.246 | 275.54 |
| 532 | sample_dat lyso_bkg | 44 | 3 | 90 | 0.057 | 0.203 | 343.92 |
| 533 | sample_dat lyso_bkg | 45 | 3 | 106 | 0.048 | 0.173 | 295.14 |
| 534 | sample_dat lyso_bkg | 46 | 3 | 72 | 0.068 | 0.235 | 259.93 |
| 535 | sample_dat lyso_bkg | 47 | 3 | 86 | 0.058 | 0.207 | 356.34 |
| 536 | sample_dat lyso_bkg | 48 | 3 | 118 | 0.046 | 0.173 | 511.22 |
| 537 | sample_dat lyso_bkg | 49 | 3 | 124 | 0.038 | 0.161 | 405.82 |
| 538 | sample_dat lyso_bkg | 50 | 3 | 154 | 0.031 | 0.119 | 404.01 |
| 539 | sample_dat lyso_bkg | 51 | 3 | 104 | 0.047 | 0.194 | 556.46 |
| 540 | sample_dat lyso_bkg | 52 | 3 | 66 | 0.077 | 0.264 | 533.58 |
| 541 | sample_dat lyso_bkg | 53 | 3 | 102 | 0.051 | 0.185 | 369.81 |
| 542 | sample_dat lyso_bkg | 54 | 3 | 72 | 0.068 | 0.235 | 312.43 |
| 543 | sample_dat lyso_bkg | 55 | 3 | 58 | 0.09 | 0.308 | 534 |
| 544 | sample_dat lyso_bkg | 56 | 3 | 58 | 0.088 | 0.301 | 379.57 |
| 545 | sample_dat lyso_bkg | 57 | 3 | 76 | 0.067 | 0.235 | 357.14 |
| 546 | sample_dat lyso_bkg | 58 | 3 | 66 | 0.08 | 0.277 | 241.58 |
| 547 | sample_dat lyso_bkg | 59 | 3 | 70 | 0.072 | 0.251 | 449.53 |
| 548 | sample_dat lyso_bkg | 60 | 3 | 268 | 0.02 | 0.091 | 412.84 |
| 549 | sample_dat lyso_bkg | 61 | 3 | 70 | 0.073 | 0.255 | 334.44 |
| 550 | sample_dat lyso_bkg | 62 | 3 | 88 | 0.055 | 0.195 | 368.61 |
| 551 | sample_dat lyso_bkg | 63 | 3 | 76 | 0.067 | 0.231 | 385.64 |
| 552 | sample_dat lyso_bkg | 64 | 3 | 58 | 0.09 | 0.308 | 290.5 |
| 553 | sample_dat lyso_bkg | 65 | 3 | 88 | 0.056 | 0.2 | 405.38 |
| 554 | sample_dat lyso_bkg | 66 | 3 | 60 | 0.086 | 0.294 | 397.5 |
| 555 | sample_dat lyso_bkg | 67 | 3 | 62 | 0.082 | 0.281 | 410 |
| 556 | sample_dat lyso_bkg | 68 | 3 | 66 | 0.077 | 0.264 | 460.92 |
| 557 | sample_dat lyso_bkg | 69 | 3 | 70 | 0.073 | 0.255 | 370.4 |
| 558 | sample_dat lyso_bkg | 70 | 3 | 262 | 0.02 | 0.1 | 397.9 |
| 559 | sample_dat lyso_bkg | 71 | 3 | 58 | 0.09 | 0.308 | 433.5 |
| 560 | sample_dat lyso_bkg | 72 | 3 | 58 | 0.09 | 0.308 | 459 |
| 561 | sample_dat lyso_bkg | 73 | 3 | 66 | 0.08 | 0.277 | 422.59 |
| 562 | sample_dat lyso_bkg | 74 | 3 | 116 | 0.041 | 0.15 | 451.09 |
| 563 | sample_dat dark_bkg | 1 | 3 | 28416 | 0.57 | 0.553 | 70.54 |

| CentreY [px | Mean488 | Mean561 |
|---|---|---|
| 80 | 692.81 | 754.91 |
| 149.54 | 965.62 | 1076.12 |
| 122.92 | 691.06 | 823.93 |
| 131.13 | 953.81 | 819.53 |
| 158.78 | 626.24 | 856.4 |
| 190.57 | 855.09 | 857.16 |
| 180.39 | 828.9 | 791.81 |
| 249.26 | 889.92 | 875.93 |
| 223.64 | 772.34 | 759.35 |
| 240.57 | 797.02 | 782.81 |
| 244 | 852.37 | 808.73 |
| 249.35 | 845.04 | 790.01 |
| 253.5 | 843.12 | 709.9 |
| 262.34 | 870.72 | 764.23 |
| 268.89 | 787.2 | 752.32 |
| 273.86 | 798.52 | 938.56 |
| 272.38 | 774.57 | 726.19 |
| 275.14 | 973.05 | 847.41 |
| 281 | 805.03 | 723.45 |
| 366.43 | 1057.29 | 1152.28 |
| 307.58 | 926.39 | 838.79 |
| 377.88 | 1094.61 | 1075.03 |
| 316.26 | 850.88 | 799.51 |
| 327.5 | 908.25 | 764.23 |
| 375.55 | 1098.45 | 948.78 |
| 334.33 | 893.05 | 727.23 |
| 343 | 875.01 | 718.58 |
| 356.16 | 962.56 | 750.47 |
| 358.88 | 961.43 | 736.64 |
| 378.61 | 1019.72 | 833.83 |
| 383.13 | 853.04 | 705.36 |
| 386.42 | 871.18 | 751.81 |
| 403.79 | 1002.69 | 840.64 |
| 400 | 809.94 | 702.92 |
| 425.64 | 764.8 | 721.11 |
| 433.07 | 1041.11 | 879.63 |
| 439.6 | 873.89 | 839.77 |
| 503.82 | 779.27 | 1077.65 |
| 484.87 | 855.37 | 783.62 |
| 509.27 | 850.36 | 977.37 |
| 504.15 | 679.77 | 925.7 |
| 488.5 | 859.88 | 812.67 |
| 501.67 | 726.21 | 935.57 |
| 524 | 790.25 | 694 |
| 531.12 | 664.62 | 689.3 ` |
| 534.17 | 931.99 | 739.27 |
| 536 | 675.05 | 587.1 |

| ROI check | |
|---|---|
| OK | OK |
| OK | |
| OK | |

| pH |
|---|
| 6.54 |
| 4.42 |
| **shape** |
| 0.9 |

| | | |
|---|---|---|
| 1 | lyso_signal | 105 |
| 1 | lyso_bkg | 105 |
| 1 | dark_bkg | 1 |
| 2 | lyso_signal | 101 |
| 2 | lyso_bkg | 101 |
| 2 | dark_bkg | 1 |
| 3 | lyso_signal | 74 |
| 3 | lyso_bkg | 74 |
| 3 | dark_bkg | 1 |

| dark_bkg | | 488 | 561 |
|---|---|---|---|
| | | 122.06 | 120.19 |
| | | 113.45 | 112.2 |
| | | 106.92 | 109.9 |
| mean_cyto | | | |
| | | 792.3971 | 573.6976 |
| | | 480.3562 | 501.0401 |
| | | 157.2723 | 421.0132 |
| cyto-dark | | | |
| | | 670.3371 | 453.5076 |
| | 6.54 | 366.9062 | 388.8401 |
| | 4.42 | 50.3523 | 311.1132 |
| lin.reg. | | 2.711878 | 3.981095 |

| ***LOCAL BACKGROUND CALIBRATION*** | | |
|---|---|---|
| **Lysosome pH** | | **all** |
| | Average | 4.788425 |
| | Std.dev. | 0.661531 |
| | Median | 4.865212 |
| | n | 105 |
| HIGH Cal. | Average | 6.52827 |
| LOW Cal. | Average | 4.334097 |

| mean_lyso | | | |
|---|---|---|---|
| | | 878.6171 | 839.2792 |
| | | 763.4751 | 792.7803 |
| | | 195.9657 | 699.7374 |
| lyso-dark | | | |
| | | 756.5571 | 719.0892 |
| | 6.54 | 650.0251 | 680.5803 |
| | 4.42 | 89.04568 | 589.8374 |

| | | |
|---|---|---|
| 539.64 | 497.73 | 790.57 |
| 560.54 | 868.21 | 857.46 |
| 554.27 | 596.86 | 739.91 |
| 568.6 | 902.08 | 702.84 |
| 568.27 | 643.21 | 661.13 |
| 708.55 | 947.01 | 1871.82 |
| 600.43 | 826.85 | 843.16 |
| 585.38 | 454.63 | 651.28 |
| 595.97 | 604.64 | 654.07 |
| 646.2 | 944.16 | 873.18 |
| 658.16 | 782.28 | 877.98 |
| 692.66 | 632.91 | 748.08 |
| 689 | 471.81 | 710.34 |
| 697.77 | 690.87 | 704.7 |
| 720.39 | 770.98 | 775.08 |
| 780.46 | 818.22 | 888.5 |
| 788.64 | 566.5 | 917.3 |
| 803.35 | 860.68 | 1054.03 |
| 841.58 | 755.94 | 939.61 |
| 847.14 | 677.86 | 759.31 |
| 860 | 706.37 | 775.09 |
| 867.01 | 789.28 | 824.99 |
| 875.29 | 827.04 | 741.86 |
| 882.33 | 936.35 | 896.3 |
| 911.94 | 829.83 | 1181.91 |
| 901.39 | 934.08 | 861.15 |
| 903.11 | 814.85 | 843.99 |
| 907.22 | 1441.44 | 1016.62 |
| 917.44 | 1165.87 | 773.81 |
| 937.53 | 958.02 | 846.09 |
| 928.69 | 913.78 | 887.65 |
| 995.3 | 1044.24 | 1021.93 |
| 940.62 | 842.08 | 739.26 |
| 942.11 | 1121.19 | 852.61 |
| 946.62 | 1091.51 | 746.33 |
| 969.21 | 1105.18 | 987.29 |
| 972.1 | 1205.64 | 864.11 |
| 974.58 | 996.61 | 954.04 |
| 974.5 | 1091.84 | 740.66 |
| 987 | 775.07 | 635.74 |
| 998.5 | 1045.79 | 723.57 |
| 1001.37 | 971.69 | 902.55 |
| 1044.53 | 1233.97 | 1113.77 |
| 1021.91 | 1200.92 | 1184.1 |
| 1042.5 | 877.37 | 749.51 |
| 1067.6 | 1121.5 | 1144.85 |
| 1066.29 | 936.12 | 724.28 |
| 1069.5 | 904.36 | 747.2 |

| | | |
|---|---|---|
| lin.reg. | 2.636364 | 4.021997 |

| ***LYSOSOME SIGNAL CALIBRATION*** | | |
|---|---|---|
| **Lysosome pH** | | **all** |
| | Average | 4.806847 |
| | Std.dev. | 0.64311 |
| | Median | 4.881496 |
| | n | 105 |
| HIGH Cal. | Average | 6.498245 |
| LOW Cal. | Average | 4.36517 |

**<u>COPY THIS:</u>**

| local cal | | lyso cal |
|---|---|---|
| mean | median | mean |
| 4.788425 | 4.865212 | 4.806847 |

| | | |
|---|---|---|
| 1080.5 | 862.88 | 712.38 |
| 1094.22 | 940.52 | 815.17 |
| 1096.29 | 1062.45 | 823.22 |
| 1095.5 | 967.66 | 730.71 |
| 1119.77 | 977.8 | 877.95 |
| 1125.41 | 991.14 | 917.49 |
| 1156.5 | 948.39 | 725.02 |
| 1221.92 | 1043.96 | 989.52 |
| 1236.39 | 1021.79 | 756.46 |
| 1255.88 | 943.3 | 807.18 |
| 80 | 696.9 | 641.44 |
| 143.75 | 789.82 | 638.13 |
| 123 | 617.83 | 615.64 |
| 131.07 | 903.99 | 667.11 |
| 158.85 | 620.29 | 643.03 |
| 191.38 | 824.81 | 642.65 |
| 180.26 | 767.74 | 578.79 |
| 250.41 | 801.4 | 605.57 |
| 223.59 | 773.05 | 629.59 |
| 240.49 | 756.44 | 578.42 |
| 244 | 809.12 | 696.82 |
| 249.42 | 783.49 | 621.93 |
| 253.5 | 790.43 | 601.28 |
| 262.42 | 819.53 | 643.19 |
| 268.93 | 705.49 | 553.46 |
| 273.89 | 811.49 | 586.18 |
| 272.43 | 771.25 | 571.03 |
| 275.09 | 924.63 | 698.89 |
| 281 | 746.02 | 560.78 |
| 366.11 | 774 | 557.2 |
| 307.56 | 826.4 | 625.78 |
| 372.48 | 945.28 | 688.57 |
| 316.34 | 853.11 | 647.6 |
| 327.5 | 897.69 | 613.37 |
| 368.98 | 894.92 | 623.65 |
| 334.4 | 855.97 | 575.43 |
| 343 | 849.34 | 597.03 |
| 355.62 | 875.42 | 601.74 |
| 358.92 | 920.59 | 641.47 |
| 378.76 | 916.6 | 581.02 |
| 383.08 | 823.06 | 573.76 |
| 386.44 | 866.89 | 600.8 |
| 403.66 | 886.58 | 588.23 |
| 400 | 791.88 | 591.8 |
| 425.58 | 786.97 | 511.07 |
| 433.05 | 902.51 | 599.67 |
| 439.57 | 589.47 | 477.11 |
| 504.35 | 524.35 | 480.17 |

| | | |
|---|---|---|
| 484.92 | 859.15 | 606.99 |
| 509.33 | 586.01 | 507.6 |
| 503.77 | 590.7 | 515.37 |
| 488.5 | 741.81 | 670.79 |
| 501.63 | 749.09 | 592.58 |
| 524 | 715.94 | 535.16 |
| 531.07 | 660.66 | 461.14 |
| 534.11 | 853.42 | 573.76 |
| 536 | 688.91 | 476.34 |
| 539.74 | 406.84 | 384.6 |
| 560.58 | 635.28 | 491.44 |
| 554.36 | 443.82 | 450.69 |
| 568.62 | 884.7 | 580.2 |
| 568.16 | 544.27 | 444.3 |
| 706.43 | 549.14 | 492.99 |
| 599.56 | 699.22 | 511.4 |
| 585.43 | 459.62 | 445.57 |
| 595.89 | 473.27 | 392.92 |
| 646.08 | 805.28 | 603.43 |
| 659.01 | 717.04 | 519.55 |
| 692.66 | 457.52 | 382.24 |
| 689 | 452.47 | 485.59 |
| 697.85 | 689.02 | 560.89 |
| 720.43 | 767.44 | 600.11 |
| 777.96 | 650.45 | 505.1 |
| 788.91 | 579.46 | 451.14 |
| 803.42 | 861.81 | 574.76 |
| 841.55 | 745.95 | 524.03 |
| 847.09 | 552.31 | 569.59 |
| 860 | 734.43 | 507.29 |
| 866.87 | 664.94 | 453.9 |
| 874.82 | 813.91 | 535.47 |
| 882.36 | 705.93 | 474.61 |
| 911.54 | 539.13 | 437.03 |
| 901.47 | 953.79 | 627.73 |
| 903.07 | 543.65 | 428.08 |
| 906.98 | 1322.42 | 787.69 |
| 917.29 | 1167.06 | 664.77 |
| 937.24 | 925.35 | 580.1 |
| 928.78 | 856.98 | 623.26 |
| 998.19 | 769.12 | 533.87 |
| 940.58 | 790.79 | 593.33 |
| 942.07 | 1064.14 | 648.8 |
| 946.57 | 972.53 | 610.83 |
| 969.33 | 776.23 | 509.81 |
| 972.97 | 1030.12 | 650.66 |
| 974.27 | 948.65 | 643.13 |
| 974.5 | 1072.15 | 627.43 |

| | | |
|---|---|---|
| 987 | 749.11 | 516.16 |
| 998.5 | 916.38 | 579.77 |
| 1001.42 | 983.76 | 641.97 |
| 1039.71 | 948.8 | 582.61 |
| 1022.2 | 907.32 | 546.2 |
| 1042.5 | 861.98 | 615.33 |
| 1068.61 | 914.86 | 618.56 |
| 1066.34 | 929.33 | 605.34 |
| 1069.5 | 890.01 | 613.04 |
| 1080.5 | 858.53 | 592.43 |
| 1094.43 | 886.79 | 634.31 |
| 1096.44 | 1020.22 | 661.01 |
| 1095.5 | 931.58 | 620.97 |
| 1119.16 | 864.2 | 560.74 |
| 1124.51 | 911.82 | 622.02 |
| 1156.5 | 857.41 | 580.1 |
| 1220.62 | 903.42 | 607.93 |
| 1236.43 | 970.14 | 584.87 |
| 1255.93 | 929.37 | 657.43 |
| 452.49 | 122.06 | 120.19 |
| 106 | 519.64 | 669.83 |
| 147.86 | 703.18 | 837.29 |
| 185.62 | 821.86 | 820.72 |
| 206.03 | 640.44 | 884.29 |
| 240.44 | 998.59 | 1119.9 |
| 246.75 | 709.74 | 875.39 |
| 249.2 | 665.53 | 654.9 |
| 248.49 | 837.66 | 829.18 |
| 297.54 | 811.88 | 791.75 |
| 292.52 | 674.95 | 903.54 |
| 309.21 | 656.96 | 734.2 |
| 311.41 | 818.92 | 710.32 |
| 325.61 | 904.21 | 910.94 |
| 335 | 772.51 | 599.77 |
| 341.01 | 830 | 845.73 |
| 338 | 740.25 | 690.12 |
| 350.12 | 997.02 | 645.35 |
| 358.23 | 717.35 | 783.86 |
| 360.34 | 683.4 | 706.05 |
| 358 | 612.6 | 817.7 |
| 377.4 | 1072.01 | 963.02 |
| 378.4 | 748.78 | 762.31 |
| 382.5 | 641.38 | 601.06 |
| 402.07 | 949.23 | 939.52 |
| 400.39 | 1306.15 | 828.74 |
| 401.89 | 535.77 | 653.08 |
| 403.13 | 965.71 | 762.28 |
| 414.08 | 836.18 | 1030.21 |

| | | |
|---|---|---|
| 428.48 | 807.15 | 723.16 |
| 485.58 | 970.27 | 934.48 |
| 464.02 | 1040.88 | 997.88 |
| 460.78 | 1022.54 | 862.6 |
| 459 | 731.69 | 733.75 |
| 472.66 | 658.34 | 661.31 |
| 466 | 991.2 | 896.03 |
| 480.33 | 624.33 | 986.9 |
| 475 | 331.29 | 555.36 |
| 486 | 657.26 | 657.03 |
| 493.33 | 513.18 | 766.98 |
| 521.07 | 1271.74 | 992.98 |
| 510.91 | 1240.25 | 1110.28 |
| 499 | 672.88 | 719.83 |
| 554.63 | 704.29 | 643.51 |
| 555 | 612.28 | 656.56 |
| 578.63 | 832.79 | 840.41 |
| 574.61 | 684.21 | 733.84 |
| 591.99 | 529.8 | 784.4 |
| 579 | 777.5 | 686.67 |
| 587.91 | 604.68 | 619.2 |
| 586 | 288.05 | 465.75 |
| 590.57 | 934.06 | 855.03 |
| 595.88 | 592.75 | 601.1 |
| 608.47 | 374.07 | 845.34 |
| 610.51 | 424.74 | 702.53 |
| 736.68 | 935.27 | 1407.48 |
| 625.11 | 659.5 | 793.38 |
| 630.03 | 741.97 | 815.61 |
| 654 | 410.44 | 635.16 |
| 662.31 | 674.67 | 868.64 |
| 667.08 | 671.36 | 755.05 |
| 674.77 | 746.53 | 836.48 |
| 693.99 | 729.43 | 733.86 |
| 694 | 783.63 | 792.56 |
| 710.36 | 614.1 | 630.2 |
| 723 | 569.63 | 761.48 |
| 733.19 | 533.97 | 761.9 |
| 731.11 | 676.63 | 651.55 |
| 751.44 | 624.63 | 825.47 |
| 764.87 | 533.39 | 682.71 |
| 767.5 | 751.7 | 663.38 |
| 777.09 | 772.71 | 833.48 |
| 785.5 | 677.51 | 637.94 |
| 809.19 | 821.16 | 1047.56 |
| 804.78 | 676.84 | 754.27 |
| 851.31 | 781.69 | 880.18 |
| 858.28 | 635.45 | 757.27 |

| | | |
|---|---|---|
| 869.43 | 644.73 | 770.94 |
| 892.07 | 1158.98 | 1065.95 |
| 898.25 | 977.71 | 846.22 |
| 907.93 | 596.25 | 974.28 |
| 915.2 | 710.23 | 705.18 |
| 914.88 | 674.48 | 659.02 |
| 944.65 | 1020.85 | 762.74 |
| 954.24 | 453.62 | 685.59 |
| 974.75 | 746.46 | 828.56 |
| 981.12 | 387.48 | 586.38 |
| 984.64 | 930.47 | 707.78 |
| 987.25 | 828.12 | 772.43 |
| 1017.33 | 1629 | 1287.7 |
| 1070.3 | 1345.4 | 1064.07 |
| 1076.5 | 342.49 | 619.29 |
| 1082.92 | 758.59 | 762.04 |
| 1081.88 | 705.71 | 714.04 |
| 1111.66 | 649.11 | 743.88 |
| 1119.44 | 518.3 | 687.23 |
| 1150.66 | 901.9 | 704.15 |
| 1179.75 | 960.21 | 795.1 |
| 1185.72 | 1556.66 | 759.93 |
| 1228.23 | 994.21 | 993.89 |
| 1289.15 | 966.51 | 891.23 |
| 1292.26 | 567.19 | 753.62 |
| 106 | 475.95 | 481.61 |
| 147.32 | 508.65 | 562.83 |
| 185.57 | 590.58 | 645.73 |
| 205.83 | 542.03 | 624.86 |
| 241.42 | 506.1 | 515.55 |
| 246.81 | 680.3 | 737.09 |
| 249.29 | 495.89 | 477.22 |
| 248.5 | 618.12 | 612.94 |
| 297.09 | 446.69 | 431.3 |
| 292.57 | 581.46 | 578.26 |
| 309.19 | 469.84 | 455.44 |
| 311.41 | 516.14 | 461.03 |
| 325.84 | 753.04 | 729.56 |
| 335 | 415.47 | 403.72 |
| 341.01 | 574.74 | 603.31 |
| 338 | 566.4 | 538.21 |
| 350.07 | 496.84 | 434.68 |
| 358.14 | 406.47 | 439.71 |
| 360.37 | 546.01 | 523.3 |
| 358 | 572.57 | 666.65 |
| 377.27 | 516.91 | 509.18 |
| 378.45 | 533.16 | 545.76 |
| 382.5 | 409.89 | 384.87 |

| | | |
|---|---|---|
| 401.56 | 847.75 | 832.11 |
| 400.43 | 486.62 | 443.29 |
| 401.93 | 421.96 | 409.75 |
| 403.26 | 454.69 | 430.08 |
| 414.24 | 456.78 | 427.86 |
| 428.48 | 410.83 | 398.87 |
| 487.58 | 397.79 | 418.13 |
| 463.17 | 658.8 | 570.45 |
| 460.88 | 507.73 | 497.59 |
| 459 | 523.55 | 531.48 |
| 472.44 | 490.7 | 410.05 |
| 466 | 908.73 | 775.98 |
| 480.05 | 496.58 | 566.56 |
| 475 | 338.02 | 375.7 |
| 486 | 493.52 | 518.58 |
| 492.88 | 465.23 | 491.65 |
| 518.18 | 547.63 | 554.5 |
| 510.67 | 460.91 | 436.88 |
| 499 | 555.39 | 590.63 |
| 554.58 | 376.99 | 404.38 |
| 555 | 422.08 | 474.42 |
| 578.57 | 324.66 | 370.41 |
| 574.57 | 528.03 | 555.27 |
| 591.6 | 363.25 | 452.66 |
| 579 | 555.64 | 552.34 |
| 587.88 | 346.68 | 367.95 |
| 586 | 295.21 | 332.34 |
| 590.53 | 511.31 | 607.47 |
| 595.67 | 351.22 | 362.69 |
| 608.49 | 400.8 | 639.05 |
| 610.51 | 357.42 | 419.13 |
| 727.57 | 455.48 | 514.01 |
| 625.05 | 409.49 | 457.8 |
| 630.12 | 434.37 | 467.36 |
| 654 | 393.41 | 459.02 |
| 662.03 | 621.41 | 606.83 |
| 667.2 | 333.33 | 378.12 |
| 674.91 | 379.68 | 455.95 |
| 694.1 | 402.82 | 431.24 |
| 694 | 676.16 | 690.05 |
| 710.42 | 343.71 | 344.81 |
| 723 | 551.78 | 638.95 |
| 732.89 | 323.12 | 464.55 |
| 731.07 | 336.23 | 384.39 |
| 751.47 | 632.13 | 622.11 |
| 765.46 | 428.6 | 452.84 |
| 767.5 | 411.85 | 420.22 |
| 777.06 | 360.14 | 420.68 |

| | | |
|---|---|---|
| 785.5 | 359.17 | 414.64 |
| 809.17 | 514.78 | 559.22 |
| 805.04 | 441.11 | 461.34 |
| 851.42 | 451.46 | 469.01 |
| 858.78 | 407.82 | 425.84 |
| 869.24 | 404.17 | 480.23 |
| 891.86 | 597.89 | 578.09 |
| 898.35 | 415.25 | 469.75 |
| 908.05 | 395.47 | 477.17 |
| 915.14 | 352.11 | 412.59 |
| 914.92 | 424.11 | 476.31 |
| 944.61 | 472.74 | 466.85 |
| 954.19 | 462.38 | 473.04 |
| 975.19 | 602.25 | 621.88 |
| 981.08 | 360 | 405.47 |
| 984.59 | 516.43 | 469.34 |
| 987.16 | 467.09 | 533.08 |
| 1017.34 | 391.86 | 432.64 |
| 1072.17 | 365.71 | 460.89 |
| 1076.5 | 333.48 | 429.05 |
| 1082.89 | 542.01 | 530.86 |
| 1081.92 | 454.26 | 493.36 |
| 1111.51 | 560.83 | 575.34 |
| 1119.47 | 438.5 | 438.95 |
| 1150.79 | 448.08 | 458.09 |
| 1180.09 | 466.57 | 510.45 |
| 1185.82 | 790.11 | 571.59 |
| 1229.06 | 509.55 | 531.18 |
| 1289.64 | 535.86 | 576.21 |
| 1292.35 | 561.47 | 612.6 |
| 457.45 | 113.45 | 112.2 |
| 278.05 | 177.55 | 832.29 |
| 307.54 | 225.23 | 778.19 |
| 323.5 | 183.29 | 722.21 |
| 347.39 | 161.75 | 630.47 |
| 363 | 239.42 | 909.18 |
| 379.3 | 207.97 | 731.33 |
| 375.63 | 160.52 | 610.91 |
| 410.17 | 235.7 | 755.39 |
| 414.64 | 248.21 | 729.87 |
| 416.38 | 210.02 | 669.11 |
| 428.26 | 184.24 | 740.63 |
| 435.37 | 209.17 | 641.38 |
| 442.65 | 220.33 | 676.4 |
| 471 | 202 | 791 |
| 453 | 244.77 | 772.56 |
| 482.74 | 237.23 | 766.68 |
| 478.29 | 247.65 | 809.47 |

| | | |
|---|---|---|
| 474 | 173.98 | 481.45 |
| 479.45 | 198.9 | 597.42 |
| 485.55 | 197.39 | 791.47 |
| 497.5 | 197.24 | 561.62 |
| 527.18 | 223.6 | 829.28 |
| 512.61 | 164.25 | 674.03 |
| 513.5 | 222.5 | 743.15 |
| 543.37 | 242.06 | 807.26 |
| 523.03 | 164.42 | 770.66 |
| 553.72 | 206.82 | 864.2 |
| 584.5 | 148.21 | 520.88 |
| 597.99 | 186.75 | 575.1 |
| 599.89 | 206.2 | 625.08 |
| 622 | 163.68 | 582.07 |
| 639.92 | 196.81 | 716.93 |
| 638.99 | 205.4 | 645.63 |
| 648.91 | 212.71 | 877.2 |
| 762.73 | 208.37 | 1153.63 |
| 684.5 | 182.53 | 646.36 |
| 719.33 | 133.71 | 585.88 |
| 759 | 131.9 | 495.92 |
| 764.5 | 159.44 | 608.46 |
| 782.31 | 183.51 | 521.25 |
| 791.04 | 172.56 | 687.48 |
| 818.38 | 205.23 | 880.37 |
| 823.23 | 185.33 | 602.47 |
| 833.51 | 200.06 | 708.19 |
| 845.09 | 232.18 | 784.11 |
| 855.11 | 141.48 | 671.01 |
| 859.99 | 201.67 | 716.15 |
| 879.19 | 194.99 | 726.94 |
| 881.37 | 211.1 | 654.85 |
| 923.7 | 225.24 | 886.68 |
| 912.9 | 158.17 | 597.31 |
| 908.11 | 191.96 | 587.55 |
| 923.57 | 220.58 | 793.55 |
| 929.6 | 137.36 | 627.81 |
| 939.5 | 183.62 | 565.99 |
| 944.89 | 141.35 | 519.89 |
| 971.72 | 142.08 | 597.96 |
| 983.12 | 183.19 | 708.32 |
| 996.98 | 220.31 | 616.11 |
| 1035.23 | 274.87 | 1032.82 |
| 1011.14 | 209.65 | 633.67 |
| 1015.52 | 225 | 688.72 |
| 1036.49 | 154.19 | 645.65 |
| 1080 | 170.36 | 618.93 |
| 1094.4 | 230.23 | 668.07 |

| | | |
|---|---|---|
| 1130.5 | 141.96 | 574.68 |
| 1153.5 | 227.93 | 666.15 |
| 1158.62 | 163.66 | 622.38 |
| 1191.15 | 251.27 | 802.3 |
| 1229.81 | 234.48 | 812.84 |
| 1211 | 170.92 | 542.39 |
| 1218.5 | 147.98 | 544.63 |
| 1285.09 | 178.74 | 676.75 |
| 1299.61 | 266.33 | 1075.85 |
| 278.01 | 172.38 | 660.03 |
| 307.36 | 166.25 | 485.19 |
| 323.5 | 203.52 | 636.77 |
| 347.42 | 154.68 | 385.26 |
| 363 | 215.38 | 761.79 |
| 380.01 | 152.16 | 393.2 |
| 375.77 | 140.53 | 319.38 |
| 410.22 | 182.72 | 517.6 |
| 414.59 | 187.19 | 516.83 |
| 416.42 | 182.84 | 502.53 |
| 428.37 | 144.67 | 338.59 |
| 435.42 | 164.93 | 418.12 |
| 442.59 | 165.01 | 407.04 |
| 471.21 | 150.51 | 373 |
| 453 | 196.79 | 612.86 |
| 483.49 | 168.69 | 419.44 |
| 478.2 | 175.52 | 495.71 |
| 474 | 137.95 | 308.55 |
| 479.47 | 148.81 | 322.07 |
| 485.53 | 199 | 622.68 |
| 497.5 | 152.15 | 352.19 |
| 528.43 | 152.51 | 383.58 |
| 512.57 | 161.14 | 444.9 |
| 513.5 | 188.67 | 583.07 |
| 542.96 | 174.12 | 414.96 |
| 523.02 | 155.74 | 571.45 |
| 553.78 | 150.07 | 364.43 |
| 584.5 | 143.03 | 319.06 |
| 597.99 | 142.43 | 322.17 |
| 599.94 | 147.77 | 324.06 |
| 622 | 143.91 | 365.11 |
| 639.8 | 138.33 | 351.54 |
| 638.99 | 140.28 | 314.65 |
| 648.97 | 178.28 | 566.24 |
| 768.17 | 156.83 | 483.07 |
| 684.5 | 174.28 | 472.5 |
| 719.4 | 131.3 | 300.61 |
| 759 | 131.16 | 322.09 |
| 764.5 | 157.92 | 431.84 |

| | | |
|---|---|---|
| 782.32 | 138.67 | 318.55 |
| 791.02 | 140.5 | 349.88 |
| 818.57 | 149.57 | 420.01 |
| 823.33 | 134.45 | 312.45 |
| 833.53 | 150.62 | 360.36 |
| 845.25 | 141.25 | 333.92 |
| 855.07 | 142.01 | 418.49 |
| 859.98 | 145.23 | 354.94 |
| 879.36 | 152.64 | 453.59 |
| 881.08 | 152.84 | 411.33 |
| 923.37 | 158.83 | 426.9 |
| 912.54 | 154.04 | 420.87 |
| 908.08 | 139.91 | 340.74 |
| 923.75 | 149.16 | 452.03 |
| 929.57 | 139.44 | 347.19 |
| 939.5 | 146.95 | 368.57 |
| 944.93 | 131.59 | 305.26 |
| 971.86 | 137.91 | 364.21 |
| 983.08 | 166.83 | 520.89 |
| 996.97 | 153.11 | 344.37 |
| 1036.1 | 141.52 | 351.14 |
| 1011.09 | 153.56 | 392.44 |
| 1015.73 | 137.07 | 324.48 |
| 1036.49 | 158.39 | 399.12 |
| 1080 | 160.41 | 471.79 |
| 1094.24 | 142.58 | 323.17 |
| 1130.5 | 140.02 | 352.52 |
| 1153.5 | 185.23 | 498.11 |
| 1158.58 | 155.03 | 384.36 |
| 1191.1 | 207.71 | 657.83 |
| 1227.81 | 170.09 | 453.41 |
| 1211 | 146.79 | 367.83 |
| 1218.5 | 149.69 | 378.97 |
| 1285.06 | 169.12 | 491.21 |
| 1299.26 | 165.94 | 499.89 |
| 437.71 | 106.92 | 109.9 |

| | 488-local | 561-local | ratio | pH LOC BKG | | pH LYSO | |
|---|---|---|---|---|---|---|---|
| 1 | -4.09 | 113.47 | -0.03604 | 3.883346 | 3.883346 | 3.92697 | 3.92697 |
| 106 | 175.8 | 437.99 | 0.401379 | 5.069586 | | 5.080179 | |
| 211 | 73.23 | 208.29 | 0.351577 | 4.934529 | 4.934529 | 4.948883 | 4.948883 |
| 212 | 49.82 | 152.42 | 0.32686 | 4.867499 | 4.867499 | 4.883719 | 4.883719 |
| 313 | 5.95 | 213.37 | 0.027886 | 4.056718 | 4.056718 | 4.095515 | 4.095515 |
| 414 | 30.28 | 214.51 | 0.141159 | 4.3639 | | 4.394144 | |
| 415 | 61.16 | 213.02 | 0.287109 | 4.7597 | 4.7597 | 4.778922 | 4.778922 |
| 489 | 88.52 | 270.36 | 0.327415 | 4.869005 | | 4.885183 | |
| 563 | -0.71 | 129.76 | -0.00547 | 3.966256 | 3.966256 | 4.007572 | 4.007572 |
| | 40.58 | 204.39 | 0.198542 | 4.519516 | | 4.545426 | |
| | 43.25 | 111.91 | 0.386471 | 5.029158 | 5.029158 | 5.040876 | 5.040876 |
| | 61.55 | 168.08 | 0.366195 | 4.97417 | 4.97417 | 4.98742 | 4.98742 |
| | 52.69 | 108.62 | 0.485086 | 5.296588 | 5.296588 | 5.30086 | 5.30086 |
| | 51.19 | 121.04 | 0.422918 | 5.127997 | | 5.136963 | |
| | 81.71 | 198.86 | 0.410892 | 5.095384 | 5.095384 | 5.105259 | 5.105259 |
| | -12.97 | 352.38 | -0.03681 | 3.881279 | 3.881279 | 3.924961 | 3.924961 |
| | 3.32 | 155.16 | 0.021397 | 4.039121 | 4.039121 | 4.078408 | 4.078408 |
| | 48.42 | 148.52 | 0.326017 | 4.865212 | 4.865212 | 4.881496 | 4.881496 |
| | 59.01 | 162.67 | 0.362759 | 4.964853 | 4.964853 | 4.978362 | 4.978362 |
| | 283.29 | 595.08 | 0.476054 | 5.272094 | | 5.277048 | |
| | 99.99 | 213.01 | 0.469415 | 5.25409 | 5.25409 | 5.259545 | 5.259545 |
| ratio | 149.33 | 386.46 | 0.386405 | 5.028977 | | 5.040701 | |
| 1.478117 | -2.23 | 151.91 | -0.01468 | 3.941285 | 3.941285 | 3.983296 | 3.983296 |
| 0.943592 | 10.56 | 150.86 | 0.069999 | 4.170922 | 4.170922 | 4.206539 | 4.206539 |
| 0.161846 | 203.53 | 325.13 | 0.625996 | 5.678719 | | 5.67235 | |
| | 37.08 | 151.8 | 0.244269 | 4.643522 | 4.643522 | 4.665979 | 4.665979 |
| | 25.67 | 121.55 | 0.211189 | 4.553813 | 4.553813 | 4.578768 | 4.578768 |
| | 87.14 | 148.73 | 0.585894 | 5.569967 | | 5.566627 | |
| | 40.84 | 95.17 | 0.429127 | 5.144834 | 5.144834 | 5.153332 | 5.153332 |
| **round** | 103.12 | 252.81 | 0.407895 | 5.087257 | 5.087257 | 5.097358 | 5.097358 |
| 4.703 | 29.98 | 131.6 | 0.227812 | 4.598892 | 4.598892 | 4.622592 | 4.622592 |
| 0.70247 | 4.29 | 151.01 | 0.028409 | 4.058136 | 4.058136 | 4.096893 | 4.096893 |
| 4.665315 | 116.11 | 252.41 | 0.460006 | 5.228574 | 5.228574 | 5.23474 | 5.23474 |
| 76 | 18.06 | 111.12 | 0.162527 | 4.421848 | 4.421848 | 4.450478 | 4.450478 |
| | -22.17 | 210.04 | -0.10555 | 3.694852 | 3.694852 | 3.743726 | 3.743726 |
| | 138.6 | 279.96 | 0.495071 | 5.323666 | 5.323666 | 5.327184 | 5.327184 |
| | 284.42 | 362.66 | 0.784261 | 6.107914 | 6.107914 | 6.089594 | 6.089594 |
| | 254.92 | 597.48 | 0.426659 | 5.138141 | | 5.146825 | |
| | -3.78 | 176.63 | -0.0214 | 3.923059 | 3.923059 | 3.965577 | 3.965577 |
| | 264.35 | 469.77 | 0.562722 | 5.507129 | | 5.505538 | |
| | 89.07 | 410.33 | 0.217069 | 4.56976 | | 4.594271 | |
| | 118.07 | 141.88 | 0.832182 | 6.237871 | 6.237871 | 6.215932 | 6.215932 |
| | -22.88 | 342.99 | -0.06671 | 3.800192 | 3.800192 | 3.846132 | 3.846132 |
| ratio | 74.31 | 158.84 | 0.467829 | 5.249791 | 5.249791 | 5.255366 | 5.255366 |
| 1.052105 | 3.96 | 228.16 | 0.017356 | 4.028163 | 4.028163 | 4.067755 | 4.067755 |
| 0.955104 | 78.57 | 165.51 | 0.474715 | 5.268463 | 5.268463 | 5.273518 | 5.273518 |
| 0.150966 | -13.86 | 110.76 | -0.12514 | 3.641742 | 3.641742 | 3.692095 | 3.692095 |

**round**
4.723801
0.68291
4.687166
76

median
4.881496

| | | | | | | |
|---|---|---|---|---|---|---|
| 90.89 | 405.97 | 0.223884 | 4.588239 | 4.588239 | 4.612236 | 4.612236 |
| 232.93 | 366.02 | 0.636386 | 5.706896 | | 5.699743 | |
| 153.04 | 289.22 | 0.529147 | 5.416078 | 5.416078 | 5.417023 | 5.417023 |
| 17.38 | 122.64 | 0.141716 | 4.36541 | | 4.395611 | |
| 98.94 | 216.83 | 0.456302 | 5.21853 | 5.21853 | 5.224976 | 5.224976 |
| 397.87 | 1378.83 | 0.288556 | 4.763624 | | 4.782737 | |
| 127.63 | 331.76 | 0.384706 | 5.02437 | | 5.036222 | |
| -4.99 | 205.71 | -0.02426 | 3.915311 | 3.915311 | 3.958046 | 3.958046 |
| 131.37 | 261.15 | 0.503044 | 5.345289 | 5.345289 | 5.348205 | 5.348205 |
| 138.88 | 269.75 | 0.514847 | 5.377297 | 5.377297 | 5.379322 | 5.379322 |
| 65.24 | 358.43 | 0.182016 | 4.4747 | | 4.501858 | |
| 175.39 | 365.84 | 0.479417 | 5.281216 | 5.281216 | 5.285916 | 5.285916 |
| 19.34 | 224.75 | 0.086051 | 4.214455 | 4.214455 | 4.24886 | 4.24886 |
| 1.85 | 143.81 | 0.012864 | 4.015981 | 4.015981 | 4.055912 | 4.055912 |
| 3.54 | 174.97 | 0.020232 | 4.035961 | 4.035961 | 4.075336 | 4.075336 |
| 167.77 | 383.4 | 0.437585 | 5.167771 | | 5.17563 | |
| -12.96 | 466.16 | -0.0278 | 3.9057 | 3.9057 | 3.948702 | 3.948702 |
| -1.13 | 479.27 | -0.00236 | 3.974701 | 3.974701 | 4.015782 | 4.015782 |
| 9.99 | 415.58 | 0.024039 | 4.046285 | 4.046285 | 4.085372 | 4.085372 |
| 125.55 | 189.72 | 0.661765 | 5.77572 | 5.77572 | 5.76665 | 5.76665 |
| -28.06 | 267.8 | -0.10478 | 3.696945 | 3.696945 | 3.74576 | 3.74576 |
| 124.34 | 371.09 | 0.335067 | 4.889755 | 4.889755 | 4.905356 | 4.905356 |
| 13.13 | 206.39 | 0.063617 | 4.153617 | | 4.189716 | |
| 230.42 | 421.69 | 0.54642 | 5.46292 | 5.46292 | 5.46256 | 5.46256 |
| 290.7 | 744.88 | 0.390264 | 5.039444 | | 5.050876 | |
| -19.71 | 233.42 | -0.08444 | 3.752103 | 3.752103 | 3.799383 | 3.799383 |
| 271.2 | 415.91 | 0.652064 | 5.749413 | 5.749413 | 5.741076 | 5.741076 |
| 119.02 | 228.93 | 0.519897 | 5.390992 | 5.390992 | 5.392635 | 5.392635 |
| -1.19 | 109.04 | -0.01091 | 3.951499 | 3.951499 | 3.993226 | 3.993226 |
| 32.67 | 265.99 | 0.122824 | 4.314179 | | 4.345807 | |
| 56.8 | 264.39 | 0.214834 | 4.563699 | 4.563699 | 4.588378 | 4.588378 |
| 275.12 | 488.06 | 0.563701 | 5.509784 | | 5.508119 | |
| 51.29 | 145.93 | 0.35147 | 4.934238 | 4.934238 | 4.9486 | 4.9486 |
| 57.05 | 203.81 | 0.279918 | 4.740197 | 4.740197 | 4.759962 | 4.759962 |
| 118.98 | 135.5 | 0.878081 | 6.362344 | 6.362344 | 6.336939 | 6.336939 |
| 328.95 | 477.48 | 0.688929 | 5.849387 | 5.849387 | 5.838266 | 5.838266 |
| 175.52 | 213.45 | 0.8223 | 6.211073 | | 6.18988 | |
| 47.96 | 310.91 | 0.154257 | 4.39942 | 4.39942 | 4.428675 | 4.428675 |
| 19.69 | 113.23 | 0.173894 | 4.452673 | 4.452673 | 4.480445 | 4.480445 |
| 25.96 | 119.58 | 0.217093 | 4.569825 | 4.569825 | 4.594334 | 4.594334 |
| 129.41 | 143.8 | 0.89993 | 6.421596 | 6.421596 | 6.394542 | 6.394542 |
| -12.07 | 260.58 | -0.04632 | 3.855481 | 3.855481 | 3.899882 | 3.899882 |
| 285.17 | 531.16 | 0.536882 | 5.437052 | | 5.437413 | |
| 293.6 | 637.9 | 0.46026 | 5.229264 | 5.229264 | 5.235411 | 5.235411 |
| 15.39 | 134.18 | 0.114697 | 4.292138 | 4.292138 | 4.32438 | 4.32438 |
| 206.64 | 526.29 | 0.392635 | 5.045874 | | 5.057127 | |
| 6.79 | 118.94 | 0.057088 | 4.135909 | 4.135909 | 4.172501 | 4.172501 |
| 14.35 | 134.16 | 0.106962 | 4.271162 | 4.271162 | 4.303988 | 4.303988 |

| | | | | | | |
|---|---|---|---|---|---|---|
| 4.35 | 119.95 | 0.036265 | 4.079441 | 4.079441 | 4.117605 | 4.117605 |
| 53.73 | 180.86 | 0.297081 | 4.786741 | | 4.80521 | |
| 42.23 | 162.21 | 0.260342 | 4.687109 | 4.687109 | 4.708352 | 4.708352 |
| 36.08 | 109.74 | 0.328777 | 4.872698 | 4.872698 | 4.888774 | 4.888774 |
| 113.6 | 317.21 | 0.358122 | 4.952279 | | 4.966138 | |
| 79.32 | 295.47 | 0.268454 | 4.709108 | | 4.729739 | |
| 90.98 | 144.92 | 0.627795 | 5.683597 | 5.683597 | 5.677093 | 5.677093 |
| 140.54 | 381.59 | 0.368301 | 4.979882 | | 4.992973 | |
| 51.65 | 171.59 | 0.301008 | 4.797392 | 4.797392 | 4.815565 | 4.815565 |
| 13.93 | 149.75 | 0.093022 | 4.233358 | 4.233358 | 4.267236 | 4.267236 |
| NaN | NaN | NaN | | | | |
| NaN | NaN | NaN | | | | |
| NaN | NaN | NaN | | | | |
| NaN | NaN | NaN | | | | |
| NaN | NaN | NaN | | | | |
| NaN | NaN | NaN | | | | |
| NaN | NaN | NaN | | | | |
| NaN | NaN | NaN | | | | |
| NaN | NaN | NaN | | | | |
| NaN | NaN | NaN | | | | |
| NaN | NaN | NaN | | | | |
| NaN | NaN | NaN | | | | |
| NaN | NaN | NaN | | | | |
| NaN | NaN | NaN | | | | |
| NaN | NaN | NaN | | | | |
| NaN | NaN | NaN | | | | |
| NaN | NaN | NaN | | | | |
| NaN | NaN | NaN | | | | |
| NaN | NaN | NaN | | | | |
| NaN | NaN | NaN | | | | |
| NaN | NaN | NaN | | | | |
| NaN | NaN | NaN | | | | |
| NaN | NaN | NaN | | | | |
| NaN | NaN | NaN | | | | |
| NaN | NaN | NaN | | | | |
| NaN | NaN | NaN | | | | |
| NaN | NaN | NaN | | | | |
| NaN | NaN | NaN | | | | |
| NaN | NaN | NaN | | | | |
| NaN | NaN | NaN | | | | |
| NaN | NaN | NaN | | | | |
| NaN | NaN | NaN | | | | |
| NaN | NaN | NaN | | | | |
| NaN | NaN | NaN | | | | |
| NaN | NaN | NaN | | | | |
| NaN | NaN | NaN | | | | |
| NaN | NaN | NaN | | | | |
| NaN | NaN | NaN | | | | |

| | | |
|---|---|---|
| NaN | NaN | NaN |
| NaN | NaN | NaN |
| NaN | NaN | NaN |
| NaN | NaN | NaN |
| NaN | NaN | NaN |
| NaN | NaN | NaN |
| NaN | NaN | NaN |
| NaN | NaN | NaN |
| NaN | NaN | NaN |
| NaN | NaN | NaN |
| NaN | NaN | NaN |
| NaN | NaN | NaN |
| NaN | NaN | NaN |
| NaN | NaN | NaN |
| NaN | NaN | NaN |
| NaN | NaN | NaN |
| NaN | NaN | NaN |
| NaN | NaN | NaN |
| NaN | NaN | NaN |
| NaN | NaN | NaN |
| NaN | NaN | NaN |
| NaN | NaN | NaN |
| NaN | NaN | NaN |
| NaN | NaN | NaN |
| NaN | NaN | NaN |
| NaN | NaN | NaN |
| NaN | NaN | NaN |
| NaN | NaN | NaN |
| NaN | NaN | NaN |
| NaN | NaN | NaN |
| NaN | NaN | NaN |
| NaN | NaN | NaN |
| NaN | NaN | NaN |
| NaN | NaN | NaN |
| NaN | NaN | NaN |
| NaN | NaN | NaN |
| NaN | NaN | NaN |
| NaN | NaN | NaN |
| NaN | NaN | NaN |
| NaN | NaN | NaN |
| NaN | NaN | NaN |
| NaN | NaN | NaN |
| NaN | NaN | NaN |
| NaN | NaN | NaN |
| NaN | NaN | NaN |
| NaN | NaN | NaN |
| NaN | NaN | NaN |
| NaN | NaN | NaN |

| | | |
|---|---|---|
| NaN | NaN | NaN |
| NaN | NaN | NaN |
| NaN | NaN | NaN |
| NaN | NaN | NaN |
| NaN | NaN | NaN |
| NaN | NaN | NaN |
| NaN | NaN | NaN |
| NaN | NaN | NaN |
| NaN | NaN | NaN |
| NaN | NaN | NaN |
| NaN | NaN | NaN |
| NaN | NaN | NaN |
| NaN | NaN | NaN |
| NaN | NaN | NaN |
| NaN | NaN | NaN |
| NaN | NaN | NaN |
| NaN | NaN | NaN |
| NaN | NaN | NaN |
| NaN | NaN | NaN |
| NaN | NaN | NaN |
| NaN | NaN | NaN |
| NaN | NaN | NaN |
| NaN | NaN | NaN |
| NaN | NaN | NaN |
| NaN | NaN | NaN |
| NaN | NaN | NaN |
| NaN | NaN | NaN |
| NaN | NaN | NaN |
| NaN | NaN | NaN |
| NaN | NaN | NaN |
| NaN | NaN | NaN |
| NaN | NaN | NaN |
| NaN | NaN | NaN |
| NaN | NaN | NaN |
| NaN | NaN | NaN |
| NaN | NaN | NaN |
| NaN | NaN | NaN |
| NaN | NaN | NaN |
| NaN | NaN | NaN |
| NaN | NaN | NaN |
| NaN | NaN | NaN |
| NaN | NaN | NaN |
| NaN | NaN | NaN |
| NaN | NaN | NaN |
| NaN | NaN | NaN |
| NaN | NaN | NaN |
| NaN | NaN | NaN |
| NaN | NaN | NaN |

| | | |
|---|---|---|
| NaN | NaN | NaN |
| NaN | NaN | NaN |
| NaN | NaN | NaN |
| NaN | NaN | NaN |
| NaN | NaN | NaN |
| NaN | NaN | NaN |
| NaN | NaN | NaN |
| NaN | NaN | NaN |
| NaN | NaN | NaN |
| NaN | NaN | NaN |
| NaN | NaN | NaN |
| NaN | NaN | NaN |
| NaN | NaN | NaN |
| NaN | NaN | NaN |
| NaN | NaN | NaN |
| NaN | NaN | NaN |
| NaN | NaN | NaN |
| NaN | NaN | NaN |
| NaN | NaN | NaN |
| NaN | NaN | NaN |
| NaN | NaN | NaN |
| NaN | NaN | NaN |
| NaN | NaN | NaN |
| NaN | NaN | NaN |
| NaN | NaN | NaN |
| NaN | NaN | NaN |
| NaN | NaN | NaN |
| NaN | NaN | NaN |
| NaN | NaN | NaN |
| NaN | NaN | NaN |
| NaN | NaN | NaN |
| NaN | NaN | NaN |
| NaN | NaN | NaN |
| NaN | NaN | NaN |
| NaN | NaN | NaN |
| NaN | NaN | NaN |
| NaN | NaN | NaN |
| NaN | NaN | NaN |
| NaN | NaN | NaN |
| NaN | NaN | NaN |
| NaN | NaN | NaN |
| NaN | NaN | NaN |
| NaN | NaN | NaN |
| NaN | NaN | NaN |
| NaN | NaN | NaN |
| NaN | NaN | NaN |
| NaN | NaN | NaN |
| NaN | NaN | NaN |

| | | |
|---|---|---|
| NaN | NaN | NaN |
| NaN | NaN | NaN |
| NaN | NaN | NaN |
| NaN | NaN | NaN |
| NaN | NaN | NaN |
| NaN | NaN | NaN |
| NaN | NaN | NaN |
| NaN | NaN | NaN |
| NaN | NaN | NaN |
| NaN | NaN | NaN |
| NaN | NaN | NaN |
| NaN | NaN | NaN |
| NaN | NaN | NaN |
| NaN | NaN | NaN |
| NaN | NaN | NaN |
| NaN | NaN | NaN |
| NaN | NaN | NaN |
| NaN | NaN | NaN |
| NaN | NaN | NaN |
| NaN | NaN | NaN |
| NaN | NaN | NaN |
| NaN | NaN | NaN |
| NaN | NaN | NaN |
| NaN | NaN | NaN |
| NaN | NaN | NaN |
| NaN | NaN | NaN |
| NaN | NaN | NaN |
| NaN | NaN | NaN |
| NaN | NaN | NaN |
| NaN | NaN | NaN |
| NaN | NaN | NaN |
| NaN | NaN | NaN |
| NaN | NaN | NaN |
| NaN | NaN | NaN |
| NaN | NaN | NaN |
| NaN | NaN | NaN |
| NaN | NaN | NaN |
| NaN | NaN | NaN |
| NaN | NaN | NaN |
| NaN | NaN | NaN |
| NaN | NaN | NaN |
| NaN | NaN | NaN |
| NaN | NaN | NaN |
| NaN | NaN | NaN |
| NaN | NaN | NaN |
| NaN | NaN | NaN |
| NaN | NaN | NaN |
| NaN | NaN | NaN |

| | | |
|---|---|---|
| NaN | NaN | NaN |
| NaN | NaN | NaN |
| NaN | NaN | NaN |
| NaN | NaN | NaN |
| NaN | NaN | NaN |
| NaN | NaN | NaN |
| NaN | NaN | NaN |
| NaN | NaN | NaN |
| NaN | NaN | NaN |
| NaN | NaN | NaN |
| NaN | NaN | NaN |
| NaN | NaN | NaN |
| NaN | NaN | NaN |
| NaN | NaN | NaN |
| NaN | NaN | NaN |
| NaN | NaN | NaN |
| NaN | NaN | NaN |
| NaN | NaN | NaN |
| NaN | NaN | NaN |
| NaN | NaN | NaN |
| NaN | NaN | NaN |
| NaN | NaN | NaN |
| NaN | NaN | NaN |
| NaN | NaN | NaN |
| NaN | NaN | NaN |
| NaN | NaN | NaN |
| NaN | NaN | NaN |
| NaN | NaN | NaN |
| NaN | NaN | NaN |
| NaN | NaN | NaN |
| NaN | NaN | NaN |
| NaN | NaN | NaN |
| NaN | NaN | NaN |
| NaN | NaN | NaN |
| NaN | NaN | NaN |
| NaN | NaN | NaN |
| NaN | NaN | NaN |
| NaN | NaN | NaN |
| NaN | NaN | NaN |
| NaN | NaN | NaN |
| NaN | NaN | NaN |
| NaN | NaN | NaN |
| NaN | NaN | NaN |
| NaN | NaN | NaN |
| NaN | NaN | NaN |
| NaN | NaN | NaN |
| NaN | NaN | NaN |
| NaN | NaN | NaN |

| | | |
|---|---|---|
| NaN | NaN | NaN |
| NaN | NaN | NaN |
| NaN | NaN | NaN |
| NaN | NaN | NaN |
| NaN | NaN | NaN |
| NaN | NaN | NaN |
| NaN | NaN | NaN |
| NaN | NaN | NaN |
| NaN | NaN | NaN |
| NaN | NaN | NaN |
| NaN | NaN | NaN |
| NaN | NaN | NaN |
| NaN | NaN | NaN |
| NaN | NaN | NaN |
| NaN | NaN | NaN |
| NaN | NaN | NaN |
| NaN | NaN | NaN |
| NaN | NaN | NaN |
| NaN | NaN | NaN |
| NaN | NaN | NaN |
| NaN | NaN | NaN |
| NaN | NaN | NaN |
| NaN | NaN | NaN |
| NaN | NaN | NaN |
| NaN | NaN | NaN |
| NaN | NaN | NaN |
| NaN | NaN | NaN |
| NaN | NaN | NaN |
| NaN | NaN | NaN |
| NaN | NaN | NaN |
| NaN | NaN | NaN |
| NaN | NaN | NaN |
| NaN | NaN | NaN |
| NaN | NaN | NaN |
| NaN | NaN | NaN |
| NaN | NaN | NaN |
| NaN | NaN | NaN |
| NaN | NaN | NaN |
| NaN | NaN | NaN |
| NaN | NaN | NaN |
| NaN | NaN | NaN |
| NaN | NaN | NaN |
| NaN | NaN | NaN |
| NaN | NaN | NaN |
| NaN | NaN | NaN |
| NaN | NaN | NaN |
| NaN | NaN | NaN |
| NaN | NaN | NaN |

| | | |
|---|---|---|
| NaN | NaN | NaN |
| NaN | NaN | NaN |
| NaN | NaN | NaN |
| NaN | NaN | NaN |
| NaN | NaN | NaN |
| NaN | NaN | NaN |
| NaN | NaN | NaN |
| NaN | NaN | NaN |
| NaN | NaN | NaN |
| NaN | NaN | NaN |
| NaN | NaN | NaN |
| NaN | NaN | NaN |
| NaN | NaN | NaN |
| NaN | NaN | NaN |
| NaN | NaN | NaN |
| NaN | NaN | NaN |
| NaN | NaN | NaN |
| NaN | NaN | NaN |
| NaN | NaN | NaN |
| NaN | NaN | NaN |
| NaN | NaN | NaN |
| NaN | NaN | NaN |
| NaN | NaN | NaN |
| NaN | NaN | NaN |
| NaN | NaN | NaN |
| NaN | NaN | NaN |
| NaN | NaN | NaN |
| NaN | NaN | NaN |
| NaN | NaN | NaN |
| NaN | NaN | NaN |
| NaN | NaN | NaN |
| NaN | NaN | NaN |
| NaN | NaN | NaN |
| NaN | NaN | NaN |
| NaN | NaN | NaN |
| NaN | NaN | NaN |
| NaN | NaN | NaN |
| NaN | NaN | NaN |
| NaN | NaN | NaN |
| NaN | NaN | NaN |
| NaN | NaN | NaN |
| NaN | NaN | NaN |
| NaN | NaN | NaN |
| NaN | NaN | NaN |
| NaN | NaN | NaN |
| NaN | NaN | NaN |
| NaN | NaN | NaN |
| NaN | NaN | NaN |

| | | |
|---|---|---|
| NaN | NaN | NaN |
| NaN | NaN | NaN |
| NaN | NaN | NaN |
| NaN | NaN | NaN |
| NaN | NaN | NaN |
| NaN | NaN | NaN |
| NaN | NaN | NaN |
| NaN | NaN | NaN |
| NaN | NaN | NaN |
| NaN | NaN | NaN |
| NaN | NaN | NaN |
| NaN | NaN | NaN |
| NaN | NaN | NaN |
| NaN | NaN | NaN |
| NaN | NaN | NaN |
| NaN | NaN | NaN |
| NaN | NaN | NaN |
| NaN | NaN | NaN |
| NaN | NaN | NaN |
| NaN | NaN | NaN |
| NaN | NaN | NaN |
| NaN | NaN | NaN |
| NaN | NaN | NaN |
| NaN | NaN | NaN |
| NaN | NaN | NaN |
| NaN | NaN | NaN |
| NaN | NaN | NaN |
| NaN | NaN | NaN |
| NaN | NaN | NaN |
| NaN | NaN | NaN |
| NaN | NaN | NaN |
| NaN | NaN | NaN |
| NaN | NaN | NaN |
| NaN | NaN | NaN |
| NaN | NaN | NaN |
| NaN | NaN | NaN |
| NaN | NaN | NaN |
| NaN | NaN | NaN |
| NaN | NaN | NaN |
| NaN | NaN | NaN |
| NaN | NaN | NaN |
| NaN | NaN | NaN |
| NaN | NaN | NaN |
| NaN | NaN | NaN |
| NaN | NaN | NaN |
| NaN | NaN | NaN |
| NaN | NaN | NaN |
| NaN | NaN | NaN |

| | | |
|---|---|---|
| NaN | NaN | NaN |
| NaN | NaN | NaN |
| NaN | NaN | NaN |
| NaN | NaN | NaN |
| NaN | NaN | NaN |
| NaN | NaN | NaN |
| NaN | NaN | NaN |
| NaN | NaN | NaN |
| NaN | NaN | NaN |
| NaN | NaN | NaN |
| NaN | NaN | NaN |
| NaN | NaN | NaN |
| NaN | NaN | NaN |
| NaN | NaN | NaN |
| NaN | NaN | NaN |
| NaN | NaN | NaN |
| NaN | NaN | NaN |
| NaN | NaN | NaN |
| NaN | NaN | NaN |
| NaN | NaN | NaN |
| NaN | NaN | NaN |
| NaN | NaN | NaN |
| NaN | NaN | NaN |
| NaN | NaN | NaN |
| NaN | NaN | NaN |
| NaN | NaN | NaN |
| NaN | NaN | NaN |
| NaN | NaN | NaN |
| NaN | NaN | NaN |
| NaN | NaN | NaN |
| NaN | NaN | NaN |
| NaN | NaN | NaN |
| NaN | NaN | NaN |
| NaN | NaN | NaN |
| NaN | NaN | NaN |
| NaN | NaN | NaN |
| NaN | NaN | NaN |
| NaN | NaN | NaN |
| NaN | NaN | NaN |
| NaN | NaN | NaN |
| NaN | NaN | NaN |
| NaN | NaN | NaN |
| NaN | NaN | NaN |
| NaN | NaN | NaN |
| NaN | NaN | NaN |
| NaN | NaN | NaN |
| NaN | NaN | NaN |
| NaN | NaN | NaN |

| | | |
|---|---|---|
| NaN | NaN | NaN |
| NaN | NaN | NaN |
| NaN | NaN | NaN |
| NaN | NaN | NaN |
| NaN | NaN | NaN |
| NaN | NaN | NaN |
| NaN | NaN | NaN |
| NaN | NaN | NaN |
| NaN | NaN | NaN |
| NaN | NaN | NaN |
| NaN | NaN | NaN |
| NaN | NaN | NaN |
| NaN | NaN | NaN |
| NaN | NaN | NaN |
| NaN | NaN | NaN |
| NaN | NaN | NaN |
| NaN | NaN | NaN |
| NaN | NaN | NaN |
| NaN | NaN | NaN |
| NaN | NaN | NaN |
| NaN | NaN | NaN |
| NaN | NaN | NaN |
| NaN | NaN | NaN |
| NaN | NaN | NaN |
| NaN | NaN | NaN |
| NaN | NaN | NaN |
| NaN | NaN | NaN |
| NaN | NaN | NaN |
| NaN | NaN | NaN |
| NaN | NaN | NaN |
| NaN | NaN | NaN |
| NaN | NaN | NaN |
| NaN | NaN | NaN |
| NaN | NaN | NaN |
| NaN | NaN | NaN |
| NaN | NaN | NaN |
| NaN | NaN | NaN |
| NaN | NaN | NaN |
| NaN | NaN | NaN |
| NaN | NaN | NaN |
| NaN | NaN | NaN |
| NaN | NaN | NaN |
| NaN | NaN | NaN |
| NaN | NaN | NaN |
| NaN | NaN | NaN |
| NaN | NaN | NaN |
| NaN | NaN | NaN |
| NaN | NaN | NaN |

| | | |
|---|---|---|
| NaN | NaN | NaN |
| NaN | NaN | NaN |
| NaN | NaN | NaN |
| NaN | NaN | NaN |
| NaN | NaN | NaN |
| NaN | NaN | NaN |
| NaN | NaN | NaN |
| NaN | NaN | NaN |
| NaN | NaN | NaN |
| NaN | NaN | NaN |
| NaN | NaN | NaN |
| NaN | NaN | NaN |
| NaN | NaN | NaN |
| NaN | NaN | NaN |
| NaN | NaN | NaN |
| NaN | NaN | NaN |
| NaN | NaN | NaN |
| NaN | NaN | NaN |
| NaN | NaN | NaN |
| NaN | NaN | NaN |
| NaN | NaN | NaN |
| NaN | NaN | NaN |
| NaN | NaN | NaN |
| NaN | NaN | NaN |
| NaN | NaN | NaN |
| NaN | NaN | NaN |
| NaN | NaN | NaN |
| NaN | NaN | NaN |
| NaN | NaN | NaN |
| NaN | NaN | NaN |
| NaN | NaN | NaN |
| NaN | NaN | NaN |
| NaN | NaN | NaN |
| NaN | NaN | NaN |
| NaN | NaN | NaN |
| NaN | NaN | NaN |
| NaN | NaN | NaN |
| NaN | NaN | NaN |
| NaN | NaN | NaN |
| NaN | NaN | NaN |
| NaN | NaN | NaN |
| NaN | NaN | NaN |
| NaN | NaN | NaN |
| NaN | NaN | NaN |
| NaN | NaN | NaN |
| NaN | NaN | NaN |
| NaN | NaN | NaN |
| NaN | NaN | NaN |

| | | |
|---|---|---|
| NaN | NaN | NaN |
| NaN | NaN | NaN |
| NaN | NaN | NaN |
| NaN | NaN | NaN |
| NaN | NaN | NaN |
| NaN | NaN | NaN |
| NaN | NaN | NaN |
| NaN | NaN | NaN |
| NaN | NaN | NaN |
| NaN | NaN | NaN |
| NaN | NaN | NaN |
| NaN | NaN | NaN |
| NaN | NaN | NaN |
| NaN | NaN | NaN |
| NaN | NaN | NaN |
| NaN | NaN | NaN |
| NaN | NaN | NaN |
| NaN | NaN | NaN |
| NaN | NaN | NaN |
| NaN | NaN | NaN |
| NaN | NaN | NaN |
| NaN | NaN | NaN |
| NaN | NaN | NaN |
| NaN | NaN | NaN |
| NaN | NaN | NaN |
| NaN | NaN | NaN |
| NaN | NaN | NaN |
| NaN | NaN | NaN |
| NaN | NaN | NaN |
| NaN | NaN | NaN |
| NaN | NaN | NaN |
| NaN | NaN | NaN |
| NaN | NaN | NaN |
| NaN | NaN | NaN |
| NaN | NaN | NaN |
| NaN | NaN | NaN |
| NaN | NaN | NaN |
| NaN | NaN | NaN |
| NaN | NaN | NaN |
| NaN | NaN | NaN |
| NaN | NaN | NaN |
| NaN | NaN | NaN |
| NaN | NaN | NaN |
| NaN | NaN | NaN |
| NaN | NaN | NaN |
| NaN | NaN | NaN |
| NaN | NaN | NaN |
| NaN | NaN | NaN |

| | | |
|---|---|---|
| NaN | NaN | NaN |
| NaN | NaN | NaN |
| NaN | NaN | NaN |
| NaN | NaN | NaN |
| NaN | NaN | NaN |
| NaN | NaN | NaN |
| NaN | NaN | NaN |
| NaN | NaN | NaN |
| NaN | NaN | NaN |
| NaN | NaN | NaN |
| NaN | NaN | NaN |
| NaN | NaN | NaN |
| NaN | NaN | NaN |
| NaN | NaN | NaN |
| NaN | NaN | NaN |
| NaN | NaN | NaN |
| NaN | NaN | NaN |
| NaN | NaN | NaN |
| NaN | NaN | NaN |
| NaN | NaN | NaN |
| NaN | NaN | NaN |
| NaN | NaN | NaN |
| NaN | NaN | NaN |
| NaN | NaN | NaN |
| NaN | NaN | NaN |
| NaN | NaN | NaN |
| NaN | NaN | NaN |
| NaN | NaN | NaN |
| NaN | NaN | NaN |
| NaN | NaN | NaN |
| NaN | NaN | NaN |
| NaN | NaN | NaN |
| NaN | NaN | NaN |
| NaN | NaN | NaN |
| NaN | NaN | NaN |
| NaN | NaN | NaN |
| NaN | NaN | NaN |
| NaN | NaN | NaN |
| NaN | NaN | NaN |
| NaN | NaN | NaN |
| NaN | NaN | NaN |
| NaN | NaN | NaN |
| NaN | NaN | NaN |
| NaN | NaN | NaN |
| NaN | NaN | NaN |
| NaN | NaN | NaN |
| NaN | NaN | NaN |
| NaN | NaN | NaN |

| | | |
|---|---|---|
| NaN | NaN | NaN |
| NaN | NaN | NaN |
| NaN | NaN | NaN |
| NaN | NaN | NaN |
| NaN | NaN | NaN |
| NaN | NaN | NaN |
| NaN | NaN | NaN |
| NaN | NaN | NaN |
| NaN | NaN | NaN |
| NaN | NaN | NaN |
| NaN | NaN | NaN |
| NaN | NaN | NaN |
| NaN | NaN | NaN |
| NaN | NaN | NaN |
| NaN | NaN | NaN |
| NaN | NaN | NaN |
| NaN | NaN | NaN |
| NaN | NaN | NaN |
| NaN | NaN | NaN |
| NaN | NaN | NaN |
| NaN | NaN | NaN |
| NaN | NaN | NaN |
| NaN | NaN | NaN |
| NaN | NaN | NaN |
| NaN | NaN | NaN |
| NaN | NaN | NaN |
| NaN | NaN | NaN |
| NaN | NaN | NaN |
| NaN | NaN | NaN |
| NaN | NaN | NaN |
| NaN | NaN | NaN |
| NaN | NaN | NaN |
| NaN | NaN | NaN |
| NaN | NaN | NaN |
| NaN | NaN | NaN |
| NaN | NaN | NaN |
| NaN | NaN | NaN |
| NaN | NaN | NaN |
| NaN | NaN | NaN |
| NaN | NaN | NaN |
| NaN | NaN | NaN |
| NaN | NaN | NaN |
| NaN | NaN | NaN |
| NaN | NaN | NaN |
| NaN | NaN | NaN |
| NaN | NaN | NaN |
| NaN | NaN | NaN |
| NaN | NaN | NaN |

| | | |
|---|---|---|
| NaN | NaN | NaN |
| NaN | NaN | NaN |
| NaN | NaN | NaN |
| NaN | NaN | NaN |
| NaN | NaN | NaN |
| NaN | NaN | NaN |
| NaN | NaN | NaN |
| NaN | NaN | NaN |
| NaN | NaN | NaN |
| NaN | NaN | NaN |
| NaN | NaN | NaN |
| NaN | NaN | NaN |
| NaN | NaN | NaN |
| NaN | NaN | NaN |
| NaN | NaN | NaN |
| NaN | NaN | NaN |
| NaN | NaN | NaN |
| NaN | NaN | NaN |
| NaN | NaN | NaN |
| NaN | NaN | NaN |
| NaN | NaN | NaN |
| NaN | NaN | NaN |
| NaN | NaN | NaN |
| NaN | NaN | NaN |
| NaN | NaN | NaN |
| NaN | NaN | NaN |
| NaN | NaN | NaN |
| NaN | NaN | NaN |
| NaN | NaN | NaN |
| NaN | NaN | NaN |
| NaN | NaN | NaN |
| NaN | NaN | NaN |
| NaN | NaN | NaN |
| NaN | NaN | NaN |
| NaN | NaN | NaN |
| NaN | NaN | NaN |
| NaN | NaN | NaN |
| NaN | NaN | NaN |
| NaN | NaN | NaN |
| NaN | NaN | NaN |
| NaN | NaN | NaN |
| NaN | NaN | NaN |
| NaN | NaN | NaN |
| NaN | NaN | NaN |
| NaN | NaN | NaN |
| NaN | NaN | NaN |
| NaN | NaN | NaN |
| NaN | NaN | NaN |

| | | |
|---|---|---|
| NaN | NaN | NaN |
| NaN | NaN | NaN |
| NaN | NaN | NaN |
| NaN | NaN | NaN |
| NaN | NaN | NaN |
| NaN | NaN | NaN |
| NaN | NaN | NaN |
| NaN | NaN | NaN |
| NaN | NaN | NaN |
| NaN | NaN | NaN |
| NaN | NaN | NaN |
| NaN | NaN | NaN |
| NaN | NaN | NaN |
| NaN | NaN | NaN |
| NaN | NaN | NaN |
| NaN | NaN | NaN |
| NaN | NaN | NaN |
| NaN | NaN | NaN |
| NaN | NaN | NaN |
| NaN | NaN | NaN |
| NaN | NaN | NaN |
| NaN | NaN | NaN |
| NaN | NaN | NaN |
| NaN | NaN | NaN |
| NaN | NaN | NaN |
| NaN | NaN | NaN |
| NaN | NaN | NaN |
| NaN | NaN | NaN |
| NaN | NaN | NaN |
| NaN | NaN | NaN |
| NaN | NaN | NaN |
| NaN | NaN | NaN |
| NaN | NaN | NaN |
| NaN | NaN | NaN |
| NaN | NaN | NaN |
| NaN | NaN | NaN |
| NaN | NaN | NaN |
| NaN | NaN | NaN |
| NaN | NaN | NaN |
| NaN | NaN | NaN |
| NaN | NaN | NaN |
| NaN | NaN | NaN |
| NaN | NaN | NaN |
| NaN | NaN | NaN |
| NaN | NaN | NaN |
| NaN | NaN | NaN |
| NaN | NaN | NaN |
| NaN | NaN | NaN |

| | | |
|---|---|---|
| NaN | NaN | NaN |
| NaN | NaN | NaN |
| NaN | NaN | NaN |
| NaN | NaN | NaN |
| NaN | NaN | NaN |
| NaN | NaN | NaN |
| NaN | NaN | NaN |
| NaN | NaN | NaN |
| NaN | NaN | NaN |
| NaN | NaN | NaN |
| NaN | NaN | NaN |
| NaN | NaN | NaN |
| NaN | NaN | NaN |
| NaN | NaN | NaN |
| NaN | NaN | NaN |
| NaN | NaN | NaN |
| NaN | NaN | NaN |
| NaN | NaN | NaN |
| NaN | NaN | NaN |
| NaN | NaN | NaN |
| NaN | NaN | NaN |
| NaN | NaN | NaN |
| NaN | NaN | NaN |
| NaN | NaN | NaN |
| NaN | NaN | NaN |
| NaN | NaN | NaN |
| NaN | NaN | NaN |
| NaN | NaN | NaN |
| NaN | NaN | NaN |
| NaN | NaN | NaN |
| NaN | NaN | NaN |
| NaN | NaN | NaN |
| NaN | NaN | NaN |
| NaN | NaN | NaN |
| NaN | NaN | NaN |
| NaN | NaN | NaN |
| NaN | NaN | NaN |
| NaN | NaN | NaN |
| NaN | NaN | NaN |
| NaN | NaN | NaN |
| NaN | NaN | NaN |
| NaN | NaN | NaN |
| NaN | NaN | NaN |
| NaN | NaN | NaN |
| NaN | NaN | NaN |
| NaN | NaN | NaN |
| NaN | NaN | NaN |
| NaN | NaN | NaN |

| | | |
|---|---|---|
| NaN | NaN | NaN |
| NaN | NaN | NaN |
| NaN | NaN | NaN |
| NaN | NaN | NaN |
| NaN | NaN | NaN |
| NaN | NaN | NaN |
| NaN | NaN | NaN |
| NaN | NaN | NaN |
| NaN | NaN | NaN |
| NaN | NaN | NaN |
| NaN | NaN | NaN |
| NaN | NaN | NaN |
| NaN | NaN | NaN |
| NaN | NaN | NaN |
| NaN | NaN | NaN |
| NaN | NaN | NaN |
| NaN | NaN | NaN |
| NaN | NaN | NaN |
| NaN | NaN | NaN |
| NaN | NaN | NaN |
| NaN | NaN | NaN |
| NaN | NaN | NaN |
| NaN | NaN | NaN |
| NaN | NaN | NaN |
| NaN | NaN | NaN |
| NaN | NaN | NaN |
| NaN | NaN | NaN |
| NaN | NaN | NaN |
| NaN | NaN | NaN |
| NaN | NaN | NaN |
| NaN | NaN | NaN |
| NaN | NaN | NaN |
| NaN | NaN | NaN |
| NaN | NaN | NaN |
| NaN | NaN | NaN |
| NaN | NaN | NaN |
| NaN | NaN | NaN |
| NaN | NaN | NaN |
| NaN | NaN | NaN |
| NaN | NaN | NaN |

| HIGH CAL | | ratio | pH LOC BKG | pH LYSO | LOW CAL | | ratio |
|---|---|---|---|---|---|---|---|
| NaN | NaN | NaN | | | NaN | NaN | NaN |
| NaN | NaN | NaN | | | NaN | NaN | NaN |
| NaN | NaN | NaN | | | NaN | NaN | NaN |
| NaN | NaN | NaN | | | NaN | NaN | NaN |
| NaN | NaN | NaN | | | NaN | NaN | NaN |
| NaN | NaN | NaN | | | NaN | NaN | NaN |
| NaN | NaN | NaN | | | NaN | NaN | NaN |
| NaN | NaN | NaN | | | NaN | NaN | NaN |
| NaN | NaN | NaN | | | NaN | NaN | NaN |
| NaN | NaN | NaN | | | NaN | NaN | NaN |
| NaN | NaN | NaN | | | NaN | NaN | NaN |
| NaN | NaN | NaN | | | NaN | NaN | NaN |
| NaN | NaN | NaN | | | NaN | NaN | NaN |
| NaN | NaN | NaN | | | NaN | NaN | NaN |
| NaN | NaN | NaN | | | NaN | NaN | NaN |
| NaN | NaN | NaN | | | NaN | NaN | NaN |
| NaN | NaN | NaN | | | NaN | NaN | NaN |
| NaN | NaN | NaN | | | NaN | NaN | NaN |
| NaN | NaN | NaN | | | NaN | NaN | NaN |
| NaN | NaN | NaN | | | NaN | NaN | NaN |
| NaN | NaN | NaN | | | NaN | NaN | NaN |
| NaN | NaN | NaN | | | NaN | NaN | NaN |
| NaN | NaN | NaN | | | NaN | NaN | NaN |
| NaN | NaN | NaN | | | NaN | NaN | NaN |
| NaN | NaN | NaN | | | NaN | NaN | NaN |
| NaN | NaN | NaN | | | NaN | NaN | NaN |
| NaN | NaN | NaN | | | NaN | NaN | NaN |
| NaN | NaN | NaN | | | NaN | NaN | NaN |
| NaN | NaN | NaN | | | NaN | NaN | NaN |
| NaN | NaN | NaN | | | NaN | NaN | NaN |
| NaN | NaN | NaN | | | NaN | NaN | NaN |
| NaN | NaN | NaN | | | NaN | NaN | NaN |
| NaN | NaN | NaN | | | NaN | NaN | NaN |
| NaN | NaN | NaN | | | NaN | NaN | NaN |
| NaN | NaN | NaN | | | NaN | NaN | NaN |
| NaN | NaN | NaN | | | NaN | NaN | NaN |
| NaN | NaN | NaN | | | NaN | NaN | NaN |
| NaN | NaN | NaN | | | NaN | NaN | NaN |
| NaN | NaN | NaN | | | NaN | NaN | NaN |
| NaN | NaN | NaN | | | NaN | NaN | NaN |
| NaN | NaN | NaN | | | NaN | NaN | NaN |
| NaN | NaN | NaN | | | NaN | NaN | NaN |
| NaN | NaN | NaN | | | NaN | NaN | NaN |
| NaN | NaN | NaN | | | NaN | NaN | NaN |
| NaN | NaN | NaN | | | NaN | NaN | NaN |
| NaN | NaN | NaN | | | NaN | NaN | NaN |
| NaN | NaN | NaN | | | NaN | NaN | NaN |

| | | | | | |
|---|---|---|---|---|---|
| NaN | NaN | NaN | NaN | NaN | NaN |
| NaN | NaN | NaN | NaN | NaN | NaN |
| NaN | NaN | NaN | NaN | NaN | NaN |
| NaN | NaN | NaN | NaN | NaN | NaN |
| NaN | NaN | NaN | NaN | NaN | NaN |
| NaN | NaN | NaN | NaN | NaN | NaN |
| NaN | NaN | NaN | NaN | NaN | NaN |
| NaN | NaN | NaN | NaN | NaN | NaN |
| NaN | NaN | NaN | NaN | NaN | NaN |
| NaN | NaN | NaN | NaN | NaN | NaN |
| NaN | NaN | NaN | NaN | NaN | NaN |
| NaN | NaN | NaN | NaN | NaN | NaN |
| NaN | NaN | NaN | NaN | NaN | NaN |
| NaN | NaN | NaN | NaN | NaN | NaN |
| NaN | NaN | NaN | NaN | NaN | NaN |
| NaN | NaN | NaN | NaN | NaN | NaN |
| NaN | NaN | NaN | NaN | NaN | NaN |
| NaN | NaN | NaN | NaN | NaN | NaN |
| NaN | NaN | NaN | NaN | NaN | NaN |
| NaN | NaN | NaN | NaN | NaN | NaN |
| NaN | NaN | NaN | NaN | NaN | NaN |
| NaN | NaN | NaN | NaN | NaN | NaN |
| NaN | NaN | NaN | NaN | NaN | NaN |
| NaN | NaN | NaN | NaN | NaN | NaN |
| NaN | NaN | NaN | NaN | NaN | NaN |
| NaN | NaN | NaN | NaN | NaN | NaN |
| NaN | NaN | NaN | NaN | NaN | NaN |
| NaN | NaN | NaN | NaN | NaN | NaN |
| NaN | NaN | NaN | NaN | NaN | NaN |
| NaN | NaN | NaN | NaN | NaN | NaN |
| NaN | NaN | NaN | NaN | NaN | NaN |
| NaN | NaN | NaN | NaN | NaN | NaN |
| NaN | NaN | NaN | NaN | NaN | NaN |
| NaN | NaN | NaN | NaN | NaN | NaN |
| NaN | NaN | NaN | NaN | NaN | NaN |
| NaN | NaN | NaN | NaN | NaN | NaN |
| NaN | NaN | NaN | NaN | NaN | NaN |
| NaN | NaN | NaN | NaN | NaN | NaN |
| NaN | NaN | NaN | NaN | NaN | NaN |
| NaN | NaN | NaN | NaN | NaN | NaN |
| NaN | NaN | NaN | NaN | NaN | NaN |
| NaN | NaN | NaN | NaN | NaN | NaN |
| NaN | NaN | NaN | NaN | NaN | NaN |
| NaN | NaN | NaN | NaN | NaN | NaN |
| NaN | NaN | NaN | NaN | NaN | NaN |
| NaN | NaN | NaN | NaN | NaN | NaN |
| NaN | NaN | NaN | NaN | NaN | NaN |
| NaN | NaN | NaN | NaN | NaN | NaN |

| | | | | | |
|---|---|---|---|---|---|
| NaN | NaN | NaN | NaN | NaN | NaN |
| NaN | NaN | NaN | NaN | NaN | NaN |
| NaN | NaN | NaN | NaN | NaN | NaN |
| NaN | NaN | NaN | NaN | NaN | NaN |
| NaN | NaN | NaN | NaN | NaN | NaN |
| NaN | NaN | NaN | NaN | NaN | NaN |
| NaN | NaN | NaN | NaN | NaN | NaN |
| NaN | NaN | NaN | NaN | NaN | NaN |
| NaN | NaN | NaN | NaN | NaN | NaN |
| NaN | NaN | NaN | NaN | NaN | NaN |
| NaN | NaN | NaN | NaN | NaN | NaN |
| NaN | NaN | NaN | NaN | NaN | NaN |
| NaN | NaN | NaN | NaN | NaN | NaN |
| NaN | NaN | NaN | NaN | NaN | NaN |
| NaN | NaN | NaN | NaN | NaN | NaN |
| NaN | NaN | NaN | NaN | NaN | NaN |
| NaN | NaN | NaN | NaN | NaN | NaN |
| NaN | NaN | NaN | NaN | NaN | NaN |
| NaN | NaN | NaN | NaN | NaN | NaN |
| NaN | NaN | NaN | NaN | NaN | NaN |
| NaN | NaN | NaN | NaN | NaN | NaN |
| NaN | NaN | NaN | NaN | NaN | NaN |
| NaN | NaN | NaN | NaN | NaN | NaN |
| NaN | NaN | NaN | NaN | NaN | NaN |
| NaN | NaN | NaN | NaN | NaN | NaN |
| NaN | NaN | NaN | NaN | NaN | NaN |
| NaN | NaN | NaN | NaN | NaN | NaN |
| NaN | NaN | NaN | NaN | NaN | NaN |
| NaN | NaN | NaN | NaN | NaN | NaN |
| NaN | NaN | NaN | NaN | NaN | NaN |
| NaN | NaN | NaN | NaN | NaN | NaN |
| NaN | NaN | NaN | NaN | NaN | NaN |
| NaN | NaN | NaN | NaN | NaN | NaN |
| NaN | NaN | NaN | NaN | NaN | NaN |
| NaN | NaN | NaN | NaN | NaN | NaN |
| NaN | NaN | NaN | NaN | NaN | NaN |
| NaN | NaN | NaN | NaN | NaN | NaN |
| NaN | NaN | NaN | NaN | NaN | NaN |
| NaN | NaN | NaN | NaN | NaN | NaN |
| NaN | NaN | NaN | NaN | NaN | NaN |
| NaN | NaN | NaN | NaN | NaN | NaN |
| NaN | NaN | NaN | NaN | NaN | NaN |
| NaN | NaN | NaN | NaN | NaN | NaN |
| NaN | NaN | NaN | NaN | NaN | NaN |
| NaN | NaN | NaN | NaN | NaN | NaN |
| NaN | NaN | NaN | NaN | NaN | NaN |
| NaN | NaN | NaN | NaN | NaN | NaN |
| NaN | NaN | NaN | NaN | NaN | NaN |

| | | | | | |
|---|---|---|---|---|---|
| NaN | NaN | NaN | NaN | NaN | NaN |
| NaN | NaN | NaN | NaN | NaN | NaN |
| NaN | NaN | NaN | NaN | NaN | NaN |
| NaN | NaN | NaN | NaN | NaN | NaN |
| NaN | NaN | NaN | NaN | NaN | NaN |
| NaN | NaN | NaN | NaN | NaN | NaN |
| NaN | NaN | NaN | NaN | NaN | NaN |
| NaN | NaN | NaN | NaN | NaN | NaN |
| NaN | NaN | NaN | NaN | NaN | NaN |
| NaN | NaN | NaN | NaN | NaN | NaN |
| NaN | NaN | NaN | NaN | NaN | NaN |
| NaN | NaN | NaN | NaN | NaN | NaN |
| NaN | NaN | NaN | NaN | NaN | NaN |
| NaN | NaN | NaN | NaN | NaN | NaN |
| NaN | NaN | NaN | NaN | NaN | NaN |
| NaN | NaN | NaN | NaN | NaN | NaN |
| NaN | NaN | NaN | NaN | NaN | NaN |
| NaN | NaN | NaN | NaN | NaN | NaN |
| NaN | NaN | NaN | NaN | NaN | NaN |
| NaN | NaN | NaN | NaN | NaN | NaN |
| NaN | NaN | NaN | NaN | NaN | NaN |
| NaN | NaN | NaN | NaN | NaN | NaN |
| NaN | NaN | NaN | NaN | NaN | NaN |
| NaN | NaN | NaN | NaN | NaN | NaN |
| NaN | NaN | NaN | NaN | NaN | NaN |
| NaN | NaN | NaN | NaN | NaN | NaN |
| NaN | NaN | NaN | NaN | NaN | NaN |
| NaN | NaN | NaN | NaN | NaN | NaN |
| NaN | NaN | NaN | NaN | NaN | NaN |
| NaN | NaN | NaN | NaN | NaN | NaN |
| NaN | NaN | NaN | NaN | NaN | NaN |
| NaN | NaN | NaN | NaN | NaN | NaN |
| NaN | NaN | NaN | NaN | NaN | NaN |
| NaN | NaN | NaN | NaN | NaN | NaN |
| NaN | NaN | NaN | NaN | NaN | NaN |
| NaN | NaN | NaN | NaN | NaN | NaN |
| NaN | NaN | NaN | NaN | NaN | NaN |
| NaN | NaN | NaN | NaN | NaN | NaN |
| NaN | NaN | NaN | NaN | NaN | NaN |
| NaN | NaN | NaN | NaN | NaN | NaN |
| NaN | NaN | NaN | NaN | NaN | NaN |
| NaN | NaN | NaN | NaN | NaN | NaN |
| NaN | NaN | NaN | NaN | NaN | NaN |
| NaN | NaN | NaN | NaN | NaN | NaN |
| NaN | NaN | NaN | NaN | NaN | NaN |
| NaN | NaN | NaN | NaN | NaN | NaN |
| NaN | NaN | NaN | NaN | NaN | NaN |
| NaN | NaN | NaN | NaN | NaN | NaN |

| | | | | | | | |
|---|---|---|---|---|---|---|---|
| NaN | NaN | NaN | | | NaN | NaN | NaN |
| NaN | NaN | NaN | | | NaN | NaN | NaN |
| NaN | NaN | NaN | | | NaN | NaN | NaN |
| NaN | NaN | NaN | | | NaN | NaN | NaN |
| NaN | NaN | NaN | | | NaN | NaN | NaN |
| NaN | NaN | NaN | | | NaN | NaN | NaN |
| NaN | NaN | NaN | | | NaN | NaN | NaN |
| NaN | NaN | NaN | | | NaN | NaN | NaN |
| NaN | NaN | NaN | | | NaN | NaN | NaN |
| NaN | NaN | NaN | | | NaN | NaN | NaN |
| NaN | NaN | NaN | | | NaN | NaN | NaN |
| NaN | NaN | NaN | | | NaN | NaN | NaN |
| NaN | NaN | NaN | | | NaN | NaN | NaN |
| NaN | NaN | NaN | | | NaN | NaN | NaN |
| NaN | NaN | NaN | | | NaN | NaN | NaN |
| NaN | NaN | NaN | | | NaN | NaN | NaN |
| NaN | NaN | NaN | | | NaN | NaN | NaN |
| NaN | NaN | NaN | | | NaN | NaN | NaN |
| NaN | NaN | NaN | | | NaN | NaN | NaN |
| NaN | NaN | NaN | | | NaN | NaN | NaN |
| 43.69 | 188.22 | 0.232122 | 4.610581 | 4.633955 | NaN | NaN | NaN |
| 194.53 | 274.46 | 0.708774 | 5.903202 | 5.890583 | NaN | NaN | NaN |
| 231.28 | 174.99 | 1.321676 | 7.565318 | 7.506415 | NaN | NaN | NaN |
| 98.41 | 259.43 | 0.379332 | 5.009796 | 5.022054 | NaN | NaN | NaN |
| 492.49 | 604.35 | 0.814909 | 6.191027 | 6.170393 | NaN | NaN | NaN |
| 29.44 | 138.3 | 0.212871 | 4.558374 | 4.583202 | NaN | NaN | NaN |
| 169.64 | 177.68 | 0.95475 | 6.570261 | 6.539066 | NaN | NaN | NaN |
| 219.54 | 216.24 | 1.015261 | 6.734358 | 6.698595 | NaN | NaN | NaN |
| 365.19 | 360.45 | 1.01315 | 6.728635 | 6.69303 | NaN | NaN | NaN |
| 93.49 | 325.28 | 0.287414 | 4.760526 | 4.779725 | NaN | NaN | NaN |
| 187.12 | 278.76 | 0.671258 | 5.801466 | 5.791679 | NaN | NaN | NaN |
| 302.78 | 249.29 | 1.214569 | 7.274859 | 7.224044 | NaN | NaN | NaN |
| 151.17 | 181.38 | 0.833444 | 6.241292 | 6.219258 | NaN | NaN | NaN |
| 357.04 | 196.05 | 1.821168 | 8.919881 | 8.823259 | NaN | NaN | NaN |
| 255.26 | 242.42 | 1.052966 | 6.83661 | 6.797999 | NaN | NaN | NaN |
| 173.85 | 151.91 | 1.144428 | 7.084643 | 7.039125 | NaN | NaN | NaN |
| 500.18 | 210.67 | 2.374235 | 10.41973 | 10.28134 | NaN | NaN | NaN |
| 310.88 | 344.15 | 0.903327 | 6.430808 | 6.403496 | NaN | NaN | NaN |
| 137.39 | 182.75 | 0.751792 | 6.019863 | 6.003995 | NaN | NaN | NaN |
| 40.03 | 151.05 | 0.265012 | 4.699774 | 4.720664 | NaN | NaN | NaN |
| 555.1 | 453.84 | 1.223118 | 7.298042 | 7.246583 | NaN | NaN | NaN |
| 215.62 | 216.55 | 0.995705 | 6.681326 | 6.647039 | NaN | NaN | NaN |
| 231.49 | 216.19 | 1.070771 | 6.884895 | 6.84494 | NaN | NaN | NaN |
| 101.48 | 107.41 | 0.944791 | 6.543253 | 6.51281 | NaN | NaN | NaN |
| 819.53 | 385.45 | 2.126164 | 9.746993 | 9.62734 | NaN | NaN | NaN |
| 113.81 | 243.33 | 0.467719 | 5.249491 | 5.255074 | NaN | NaN | NaN |
| 511.02 | 332.2 | 1.53829 | 8.15275 | 8.07749 | NaN | NaN | NaN |
| 379.4 | 602.35 | 0.629866 | 5.689215 | 5.682554 | NaN | NaN | NaN |

| | | | | | | | |
|---|---|---|---|---|---|---|---|
| 396.32 | 324.29 | 1.222116 | 7.295324 | 7.24394 | NaN | NaN | NaN |
| 572.48 | 516.35 | 1.108705 | 6.987768 | 6.944948 | NaN | NaN | NaN |
| 382.08 | 427.43 | 0.893901 | 6.405245 | 6.378645 | NaN | NaN | NaN |
| 514.81 | 365.01 | 1.4104 | 7.805927 | 7.740325 | NaN | NaN | NaN |
| 208.14 | 202.27 | 1.029021 | 6.771673 | 6.73487 | NaN | NaN | NaN |
| 167.64 | 251.26 | 0.667197 | 5.790452 | 5.780972 | NaN | NaN | NaN |
| 82.47 | 120.05 | 0.686964 | 5.844057 | 5.833084 | NaN | NaN | NaN |
| 127.75 | 420.34 | 0.303921 | 4.80529 | 4.823243 | NaN | NaN | NaN |
| -6.73 | 179.66 | -0.03746 | 3.879509 | 3.92324 | NaN | NaN | NaN |
| 163.74 | 138.45 | 1.182665 | 7.188339 | 7.139934 | NaN | NaN | NaN |
| 47.95 | 275.33 | 0.174155 | 4.453381 | 4.481132 | NaN | NaN | NaN |
| 724.11 | 438.48 | 1.651409 | 8.459516 | 8.375714 | NaN | NaN | NaN |
| 779.34 | 673.4 | 1.157321 | 7.119608 | 7.073117 | NaN | NaN | NaN |
| 117.49 | 129.2 | 0.909365 | 6.447183 | 6.419415 | NaN | NaN | NaN |
| 327.3 | 239.13 | 1.368712 | 7.692874 | 7.630419 | NaN | NaN | NaN |
| 190.2 | 182.14 | 1.044252 | 6.812978 | 6.775025 | NaN | NaN | NaN |
| 508.13 | 470 | 1.081128 | 6.912981 | 6.872244 | NaN | NaN | NaN |
| 156.18 | 178.57 | 0.874615 | 6.352944 | 6.327801 | NaN | NaN | NaN |
| 166.55 | 331.74 | 0.50205 | 5.342592 | 5.345583 | NaN | NaN | NaN |
| 221.86 | 134.33 | 1.651604 | 8.460044 | 8.376228 | NaN | NaN | NaN |
| 258 | 251.25 | 1.026866 | 6.765829 | 6.729189 | NaN | NaN | NaN |
| -7.16 | 133.41 | -0.05367 | 3.83555 | 3.880506 | NaN | NaN | NaN |
| 422.75 | 247.56 | 1.707667 | 8.612079 | 8.524029 | NaN | NaN | NaN |
| 241.53 | 238.41 | 1.013087 | 6.728462 | 6.692863 | NaN | NaN | NaN |
| -26.73 | 206.29 | -0.12957 | 3.629703 | 3.680391 | NaN | NaN | NaN |
| 67.32 | 283.4 | 0.237544 | 4.625285 | 4.64825 | NaN | NaN | NaN |
| 479.79 | 893.47 | 0.536996 | 5.437363 | 5.437715 | NaN | NaN | NaN |
| 250.01 | 335.58 | 0.745009 | 6.001467 | 5.986111 | NaN | NaN | NaN |
| 307.6 | 348.25 | 0.883274 | 6.376425 | 6.350628 | NaN | NaN | NaN |
| 17.03 | 176.14 | 0.096684 | 4.243291 | 4.276893 | NaN | NaN | NaN |
| 53.26 | 261.81 | 0.20343 | 4.532772 | 4.558313 | NaN | NaN | NaN |
| 338.03 | 376.93 | 0.896798 | 6.413101 | 6.386283 | NaN | NaN | NaN |
| 366.85 | 380.53 | 0.96405 | 6.595481 | 6.563585 | NaN | NaN | NaN |
| 326.61 | 302.62 | 1.079274 | 6.907955 | 6.867358 | NaN | NaN | NaN |
| 107.47 | 102.51 | 1.048386 | 6.824188 | 6.785923 | NaN | NaN | NaN |
| 270.39 | 285.39 | 0.94744 | 6.550437 | 6.519795 | NaN | NaN | NaN |
| 17.85 | 122.53 | 0.145679 | 4.376157 | 4.406059 | NaN | NaN | NaN |
| 210.85 | 297.35 | 0.709097 | 5.904079 | 5.891435 | NaN | NaN | NaN |
| 340.4 | 267.16 | 1.274143 | 7.436415 | 7.381102 | NaN | NaN | NaN |
| -7.5 | 203.36 | -0.03688 | 3.881079 | 3.924767 | NaN | NaN | NaN |
| 104.79 | 229.87 | 0.455866 | 5.217349 | 5.223827 | NaN | NaN | NaN |
| 339.85 | 243.16 | 1.397639 | 7.771322 | 7.706684 | NaN | NaN | NaN |
| 412.57 | 412.8 | 0.999443 | 6.691462 | 6.656893 | NaN | NaN | NaN |
| 318.34 | 223.3 | 1.425616 | 7.847191 | 7.78044 | NaN | NaN | NaN |
| 306.38 | 488.34 | 0.627391 | 5.682502 | 5.676028 | NaN | NaN | NaN |
| 235.73 | 292.93 | 0.804732 | 6.163428 | 6.143563 | NaN | NaN | NaN |
| 330.23 | 411.17 | 0.803147 | 6.159132 | 6.139386 | NaN | NaN | NaN |
| 227.63 | 331.43 | 0.686812 | 5.843644 | 5.832683 | NaN | NaN | NaN |

| | | | | | | | |
|---|---|---|---|---|---|---|---|
| 240.56 | 290.71 | 0.827491 | 6.22515 | 6.203566 | NaN | NaN | NaN |
| 561.09 | 487.86 | 1.150105 | 7.100038 | 7.054092 | NaN | NaN | NaN |
| 562.46 | 376.47 | 1.494037 | 8.03274 | 7.960822 | NaN | NaN | NaN |
| 200.78 | 497.11 | 0.403895 | 5.076407 | 5.08681 | NaN | NaN | NaN |
| 358.12 | 292.59 | 1.223965 | 7.300339 | 7.248815 | NaN | NaN | NaN |
| 250.37 | 182.71 | 1.370314 | 7.697218 | 7.634643 | NaN | NaN | NaN |
| 548.11 | 295.89 | 1.852411 | 9.004609 | 8.905628 | NaN | NaN | NaN |
| -8.76 | 212.55 | -0.04121 | 3.869328 | 3.913343 | NaN | NaN | NaN |
| 144.21 | 206.68 | 0.697745 | 5.873295 | 5.861508 | NaN | NaN | NaN |
| 27.48 | 180.91 | 0.151899 | 4.393025 | 4.422458 | NaN | NaN | NaN |
| 414.04 | 238.44 | 1.736454 | 8.690145 | 8.599921 | NaN | NaN | NaN |
| 361.03 | 239.35 | 1.508377 | 8.071629 | 7.998628 | NaN | NaN | NaN |
| 1237.14 | 855.06 | 1.446846 | 7.904764 | 7.83641 | NaN | NaN | NaN |
| 979.69 | 603.18 | 1.624208 | 8.38575 | 8.304002 | NaN | NaN | NaN |
| 9.01 | 190.24 | 0.047361 | 4.109532 | 4.146859 | NaN | NaN | NaN |
| 216.58 | 231.18 | 0.936846 | 6.521706 | 6.491864 | NaN | NaN | NaN |
| 251.45 | 220.68 | 1.139433 | 7.071097 | 7.025957 | NaN | NaN | NaN |
| 88.28 | 168.54 | 0.523793 | 5.401556 | 5.402905 | NaN | NaN | NaN |
| 79.8 | 248.28 | 0.321411 | 4.852723 | 4.869355 | NaN | NaN | NaN |
| 453.82 | 246.06 | 1.844347 | 8.982739 | 8.884367 | NaN | NaN | NaN |
| 493.64 | 284.65 | 1.7342 | 8.684034 | 8.59398 | NaN | NaN | NaN |
| 766.55 | 188.34 | 4.070033 | 15.01853 | 14.75209 | NaN | NaN | NaN |
| 484.66 | 462.71 | 1.047438 | 6.821619 | 6.783425 | NaN | NaN | NaN |
| 430.65 | 315.02 | 1.367056 | 7.688384 | 7.626055 | NaN | NaN | NaN |
| 5.72 | 141.02 | 0.040562 | 4.091093 | 4.128933 | NaN | NaN | NaN |
| NaN | NaN | NaN | | | NaN | NaN | NaN |
| NaN | NaN | NaN | | | NaN | NaN | NaN |
| NaN | NaN | NaN | | | NaN | NaN | NaN |
| NaN | NaN | NaN | | | NaN | NaN | NaN |
| NaN | NaN | NaN | | | NaN | NaN | NaN |
| NaN | NaN | NaN | | | NaN | NaN | NaN |
| NaN | NaN | NaN | | | NaN | NaN | NaN |
| NaN | NaN | NaN | | | NaN | NaN | NaN |
| NaN | NaN | NaN | | | NaN | NaN | NaN |
| NaN | NaN | NaN | | | NaN | NaN | NaN |
| NaN | NaN | NaN | | | NaN | NaN | NaN |
| NaN | NaN | NaN | | | NaN | NaN | NaN |
| NaN | NaN | NaN | | | NaN | NaN | NaN |
| NaN | NaN | NaN | | | NaN | NaN | NaN |
| NaN | NaN | NaN | | | NaN | NaN | NaN |
| NaN | NaN | NaN | | | NaN | NaN | NaN |
| NaN | NaN | NaN | | | NaN | NaN | NaN |
| NaN | NaN | NaN | | | NaN | NaN | NaN |
| NaN | NaN | NaN | | | NaN | NaN | NaN |
| NaN | NaN | NaN | | | NaN | NaN | NaN |
| NaN | NaN | NaN | | | NaN | NaN | NaN |
| NaN | NaN | NaN | | | NaN | NaN | NaN |
| NaN | NaN | NaN | | | NaN | NaN | NaN |

| | | | | | |
|---|---|---|---|---|---|
| NaN | NaN | NaN | NaN | NaN | NaN |
| NaN | NaN | NaN | NaN | NaN | NaN |
| NaN | NaN | NaN | NaN | NaN | NaN |
| NaN | NaN | NaN | NaN | NaN | NaN |
| NaN | NaN | NaN | NaN | NaN | NaN |
| NaN | NaN | NaN | NaN | NaN | NaN |
| NaN | NaN | NaN | NaN | NaN | NaN |
| NaN | NaN | NaN | NaN | NaN | NaN |
| NaN | NaN | NaN | NaN | NaN | NaN |
| NaN | NaN | NaN | NaN | NaN | NaN |
| NaN | NaN | NaN | NaN | NaN | NaN |
| NaN | NaN | NaN | NaN | NaN | NaN |
| NaN | NaN | NaN | NaN | NaN | NaN |
| NaN | NaN | NaN | NaN | NaN | NaN |
| NaN | NaN | NaN | NaN | NaN | NaN |
| NaN | NaN | NaN | NaN | NaN | NaN |
| NaN | NaN | NaN | NaN | NaN | NaN |
| NaN | NaN | NaN | NaN | NaN | NaN |
| NaN | NaN | NaN | NaN | NaN | NaN |
| NaN | NaN | NaN | NaN | NaN | NaN |
| NaN | NaN | NaN | NaN | NaN | NaN |
| NaN | NaN | NaN | NaN | NaN | NaN |
| NaN | NaN | NaN | NaN | NaN | NaN |
| NaN | NaN | NaN | NaN | NaN | NaN |
| NaN | NaN | NaN | NaN | NaN | NaN |
| NaN | NaN | NaN | NaN | NaN | NaN |
| NaN | NaN | NaN | NaN | NaN | NaN |
| NaN | NaN | NaN | NaN | NaN | NaN |
| NaN | NaN | NaN | NaN | NaN | NaN |
| NaN | NaN | NaN | NaN | NaN | NaN |
| NaN | NaN | NaN | NaN | NaN | NaN |
| NaN | NaN | NaN | NaN | NaN | NaN |
| NaN | NaN | NaN | NaN | NaN | NaN |
| NaN | NaN | NaN | NaN | NaN | NaN |
| NaN | NaN | NaN | NaN | NaN | NaN |
| NaN | NaN | NaN | NaN | NaN | NaN |
| NaN | NaN | NaN | NaN | NaN | NaN |
| NaN | NaN | NaN | NaN | NaN | NaN |
| NaN | NaN | NaN | NaN | NaN | NaN |
| NaN | NaN | NaN | NaN | NaN | NaN |
| NaN | NaN | NaN | NaN | NaN | NaN |
| NaN | NaN | NaN | NaN | NaN | NaN |
| NaN | NaN | NaN | NaN | NaN | NaN |
| NaN | NaN | NaN | NaN | NaN | NaN |
| NaN | NaN | NaN | NaN | NaN | NaN |
| NaN | NaN | NaN | NaN | NaN | NaN |
| NaN | NaN | NaN | NaN | NaN | NaN |
| NaN | NaN | NaN | NaN | NaN | NaN |

| | | | | | |
|---|---|---|---|---|---|
| NaN | NaN | NaN | NaN | NaN | NaN |
| NaN | NaN | NaN | NaN | NaN | NaN |
| NaN | NaN | NaN | NaN | NaN | NaN |
| NaN | NaN | NaN | NaN | NaN | NaN |
| NaN | NaN | NaN | NaN | NaN | NaN |
| NaN | NaN | NaN | NaN | NaN | NaN |
| NaN | NaN | NaN | NaN | NaN | NaN |
| NaN | NaN | NaN | NaN | NaN | NaN |
| NaN | NaN | NaN | NaN | NaN | NaN |
| NaN | NaN | NaN | NaN | NaN | NaN |
| NaN | NaN | NaN | NaN | NaN | NaN |
| NaN | NaN | NaN | NaN | NaN | NaN |
| NaN | NaN | NaN | NaN | NaN | NaN |
| NaN | NaN | NaN | NaN | NaN | NaN |
| NaN | NaN | NaN | NaN | NaN | NaN |
| NaN | NaN | NaN | NaN | NaN | NaN |
| NaN | NaN | NaN | NaN | NaN | NaN |
| NaN | NaN | NaN | NaN | NaN | NaN |
| NaN | NaN | NaN | NaN | NaN | NaN |
| NaN | NaN | NaN | NaN | NaN | NaN |
| NaN | NaN | NaN | NaN | NaN | NaN |
| NaN | NaN | NaN | NaN | NaN | NaN |
| NaN | NaN | NaN | NaN | NaN | NaN |
| NaN | NaN | NaN | NaN | NaN | NaN |
| NaN | NaN | NaN | NaN | NaN | NaN |
| NaN | NaN | NaN | NaN | NaN | NaN |
| NaN | NaN | NaN | NaN | NaN | NaN |
| NaN | NaN | NaN | NaN | NaN | NaN |
| NaN | NaN | NaN | NaN | NaN | NaN |
| NaN | NaN | NaN | NaN | NaN | NaN |
| NaN | NaN | NaN | NaN | NaN | NaN |
| NaN | NaN | NaN | 5.17 | 172.26 | 0.030013 |
| NaN | NaN | NaN | 58.98 | 293 | 0.201297 |
| NaN | NaN | NaN | -20.23 | 85.44 | -0.23677 |
| NaN | NaN | NaN | 7.07 | 245.21 | 0.028832 |
| NaN | NaN | NaN | 24.04 | 147.39 | 0.163105 |
| NaN | NaN | NaN | 55.81 | 338.13 | 0.165055 |
| NaN | NaN | NaN | 19.99 | 291.53 | 0.068569 |
| NaN | NaN | NaN | 52.98 | 237.79 | 0.222802 |
| NaN | NaN | NaN | 61.02 | 213.04 | 0.286425 |
| NaN | NaN | NaN | 27.18 | 166.58 | 0.163165 |
| NaN | NaN | NaN | 39.57 | 402.04 | 0.098423 |
| NaN | NaN | NaN | 44.24 | 223.26 | 0.198155 |
| NaN | NaN | NaN | 55.32 | 269.36 | 0.205376 |
| NaN | NaN | NaN | 51.49 | 418 | 0.123182 |
| NaN | NaN | NaN | 47.98 | 159.7 | 0.300438 |
| NaN | NaN | NaN | 68.54 | 347.24 | 0.197385 |
| NaN | NaN | NaN | 72.13 | 313.76 | 0.229889 |

| | | | | | |
|---|---|---|---|---|---|
| NaN | NaN | NaN | 36.03 | 172.9 | 0.208386 |
| NaN | NaN | NaN | 50.09 | 275.35 | 0.181914 |
| NaN | NaN | NaN | -1.61 | 168.79 | -0.00954 |
| NaN | NaN | NaN | 45.09 | 209.43 | 0.215299 |
| NaN | NaN | NaN | 71.09 | 445.7 | 0.159502 |
| NaN | NaN | NaN | 3.11 | 229.13 | 0.013573 |
| NaN | NaN | NaN | 33.83 | 160.08 | 0.211332 |
| NaN | NaN | NaN | 67.94 | 392.3 | 0.173184 |
| NaN | NaN | NaN | 8.68 | 199.21 | 0.043572 |
| NaN | NaN | NaN | 56.75 | 499.77 | 0.113552 |
| NaN | NaN | NaN | 5.18 | 201.82 | 0.025666 |
| NaN | NaN | NaN | 44.32 | 252.93 | 0.175226 |
| NaN | NaN | NaN | 58.43 | 301.02 | 0.194107 |
| NaN | NaN | NaN | 19.77 | 216.96 | 0.091123 |
| NaN | NaN | NaN | 58.48 | 365.39 | 0.160048 |
| NaN | NaN | NaN | 65.12 | 330.98 | 0.196749 |
| NaN | NaN | NaN | 34.43 | 310.96 | 0.110722 |
| NaN | NaN | NaN | 51.54 | 670.56 | 0.076861 |
| NaN | NaN | NaN | 8.25 | 173.86 | 0.047452 |
| NaN | NaN | NaN | 2.41 | 285.27 | 0.008448 |
| NaN | NaN | NaN | 0.74 | 173.83 | 0.004257 |
| NaN | NaN | NaN | 1.52 | 176.62 | 0.008606 |
| NaN | NaN | NaN | 44.84 | 202.7 | 0.221214 |
| NaN | NaN | NaN | 32.06 | 337.6 | 0.094964 |
| NaN | NaN | NaN | 55.66 | 460.36 | 0.120905 |
| NaN | NaN | NaN | 50.88 | 290.02 | 0.175436 |
| NaN | NaN | NaN | 49.44 | 347.83 | 0.142138 |
| NaN | NaN | NaN | 90.93 | 450.19 | 0.201981 |
| NaN | NaN | NaN | -0.53 | 252.52 | -0.0021 |
| NaN | NaN | NaN | 56.44 | 361.21 | 0.156253 |
| NaN | NaN | NaN | 42.35 | 273.35 | 0.15493 |
| NaN | NaN | NaN | 58.26 | 243.52 | 0.239241 |
| NaN | NaN | NaN | 66.41 | 459.78 | 0.144439 |
| NaN | NaN | NaN | 4.13 | 176.44 | 0.023407 |
| NaN | NaN | NaN | 52.05 | 246.81 | 0.210891 |
| NaN | NaN | NaN | 71.42 | 341.52 | 0.209124 |
| NaN | NaN | NaN | -2.08 | 280.62 | -0.00741 |
| NaN | NaN | NaN | 36.67 | 197.42 | 0.185746 |
| NaN | NaN | NaN | 9.76 | 214.63 | 0.045474 |
| NaN | NaN | NaN | 4.17 | 233.75 | 0.01784 |
| NaN | NaN | NaN | 16.36 | 187.43 | 0.087286 |
| NaN | NaN | NaN | 67.2 | 271.74 | 0.247295 |
| NaN | NaN | NaN | 133.35 | 681.68 | 0.19562 |
| NaN | NaN | NaN | 56.09 | 241.23 | 0.232517 |
| NaN | NaN | NaN | 87.93 | 364.24 | 0.241407 |
| NaN | NaN | NaN | -4.2 | 246.53 | -0.01704 |
| NaN | NaN | NaN | 9.95 | 147.14 | 0.067623 |
| NaN | NaN | NaN | 87.65 | 344.9 | 0.254132 |

| | | | | | |
|---|---|---|---|---|---|
| NaN | NaN | NaN | 1.94 | 222.16 | 0.008732 |
| NaN | NaN | NaN | 42.7 | 168.04 | 0.254106 |
| NaN | NaN | NaN | 8.63 | 238.02 | 0.036257 |
| NaN | NaN | NaN | 43.56 | 144.47 | 0.301516 |
| NaN | NaN | NaN | 64.39 | 359.43 | 0.179145 |
| NaN | NaN | NaN | 24.13 | 174.56 | 0.138233 |
| NaN | NaN | NaN | -1.71 | 165.66 | -0.01032 |
| NaN | NaN | NaN | 9.62 | 185.54 | 0.051849 |
| NaN | NaN | NaN | 100.39 | 575.96 | 0.1743 |
| NaN | NaN | NaN | NaN | NaN | NaN |
| NaN | NaN | NaN | NaN | NaN | NaN |
| NaN | NaN | NaN | NaN | NaN | NaN |
| NaN | NaN | NaN | NaN | NaN | NaN |
| NaN | NaN | NaN | NaN | NaN | NaN |
| NaN | NaN | NaN | NaN | NaN | NaN |
| NaN | NaN | NaN | NaN | NaN | NaN |
| NaN | NaN | NaN | NaN | NaN | NaN |
| NaN | NaN | NaN | NaN | NaN | NaN |
| NaN | NaN | NaN | NaN | NaN | NaN |
| NaN | NaN | NaN | NaN | NaN | NaN |
| NaN | NaN | NaN | NaN | NaN | NaN |
| NaN | NaN | NaN | NaN | NaN | NaN |
| NaN | NaN | NaN | NaN | NaN | NaN |
| NaN | NaN | NaN | NaN | NaN | NaN |
| NaN | NaN | NaN | NaN | NaN | NaN |
| NaN | NaN | NaN | NaN | NaN | NaN |
| NaN | NaN | NaN | NaN | NaN | NaN |
| NaN | NaN | NaN | NaN | NaN | NaN |
| NaN | NaN | NaN | NaN | NaN | NaN |
| NaN | NaN | NaN | NaN | NaN | NaN |
| NaN | NaN | NaN | NaN | NaN | NaN |
| NaN | NaN | NaN | NaN | NaN | NaN |
| NaN | NaN | NaN | NaN | NaN | NaN |
| NaN | NaN | NaN | NaN | NaN | NaN |
| NaN | NaN | NaN | NaN | NaN | NaN |
| NaN | NaN | NaN | NaN | NaN | NaN |
| NaN | NaN | NaN | NaN | NaN | NaN |
| NaN | NaN | NaN | NaN | NaN | NaN |
| NaN | NaN | NaN | NaN | NaN | NaN |
| NaN | NaN | NaN | NaN | NaN | NaN |
| NaN | NaN | NaN | NaN | NaN | NaN |
| NaN | NaN | NaN | NaN | NaN | NaN |
| NaN | NaN | NaN | NaN | NaN | NaN |
| NaN | NaN | NaN | NaN | NaN | NaN |
| NaN | NaN | NaN | NaN | NaN | NaN |
| NaN | NaN | NaN | NaN | NaN | NaN |
| NaN | NaN | NaN | NaN | NaN | NaN |
| NaN | NaN | NaN | NaN | NaN | NaN |

| | | | | | |
|---|---|---|---|---|---|
| NaN | NaN | NaN | NaN | NaN | NaN |
| NaN | NaN | NaN | NaN | NaN | NaN |
| NaN | NaN | NaN | NaN | NaN | NaN |
| NaN | NaN | NaN | NaN | NaN | NaN |
| NaN | NaN | NaN | NaN | NaN | NaN |
| NaN | NaN | NaN | NaN | NaN | NaN |
| NaN | NaN | NaN | NaN | NaN | NaN |
| NaN | NaN | NaN | NaN | NaN | NaN |
| NaN | NaN | NaN | NaN | NaN | NaN |
| NaN | NaN | NaN | NaN | NaN | NaN |
| NaN | NaN | NaN | NaN | NaN | NaN |
| NaN | NaN | NaN | NaN | NaN | NaN |
| NaN | NaN | NaN | NaN | NaN | NaN |
| NaN | NaN | NaN | NaN | NaN | NaN |
| NaN | NaN | NaN | NaN | NaN | NaN |
| NaN | NaN | NaN | NaN | NaN | NaN |
| NaN | NaN | NaN | NaN | NaN | NaN |
| NaN | NaN | NaN | NaN | NaN | NaN |
| NaN | NaN | NaN | NaN | NaN | NaN |
| NaN | NaN | NaN | NaN | NaN | NaN |
| NaN | NaN | NaN | NaN | NaN | NaN |
| NaN | NaN | NaN | NaN | NaN | NaN |
| NaN | NaN | NaN | NaN | NaN | NaN |
| NaN | NaN | NaN | NaN | NaN | NaN |
| NaN | NaN | NaN | NaN | NaN | NaN |
| NaN | NaN | NaN | NaN | NaN | NaN |
| NaN | NaN | NaN | NaN | NaN | NaN |
| NaN | NaN | NaN | NaN | NaN | NaN |
| NaN | NaN | NaN | NaN | NaN | NaN |
| NaN | NaN | NaN | NaN | NaN | NaN |
| NaN | NaN | NaN | NaN | NaN | NaN |
| NaN | NaN | NaN | NaN | NaN | NaN |
| NaN | NaN | NaN | NaN | NaN | NaN |
| NaN | NaN | NaN | NaN | NaN | NaN |
| NaN | NaN | NaN | NaN | NaN | NaN |
| NaN | NaN | NaN | NaN | NaN | NaN |
| NaN | NaN | NaN | NaN | NaN | NaN |
| NaN | NaN | NaN | NaN | NaN | NaN |
| NaN | NaN | NaN | NaN | NaN | NaN |
| NaN | NaN | NaN | NaN | NaN | NaN |
| NaN | NaN | NaN | NaN | NaN | NaN |
| NaN | NaN | NaN | NaN | NaN | NaN |
| NaN | NaN | NaN | NaN | NaN | NaN |
| NaN | NaN | NaN | NaN | NaN | NaN |
| NaN | NaN | NaN | NaN | NaN | NaN |
| NaN | NaN | NaN | NaN | NaN | NaN |
| NaN | NaN | NaN | NaN | NaN | NaN |
| NaN | NaN | NaN | NaN | NaN | NaN |

| | | | | | |
|---|---|---|---|---|---|
| NaN | NaN | NaN | NaN | NaN | NaN |
| NaN | NaN | NaN | NaN | NaN | NaN |
| NaN | NaN | NaN | NaN | NaN | NaN |
| NaN | NaN | NaN | NaN | NaN | NaN |
| NaN | NaN | NaN | NaN | NaN | NaN |
| NaN | NaN | NaN | NaN | NaN | NaN |
| NaN | NaN | NaN | NaN | NaN | NaN |
| NaN | NaN | NaN | NaN | NaN | NaN |
| NaN | NaN | NaN | NaN | NaN | NaN |
| NaN | NaN | NaN | NaN | NaN | NaN |
| NaN | NaN | NaN | NaN | NaN | NaN |
| NaN | NaN | NaN | NaN | NaN | NaN |
| NaN | NaN | NaN | NaN | NaN | NaN |
| NaN | NaN | NaN | NaN | NaN | NaN |
| NaN | NaN | NaN | NaN | NaN | NaN |
| NaN | NaN | NaN | NaN | NaN | NaN |
| NaN | NaN | NaN | NaN | NaN | NaN |
| NaN | NaN | NaN | NaN | NaN | NaN |
| NaN | NaN | NaN | NaN | NaN | NaN |
| NaN | NaN | NaN | NaN | NaN | NaN |
| NaN | NaN | NaN | NaN | NaN | NaN |
| NaN | NaN | NaN | NaN | NaN | NaN |
| NaN | NaN | NaN | NaN | NaN | NaN |
| NaN | NaN | NaN | NaN | NaN | NaN |
| NaN | NaN | NaN | NaN | NaN | NaN |
| NaN | NaN | NaN | NaN | NaN | NaN |
| NaN | NaN | NaN | NaN | NaN | NaN |
| NaN | NaN | NaN | NaN | NaN | NaN |
| NaN | NaN | NaN | NaN | NaN | NaN |
| NaN | NaN | NaN | NaN | NaN | NaN |
| NaN | NaN | NaN | NaN | NaN | NaN |
| NaN | NaN | NaN | NaN | NaN | NaN |
| NaN | NaN | NaN | NaN | NaN | NaN |
| NaN | NaN | NaN | NaN | NaN | NaN |
| NaN | NaN | NaN | NaN | NaN | NaN |
| NaN | NaN | NaN | NaN | NaN | NaN |
| NaN | NaN | NaN | NaN | NaN | NaN |
| NaN | NaN | NaN | NaN | NaN | NaN |
| NaN | NaN | NaN | NaN | NaN | NaN |
| NaN | NaN | NaN | NaN | NaN | NaN |
| NaN | NaN | NaN | NaN | NaN | NaN |
| NaN | NaN | NaN | NaN | NaN | NaN |
| NaN | NaN | NaN | NaN | NaN | NaN |
| NaN | NaN | NaN | NaN | NaN | NaN |
| NaN | NaN | NaN | NaN | NaN | NaN |
| NaN | NaN | NaN | NaN | NaN | NaN |
| NaN | NaN | NaN | NaN | NaN | NaN |
| NaN | NaN | NaN | NaN | NaN | NaN |

| | | | | | |
|---|---|---|---|---|---|
| NaN | NaN | NaN | NaN | NaN | NaN |
| NaN | NaN | NaN | NaN | NaN | NaN |
| NaN | NaN | NaN | NaN | NaN | NaN |
| NaN | NaN | NaN | NaN | NaN | NaN |
| NaN | NaN | NaN | NaN | NaN | NaN |
| NaN | NaN | NaN | NaN | NaN | NaN |
| NaN | NaN | NaN | NaN | NaN | NaN |
| NaN | NaN | NaN | NaN | NaN | NaN |
| NaN | NaN | NaN | NaN | NaN | NaN |
| NaN | NaN | NaN | NaN | NaN | NaN |
| NaN | NaN | NaN | NaN | NaN | NaN |
| NaN | NaN | NaN | NaN | NaN | NaN |
| NaN | NaN | NaN | NaN | NaN | NaN |
| NaN | NaN | NaN | NaN | NaN | NaN |
| NaN | NaN | NaN | NaN | NaN | NaN |
| NaN | NaN | NaN | NaN | NaN | NaN |
| NaN | NaN | NaN | NaN | NaN | NaN |
| NaN | NaN | NaN | NaN | NaN | NaN |
| NaN | NaN | NaN | NaN | NaN | NaN |
| NaN | NaN | NaN | NaN | NaN | NaN |
| NaN | NaN | NaN | NaN | NaN | NaN |
| NaN | NaN | NaN | NaN | NaN | NaN |
| NaN | NaN | NaN | NaN | NaN | NaN |
| NaN | NaN | NaN | NaN | NaN | NaN |
| NaN | NaN | NaN | NaN | NaN | NaN |
| NaN | NaN | NaN | NaN | NaN | NaN |
| NaN | NaN | NaN | NaN | NaN | NaN |
| NaN | NaN | NaN | NaN | NaN | NaN |
| NaN | NaN | NaN | NaN | NaN | NaN |
| NaN | NaN | NaN | NaN | NaN | NaN |
| NaN | NaN | NaN | NaN | NaN | NaN |
| NaN | NaN | NaN | NaN | NaN | NaN |
| NaN | NaN | NaN | NaN | NaN | NaN |
| NaN | NaN | NaN | NaN | NaN | NaN |
| NaN | NaN | NaN | NaN | NaN | NaN |
| NaN | NaN | NaN | NaN | NaN | NaN |
| NaN | NaN | NaN | NaN | NaN | NaN |
| NaN | NaN | NaN | NaN | NaN | NaN |
| NaN | NaN | NaN | NaN | NaN | NaN |
| NaN | NaN | NaN | NaN | NaN | NaN |
| NaN | NaN | NaN | NaN | NaN | NaN |
| NaN | NaN | NaN | NaN | NaN | NaN |
| NaN | NaN | NaN | NaN | NaN | NaN |
| NaN | NaN | NaN | NaN | NaN | NaN |
| NaN | NaN | NaN | NaN | NaN | NaN |
| NaN | NaN | NaN | NaN | NaN | NaN |
| NaN | NaN | NaN | NaN | NaN | NaN |
| NaN | NaN | NaN | NaN | NaN | NaN |

| | | | | | |
|---|---|---|---|---|---|
| NaN | NaN | NaN | NaN | NaN | NaN |
| NaN | NaN | NaN | NaN | NaN | NaN |
| NaN | NaN | NaN | NaN | NaN | NaN |
| NaN | NaN | NaN | NaN | NaN | NaN |
| NaN | NaN | NaN | NaN | NaN | NaN |
| NaN | NaN | NaN | NaN | NaN | NaN |
| NaN | NaN | NaN | NaN | NaN | NaN |
| NaN | NaN | NaN | NaN | NaN | NaN |
| NaN | NaN | NaN | NaN | NaN | NaN |
| NaN | NaN | NaN | NaN | NaN | NaN |
| NaN | NaN | NaN | NaN | NaN | NaN |
| NaN | NaN | NaN | NaN | NaN | NaN |
| NaN | NaN | NaN | NaN | NaN | NaN |
| NaN | NaN | NaN | NaN | NaN | NaN |
| NaN | NaN | NaN | NaN | NaN | NaN |
| NaN | NaN | NaN | NaN | NaN | NaN |
| NaN | NaN | NaN | NaN | NaN | NaN |
| NaN | NaN | NaN | NaN | NaN | NaN |
| NaN | NaN | NaN | NaN | NaN | NaN |
| NaN | NaN | NaN | NaN | NaN | NaN |
| NaN | NaN | NaN | NaN | NaN | NaN |
| NaN | NaN | NaN | NaN | NaN | NaN |
| NaN | NaN | NaN | NaN | NaN | NaN |
| NaN | NaN | NaN | NaN | NaN | NaN |
| NaN | NaN | NaN | NaN | NaN | NaN |
| NaN | NaN | NaN | NaN | NaN | NaN |
| NaN | NaN | NaN | NaN | NaN | NaN |
| NaN | NaN | NaN | NaN | NaN | NaN |
| NaN | NaN | NaN | NaN | NaN | NaN |
| NaN | NaN | NaN | NaN | NaN | NaN |
| NaN | NaN | NaN | NaN | NaN | NaN |
| NaN | NaN | NaN | NaN | NaN | NaN |
| NaN | NaN | NaN | NaN | NaN | NaN |
| NaN | NaN | NaN | NaN | NaN | NaN |
| NaN | NaN | NaN | NaN | NaN | NaN |
| NaN | NaN | NaN | NaN | NaN | NaN |
| NaN | NaN | NaN | NaN | NaN | NaN |
| NaN | NaN | NaN | NaN | NaN | NaN |
| NaN | NaN | NaN | NaN | NaN | NaN |
| NaN | NaN | NaN | NaN | NaN | NaN |
| NaN | NaN | NaN | NaN | NaN | NaN |
| NaN | NaN | NaN | NaN | NaN | NaN |
| NaN | NaN | NaN | NaN | NaN | NaN |
| NaN | NaN | NaN | NaN | NaN | NaN |
| NaN | NaN | NaN | NaN | NaN | NaN |
| NaN | NaN | NaN | NaN | NaN | NaN |
| NaN | NaN | NaN | NaN | NaN | NaN |
| NaN | NaN | NaN | NaN | NaN | NaN |

| | | | | | |
|---|---|---|---|---|---|
| NaN | NaN | NaN | NaN | NaN | NaN |
| NaN | NaN | NaN | NaN | NaN | NaN |
| NaN | NaN | NaN | NaN | NaN | NaN |
| NaN | NaN | NaN | NaN | NaN | NaN |
| NaN | NaN | NaN | NaN | NaN | NaN |
| NaN | NaN | NaN | NaN | NaN | NaN |
| NaN | NaN | NaN | NaN | NaN | NaN |
| NaN | NaN | NaN | NaN | NaN | NaN |
| NaN | NaN | NaN | NaN | NaN | NaN |
| NaN | NaN | NaN | NaN | NaN | NaN |
| NaN | NaN | NaN | NaN | NaN | NaN |
| NaN | NaN | NaN | NaN | NaN | NaN |
| NaN | NaN | NaN | NaN | NaN | NaN |
| NaN | NaN | NaN | NaN | NaN | NaN |
| NaN | NaN | NaN | NaN | NaN | NaN |
| NaN | NaN | NaN | NaN | NaN | NaN |
| NaN | NaN | NaN | NaN | NaN | NaN |
| NaN | NaN | NaN | NaN | NaN | NaN |
| NaN | NaN | NaN | NaN | NaN | NaN |
| NaN | NaN | NaN | NaN | NaN | NaN |
| NaN | NaN | NaN | NaN | NaN | NaN |
| NaN | NaN | NaN | NaN | NaN | NaN |
| NaN | NaN | NaN | NaN | NaN | NaN |
| NaN | NaN | NaN | NaN | NaN | NaN |
| NaN | NaN | NaN | NaN | NaN | NaN |
| NaN | NaN | NaN | NaN | NaN | NaN |
| NaN | NaN | NaN | NaN | NaN | NaN |
| NaN | NaN | NaN | NaN | NaN | NaN |
| NaN | NaN | NaN | NaN | NaN | NaN |
| NaN | NaN | NaN | NaN | NaN | NaN |
| NaN | NaN | NaN | NaN | NaN | NaN |
| NaN | NaN | NaN | NaN | NaN | NaN |
| NaN | NaN | NaN | NaN | NaN | NaN |
| NaN | NaN | NaN | NaN | NaN | NaN |
| NaN | NaN | NaN | NaN | NaN | NaN |
| NaN | NaN | NaN | NaN | NaN | NaN |
| NaN | NaN | NaN | NaN | NaN | NaN |
| NaN | NaN | NaN | NaN | NaN | NaN |
| NaN | NaN | NaN | NaN | NaN | NaN |
| NaN | NaN | NaN | NaN | NaN | NaN |
| NaN | NaN | NaN | NaN | NaN | NaN |
| NaN | NaN | NaN | NaN | NaN | NaN |
| NaN | NaN | NaN | NaN | NaN | NaN |
| NaN | NaN | NaN | NaN | NaN | NaN |
| NaN | NaN | NaN | NaN | NaN | NaN |
| NaN | NaN | NaN | NaN | NaN | NaN |
| NaN | NaN | NaN | NaN | NaN | NaN |
| NaN | NaN | NaN | NaN | NaN | NaN |

| | | | | | |
|---|---|---|---|---|---|
| NaN | NaN | NaN | NaN | NaN | NaN |
| NaN | NaN | NaN | NaN | NaN | NaN |
| NaN | NaN | NaN | NaN | NaN | NaN |
| NaN | NaN | NaN | NaN | NaN | NaN |
| NaN | NaN | NaN | NaN | NaN | NaN |
| NaN | NaN | NaN | NaN | NaN | NaN |
| NaN | NaN | NaN | NaN | NaN | NaN |
| NaN | NaN | NaN | NaN | NaN | NaN |
| NaN | NaN | NaN | NaN | NaN | NaN |
| NaN | NaN | NaN | NaN | NaN | NaN |
| NaN | NaN | NaN | NaN | NaN | NaN |
| NaN | NaN | NaN | NaN | NaN | NaN |
| NaN | NaN | NaN | NaN | NaN | NaN |
| NaN | NaN | NaN | NaN | NaN | NaN |
| NaN | NaN | NaN | NaN | NaN | NaN |
| NaN | NaN | NaN | NaN | NaN | NaN |
| NaN | NaN | NaN | NaN | NaN | NaN |
| NaN | NaN | NaN | NaN | NaN | NaN |
| NaN | NaN | NaN | NaN | NaN | NaN |
| NaN | NaN | NaN | NaN | NaN | NaN |
| NaN | NaN | NaN | NaN | NaN | NaN |
| NaN | NaN | NaN | NaN | NaN | NaN |
| NaN | NaN | NaN | NaN | NaN | NaN |
| NaN | NaN | NaN | NaN | NaN | NaN |
| NaN | NaN | NaN | NaN | NaN | NaN |
| NaN | NaN | NaN | NaN | NaN | NaN |
| NaN | NaN | NaN | NaN | NaN | NaN |
| NaN | NaN | NaN | NaN | NaN | NaN |
| NaN | NaN | NaN | NaN | NaN | NaN |
| NaN | NaN | NaN | NaN | NaN | NaN |
| NaN | NaN | NaN | NaN | NaN | NaN |
| NaN | NaN | NaN | NaN | NaN | NaN |
| NaN | NaN | NaN | NaN | NaN | NaN |
| NaN | NaN | NaN | NaN | NaN | NaN |
| NaN | NaN | NaN | NaN | NaN | NaN |
| NaN | NaN | NaN | NaN | NaN | NaN |
| NaN | NaN | NaN | NaN | NaN | NaN |
| NaN | NaN | NaN | NaN | NaN | NaN |
| NaN | NaN | NaN | NaN | NaN | NaN |
| NaN | NaN | NaN | NaN | NaN | NaN |
| NaN | NaN | NaN | NaN | NaN | NaN |
| NaN | NaN | NaN | NaN | NaN | NaN |
| NaN | NaN | NaN | NaN | NaN | NaN |
| NaN | NaN | NaN | NaN | NaN | NaN |
| NaN | NaN | NaN | NaN | NaN | NaN |
| NaN | NaN | NaN | NaN | NaN | NaN |
| NaN | NaN | NaN | NaN | NaN | NaN |
| NaN | NaN | NaN | NaN | NaN | NaN |

| | | | | | |
|---|---|---|---|---|---|
| NaN | NaN | NaN | NaN | NaN | NaN |
| NaN | NaN | NaN | NaN | NaN | NaN |
| NaN | NaN | NaN | NaN | NaN | NaN |
| NaN | NaN | NaN | NaN | NaN | NaN |
| NaN | NaN | NaN | NaN | NaN | NaN |
| NaN | NaN | NaN | NaN | NaN | NaN |
| NaN | NaN | NaN | NaN | NaN | NaN |
| NaN | NaN | NaN | NaN | NaN | NaN |
| NaN | NaN | NaN | NaN | NaN | NaN |
| NaN | NaN | NaN | NaN | NaN | NaN |
| NaN | NaN | NaN | NaN | NaN | NaN |
| NaN | NaN | NaN | NaN | NaN | NaN |
| NaN | NaN | NaN | NaN | NaN | NaN |
| NaN | NaN | NaN | NaN | NaN | NaN |
| NaN | NaN | NaN | NaN | NaN | NaN |
| NaN | NaN | NaN | NaN | NaN | NaN |
| NaN | NaN | NaN | NaN | NaN | NaN |
| NaN | NaN | NaN | NaN | NaN | NaN |
| NaN | NaN | NaN | NaN | NaN | NaN |
| NaN | NaN | NaN | NaN | NaN | NaN |
| NaN | NaN | NaN | NaN | NaN | NaN |
| NaN | NaN | NaN | NaN | NaN | NaN |
| NaN | NaN | NaN | NaN | NaN | NaN |
| NaN | NaN | NaN | NaN | NaN | NaN |
| NaN | NaN | NaN | NaN | NaN | NaN |
| NaN | NaN | NaN | NaN | NaN | NaN |
| NaN | NaN | NaN | NaN | NaN | NaN |
| NaN | NaN | NaN | NaN | NaN | NaN |
| NaN | NaN | NaN | NaN | NaN | NaN |
| NaN | NaN | NaN | NaN | NaN | NaN |
| NaN | NaN | NaN | NaN | NaN | NaN |
| NaN | NaN | NaN | NaN | NaN | NaN |
| NaN | NaN | NaN | NaN | NaN | NaN |
| NaN | NaN | NaN | NaN | NaN | NaN |
| NaN | NaN | NaN | NaN | NaN | NaN |
| NaN | NaN | NaN | NaN | NaN | NaN |
| NaN | NaN | NaN | NaN | NaN | NaN |
| NaN | NaN | NaN | NaN | NaN | NaN |
| NaN | NaN | NaN | NaN | NaN | NaN |
| NaN | NaN | NaN | NaN | NaN | NaN |
| NaN | NaN | NaN | NaN | NaN | NaN |
| NaN | NaN | NaN | NaN | NaN | NaN |
| NaN | NaN | NaN | NaN | NaN | NaN |
| NaN | NaN | NaN | NaN | NaN | NaN |
| NaN | NaN | NaN | NaN | NaN | NaN |
| NaN | NaN | NaN | NaN | NaN | NaN |
| NaN | NaN | NaN | NaN | NaN | NaN |
| NaN | NaN | NaN | NaN | NaN | NaN |

| | | | | | |
|---|---|---|---|---|---|
| NaN | NaN | NaN | NaN | NaN | NaN |
| NaN | NaN | NaN | NaN | NaN | NaN |
| NaN | NaN | NaN | NaN | NaN | NaN |
| NaN | NaN | NaN | NaN | NaN | NaN |
| NaN | NaN | NaN | NaN | NaN | NaN |
| NaN | NaN | NaN | NaN | NaN | NaN |
| NaN | NaN | NaN | NaN | NaN | NaN |
| NaN | NaN | NaN | NaN | NaN | NaN |
| NaN | NaN | NaN | NaN | NaN | NaN |
| NaN | NaN | NaN | NaN | NaN | NaN |
| NaN | NaN | NaN | NaN | NaN | NaN |
| NaN | NaN | NaN | NaN | NaN | NaN |
| NaN | NaN | NaN | NaN | NaN | NaN |
| NaN | NaN | NaN | NaN | NaN | NaN |
| NaN | NaN | NaN | NaN | NaN | NaN |
| NaN | NaN | NaN | NaN | NaN | NaN |
| NaN | NaN | NaN | NaN | NaN | NaN |
| NaN | NaN | NaN | NaN | NaN | NaN |
| NaN | NaN | NaN | NaN | NaN | NaN |
| NaN | NaN | NaN | NaN | NaN | NaN |
| NaN | NaN | NaN | NaN | NaN | NaN |
| NaN | NaN | NaN | NaN | NaN | NaN |
| NaN | NaN | NaN | NaN | NaN | NaN |
| NaN | NaN | NaN | NaN | NaN | NaN |
| NaN | NaN | NaN | NaN | NaN | NaN |
| NaN | NaN | NaN | NaN | NaN | NaN |
| NaN | NaN | NaN | NaN | NaN | NaN |
| NaN | NaN | NaN | NaN | NaN | NaN |
| NaN | NaN | NaN | NaN | NaN | NaN |
| NaN | NaN | NaN | NaN | NaN | NaN |
| NaN | NaN | NaN | NaN | NaN | NaN |
| NaN | NaN | NaN | NaN | NaN | NaN |
| NaN | NaN | NaN | NaN | NaN | NaN |
| NaN | NaN | NaN | NaN | NaN | NaN |
| NaN | NaN | NaN | NaN | NaN | NaN |
| NaN | NaN | NaN | NaN | NaN | NaN |
| NaN | NaN | NaN | NaN | NaN | NaN |
| NaN | NaN | NaN | NaN | NaN | NaN |
| NaN | NaN | NaN | NaN | NaN | NaN |
| NaN | NaN | NaN | NaN | NaN | NaN |
| NaN | NaN | NaN | NaN | NaN | NaN |
| NaN | NaN | NaN | NaN | NaN | NaN |
| NaN | NaN | NaN | NaN | NaN | NaN |
| NaN | NaN | NaN | NaN | NaN | NaN |
| NaN | NaN | NaN | NaN | NaN | NaN |
| NaN | NaN | NaN | NaN | NaN | NaN |
| NaN | NaN | NaN | NaN | NaN | NaN |
| NaN | NaN | NaN | NaN | NaN | NaN |

| | | | | | |
|---|---|---|---|---|---|
| NaN | NaN | NaN | NaN | NaN | NaN |
| NaN | NaN | NaN | NaN | NaN | NaN |
| NaN | NaN | NaN | NaN | NaN | NaN |
| NaN | NaN | NaN | NaN | NaN | NaN |
| NaN | NaN | NaN | NaN | NaN | NaN |
| NaN | NaN | NaN | NaN | NaN | NaN |
| NaN | NaN | NaN | NaN | NaN | NaN |
| NaN | NaN | NaN | NaN | NaN | NaN |
| NaN | NaN | NaN | NaN | NaN | NaN |
| NaN | NaN | NaN | NaN | NaN | NaN |
| NaN | NaN | NaN | NaN | NaN | NaN |
| NaN | NaN | NaN | NaN | NaN | NaN |
| NaN | NaN | NaN | NaN | NaN | NaN |
| NaN | NaN | NaN | NaN | NaN | NaN |
| NaN | NaN | NaN | NaN | NaN | NaN |
| NaN | NaN | NaN | NaN | NaN | NaN |
| NaN | NaN | NaN | NaN | NaN | NaN |
| NaN | NaN | NaN | NaN | NaN | NaN |
| NaN | NaN | NaN | NaN | NaN | NaN |
| NaN | NaN | NaN | NaN | NaN | NaN |
| NaN | NaN | NaN | NaN | NaN | NaN |
| NaN | NaN | NaN | NaN | NaN | NaN |
| NaN | NaN | NaN | NaN | NaN | NaN |
| NaN | NaN | NaN | NaN | NaN | NaN |
| NaN | NaN | NaN | NaN | NaN | NaN |
| NaN | NaN | NaN | NaN | NaN | NaN |
| NaN | NaN | NaN | NaN | NaN | NaN |
| NaN | NaN | NaN | NaN | NaN | NaN |
| NaN | NaN | NaN | NaN | NaN | NaN |
| NaN | NaN | NaN | NaN | NaN | NaN |
| NaN | NaN | NaN | NaN | NaN | NaN |
| NaN | NaN | NaN | NaN | NaN | NaN |
| NaN | NaN | NaN | NaN | NaN | NaN |
| NaN | NaN | NaN | NaN | NaN | NaN |
| NaN | NaN | NaN | NaN | NaN | NaN |
| NaN | NaN | NaN | NaN | NaN | NaN |
| NaN | NaN | NaN | NaN | NaN | NaN |
| NaN | NaN | NaN | NaN | NaN | NaN |
| NaN | NaN | NaN | NaN | NaN | NaN |
| NaN | NaN | NaN | NaN | NaN | NaN |
| NaN | NaN | NaN | NaN | NaN | NaN |
| NaN | NaN | NaN | NaN | NaN | NaN |
| NaN | NaN | NaN | NaN | NaN | NaN |
| NaN | NaN | NaN | NaN | NaN | NaN |
| NaN | NaN | NaN | NaN | NaN | NaN |
| NaN | NaN | NaN | NaN | NaN | NaN |
| NaN | NaN | NaN | NaN | NaN | NaN |
| NaN | NaN | NaN | NaN | NaN | NaN |

| | | | | | |
|---|---|---|---|---|---|
| NaN | NaN | NaN | NaN | NaN | NaN |
| NaN | NaN | NaN | NaN | NaN | NaN |
| NaN | NaN | NaN | NaN | NaN | NaN |
| NaN | NaN | NaN | NaN | NaN | NaN |
| NaN | NaN | NaN | NaN | NaN | NaN |
| NaN | NaN | NaN | NaN | NaN | NaN |
| NaN | NaN | NaN | NaN | NaN | NaN |
| NaN | NaN | NaN | NaN | NaN | NaN |
| NaN | NaN | NaN | NaN | NaN | NaN |
| NaN | NaN | NaN | NaN | NaN | NaN |
| NaN | NaN | NaN | NaN | NaN | NaN |
| NaN | NaN | NaN | NaN | NaN | NaN |
| NaN | NaN | NaN | NaN | NaN | NaN |
| NaN | NaN | NaN | NaN | NaN | NaN |
| NaN | NaN | NaN | NaN | NaN | NaN |
| NaN | NaN | NaN | NaN | NaN | NaN |
| NaN | NaN | NaN | NaN | NaN | NaN |
| NaN | NaN | NaN | NaN | NaN | NaN |
| NaN | NaN | NaN | NaN | NaN | NaN |
| NaN | NaN | NaN | NaN | NaN | NaN |
| NaN | NaN | NaN | NaN | NaN | NaN |
| NaN | NaN | NaN | NaN | NaN | NaN |
| NaN | NaN | NaN | NaN | NaN | NaN |
| NaN | NaN | NaN | NaN | NaN | NaN |
| NaN | NaN | NaN | NaN | NaN | NaN |
| NaN | NaN | NaN | NaN | NaN | NaN |
| NaN | NaN | NaN | NaN | NaN | NaN |
| NaN | NaN | NaN | NaN | NaN | NaN |
| NaN | NaN | NaN | NaN | NaN | NaN |
| NaN | NaN | NaN | NaN | NaN | NaN |
| NaN | NaN | NaN | NaN | NaN | NaN |
| NaN | NaN | NaN | NaN | NaN | NaN |
| NaN | NaN | NaN | NaN | NaN | NaN |
| NaN | NaN | NaN | NaN | NaN | NaN |
| NaN | NaN | NaN | NaN | NaN | NaN |
| NaN | NaN | NaN | NaN | NaN | NaN |
| NaN | NaN | NaN | NaN | NaN | NaN |
| NaN | NaN | NaN | NaN | NaN | NaN |
| NaN | NaN | NaN | NaN | NaN | NaN |
| NaN | NaN | NaN | NaN | NaN | NaN |

pH LOC BKG pH LYSO

| | |
|---|---|
| 4.062486 | 4.101122 |
| 4.526987 | 4.552689 |
| 3.338991 | 3.397774 |
| 4.059285 | 4.09801 |
| 4.423415 | 4.452001 |
| 4.428703 | 4.457142 |
| 4.167046 | 4.202771 |
| 4.585305 | 4.609384 |
| 4.757845 | 4.777118 |
| 4.423578 | 4.452159 |
| 4.248006 | 4.281476 |
| 4.518466 | 4.544405 |
| 4.538048 | 4.563443 |
| 4.315149 | 4.34675 |
| 4.795847 | 4.814062 |
| 4.516379 | 4.542376 |
| 4.604526 | 4.628069 |

| | |
|---|---|
| 4.546213 | 4.57138 |
| 4.474423 | 4.501589 |
| 3.955227 | 3.996851 |
| 4.564958 | 4.589603 |
| 4.413644 | 4.442503 |
| 4.017903 | 4.057781 |
| 4.554201 | 4.579145 |
| 4.450748 | 4.478573 |
| 4.099257 | 4.136869 |
| 4.289034 | 4.321362 |
| 4.050699 | 4.089663 |
| 4.456287 | 4.483958 |
| 4.507488 | 4.533733 |
| 4.228208 | 4.26223 |
| 4.415126 | 4.443943 |
| 4.514654 | 4.5407 |
| 4.281358 | 4.3139 |
| 4.189533 | 4.224631 |
| 4.109779 | 4.147098 |
| 4.004005 | 4.04427 |
| 3.992639 | 4.033221 |
| 4.004433 | 4.044686 |
| 4.580999 | 4.605197 |
| 4.238627 | 4.272358 |
| 4.308975 | 4.340748 |
| 4.456856 | 4.484511 |
| 4.366557 | 4.396726 |
| 4.528843 | 4.554494 |
| 3.975403 | 4.016464 |
| 4.404833 | 4.433936 |
| 4.401245 | 4.430448 |
| 4.629887 | 4.652724 |
| 4.372795 | 4.40279 |
| 4.044573 | 4.083708 |
| 4.553005 | 4.577983 |
| 4.548213 | 4.573324 |
| 3.960994 | 4.002456 |
| 4.484815 | 4.511692 |
| 4.104413 | 4.141882 |
| 4.029473 | 4.069029 |
| 4.217803 | 4.252115 |
| 4.651729 | 4.673958 |
| 4.511591 | 4.537722 |
| 4.611651 | 4.634996 |
| 4.63576 | 4.658434 |
| 3.934894 | 3.977083 |
| 4.164479 | 4.200275 |
| 4.670269 | 4.691981 |

| | |
|---|---|
| 4.004776 | 4.045019 |
| 4.6702 | 4.691914 |
| 4.07942 | 4.117585 |
| 4.798769 | 4.816903 |
| 4.466913 | 4.494288 |
| 4.355966 | 4.386431 |
| 3.953102 | 3.994784 |
| 4.121702 | 4.158689 |
| 4.453776 | 4.481516 |
